# Instantaneous phase of rhythmic behaviour under volitional control

**DOI:** 10.1101/2023.11.01.564135

**Authors:** Leonardo Lancia

## Abstract

The phase of a signal representing a cyclic behavioural pattern provides valuable information for understanding the mechanisms driving the observed behaviour. Methods usually adopted to estimate the phase, which are based on projecting the signal onto the complex plane, have strict requirements on its frequency content, which limits their application. To overcome these limitations, input signals can be processed using band-pass filters or decomposition techniques. In this paper, we briefly review these approaches and propose a new one. Our approach is based on the principles of Empirical Mode Decomposition (EMD), but unlike EMD, it does not aim to decompose the input signal. This avoids the many problems that can occur when extracting a signal’s components one by one. The proposed approach estimates the phase of experimental signals that have one main oscillatory component modulated by slower activity and perturbed by weak, sparse, or random activity at faster time scales. We illustrate how our approach works by estimating the phase dynamics of synthetic signals and real-world signals representing knee angles during flexion/extension activity, heel height during gait, and the activity of different organs involved in speech production.

## 1. Introduction

### 1.1. The phase of goal-oriented behaviour

Many goal-oriented behaviours, such as walking, chewing, and speaking, display cyclic patterns. Often the observed cycles are nested within each other, forming hierarchies of oscillatory activity that characterise the behaviour of potentially many observable variables. In studies focusing on the organisational principles underlying these rhythmic patterns of activity, the notion of phase, representing the position in the cycle at each instant, plays a central role [1, 2]. Measuring the relations between events in terms of phase lags permits measuring durations in terms that are meaningful to the functioning of the systems under study, because a cycle of activity is a meaningful unit of description of an oscillatory behaviour [3, 4]. At the same time, the phase of a signal is an abstract quantity whose meaning does not depend on the material implementation of the cycles of activity (i.e. the phase should have the same meaning regardless the shape of the cycles). This is therefore an ideal quantity to characterize the relations between heterogeneous signals and systems.

Unfortunately, the implementation of the notion of phase is not straightforward for general time series with arbitrary cycle shapes [5]. The usual approach consists in projecting the behaviour of an observed real-valued signal *x* on that of a signal drawing circles on a plane via the application of the Hilbert transform [6], which approximates its quadrature signal (*y* = ℋ(*x*)). The paired values of *x* and *y* constitute the complex-valued Analytic Signal and represent the coordinates of a point that traces a complete circle around the origin of the axes each time that the cyclic pattern of activity in the observed signal repeats itself. The phase of such a signal can be obtained as 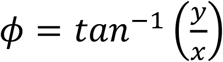.

Principled applications of this definition of phase (henceforth referred to as the Hilbert phase) are limited to time series that can be approximated by a sinusoid with amplitude varying much slower than its phase [5, 6]. Failure to meet this requirement results in incorrect phase estimates, as exemplified in Figure 1. Alternative strategies consist in adopting definitions of phase that are specific to the signal under study (e.g.: 7, 8) and that often focus on a few notable values characterising each cycle (e.g.: velocity peaks, zero-crossing points, points at which thresholds are crossed, etc., e.g.: [9]).

**Figure 1.**
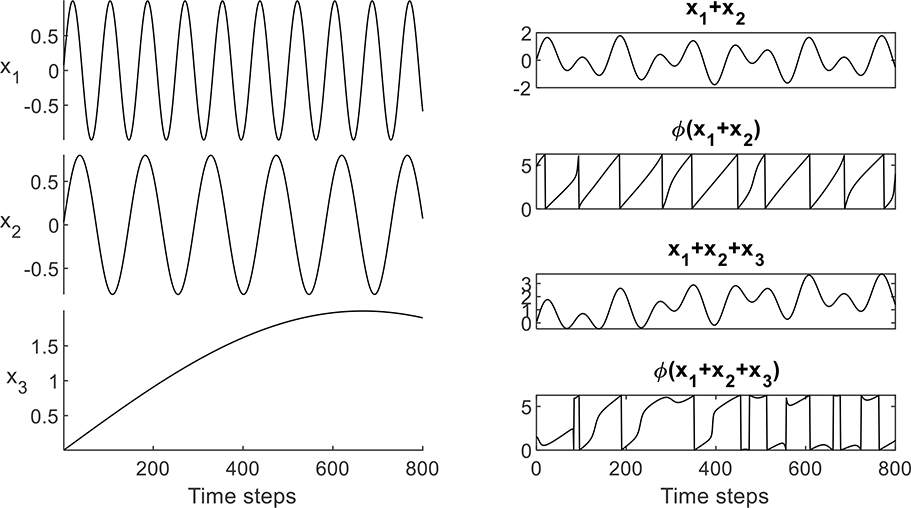
Issues in the application of Hilbert transform. Left column panels: sinusoidal signals composing the time series displayed in the right column. Right column panels: The sum of the first and second panel from the top in the left column is displayed in the topmost panel. Each extremum is surrounded by two zero crossing points but amplitude changes relatively rapidly with respect to phase. Consequently, the dynamics of the signal’s Hilbert phase (displayed in the second panel from top) feature a degree of variability not observed in the (first component of) the input signal. The sum of all time series from the left column is displayed in the third panel from top and its Hilbert phase in the bottommost panel. In this case, the phase does not increase from 0 to 2π in each cycle and discontinuities are observed.

The issues related to phase estimation are greatly reduced when this is conducted on signals submitted to band-pass filters or to decomposition techniques aimed at representing a signal via a combination of oscillatory patterns, because both methods return oscillations with slowly varying amplitude and centred on zero (cf. [5, 10]). For example, in [5] it is proposed to submit the observed signal to Empirical Mode Decomposition (EMD, [11]). This operation produces a small number of oscillatory signals whose amplitude can be further processed so that in each cycle the curves oscillate between -1 and 1 and the quadrature signal can easily be computed. However, approaches based on band-pass filters or on decomposition schemes suffer from two kinds of problem. On the one hand, the selection of the filtering parameters, and the lack of stability of the decomposition algorithms, complicate the identification of the cycles of interest in the signal under study. On the other hand, the application of band-pass filters or of decomposition methods may wash out properties of phase dynamics that make the cycle shape deviate from that of a sinusoid.

### 1.2. Analysing the phase of signals from goal-oriented behaviours

The work presented in the current paper is based on the observation that we can often identify a goal-oriented behaviour through the pattern of variation of one observable quantity or of a quantity that can be obtained by combining different observables. In experiments eliciting rhythmic behaviour; this usually results in signals that are dominated by one oscillatory pattern permitting to characterize the execution of the task under study.

Consider, for example, running or walking. We can characterize the cycles of activity by measuring the height of the heel from the ground. In speech production, we can compute from the observed acoustic signal a sonority coefficient, which oscillates at the rate of syllables production [12] or a spectral modulation coefficient, which oscillates at the rate of speech sounds’ production [13]. Although such signals often tend to display one main peak per cycle, their shape may deviate from that of a sinusoid (see examples in Sections 3.3 and 3.6 below) and they may contain events that are not the expression of the main oscillatory pattern (see examples in Sections 3.4 below). For example, in cases of complex rhythmic patterns involving several interconnected variables and representing the production of sequences of different actions (e.g. speech production, typing, dancing, etc.), we may observe disturbances in the behaviour of each variable, due to mutual dependencies with the other variables. Hierarchically organised rhythmic patterns [14] and general non-stationarity of behaviour may produce slow changes that modulate the activity of the dominant oscillation [15, 16]. Finally, measurement noise may corrupt the observed signal. Still, if we exclude the effect of random noise or of weak and potentially sparse components, a signal collected during the execution of a controlled task can often be modelled as due to the activity of the oscillatory component of interest, potentially modulated by slower oscillatory patterns.

These considerations motivate the approach proposed in the present paper, in which, like in [5], we centre the observed cycles of activity and demodulate them. However, we do not need to submit it to a full decomposition or to narrow band-pass filters with predetermined coefficients because, once the effect of weak and potentially sparse components is excluded, the oscillatory pattern of interest is expected to be the fastest oscillation present in the signal.

The remainder of the paper is structured as follows: we will first summarise the approach proposed in [5] to overcome the issues related to the construction of the Analytic Signal. After discussing the limitations of this approach, we will illustrate the features of our proposed approach. We also illustrate an adaptive filtering approach based on [17] and aiming to remove high frequency random components often present in the observed signal due to measurement noise. In the second part of the paper, we will use several synthetic and real-life example signals to illustrate the functioning of our algorithm. In the discussion section, we will build on the results observed to comment on the features of the proposed methodology. Moreover, we will discuss the implications of our findings for decomposition procedures based on variants of EMD informed by knowledge of the signal’s components. The detailed description of the computations performed in this work is included in the Appendix that, together with the analysed data and the code (Matlab and Python) allowing for computation of the phase, are provided as supplementary material (code also available at https://github.com/LeonardoLancia/Get_Phase_Tools).

## 2. Methods

### 2.1. Huang et al., proposal

#### 2.1.1. Empirical Mode Decomposition

The first step of the approach proposed in [5] is to submit the input signal x[n] (with *n* ∈ {1, …, *N*}) to Empirical Mode Decomposition (EMD). This is an iterative procedure in which the input signal submitted to the *k*^*th*^ iteration is centred on its time-varying mean. The signal obtained (*d*_*k*_*[n]*) is referred to as an Intrinsic Mode Function (IMF) and is considered to represent a component of the input signal. The residual signal *r*_*k*_, obtained by subtracting *d*_*k*_ from the input signal, is used as the input for the next iteration (therefore *r*[*n*]_*k*_ = *d*_*k*−1_[*n*] − *d*_*k*_[*n*]).

The implementation of the centring operation is based on a further iterative process termed Sifting (henceforth referred to as *S*_*q*_(·), with *q* indicating the maximum allowed number of iterations). At each iteration of sifting, two envelopes (*e*^+^and *e*^−^) are obtained by interpolating separately via cubic splines the signal’s local minima and maxima; then the mean of the two envelopes is subtracted from the input, and the output of each iteration is used as an input to the following one (see Figure 2). The process is stopped before the maximum allowed number of iterations has been reached when the signal’s local maxima are all positive and its local minima are all negative, or if other more or less stringent criteria are met.

**Figure 2.**
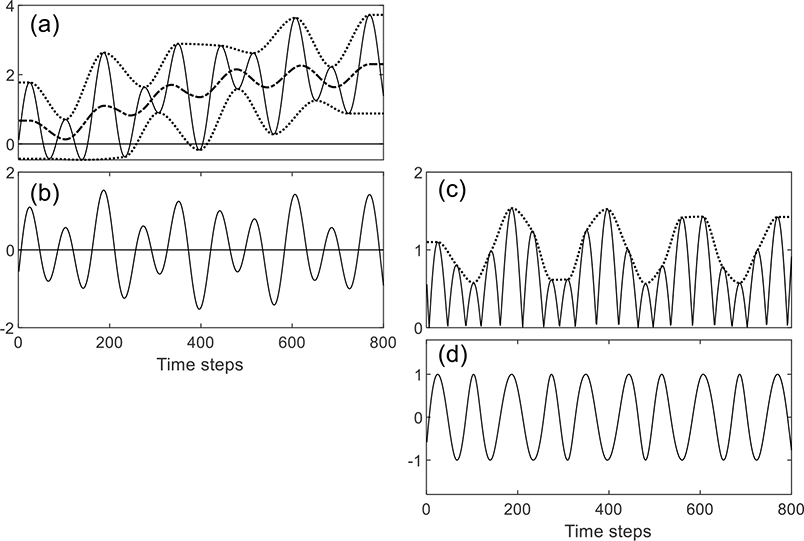
Centring and demodulation operations. Panel (a): Input signal (continuous line), minimum and maximum envelopes (dotted lines) and mean envelope (dashed line) operator. Panel (b): Output of one centring iteration. Panel (c): absolute value of the centred signal displayed in panel (b), and its maximum envelope. Panel (d): signal obtained by dividing the centred signal in panel (b) by the envelope in panel (c).

#### 2.1.2. Demodulation and phase estimation

The IMF obtained from EMD that matches the frequency of the oscillation of interest is submitted to the following demodulation operation (henceforth denoted as *D*(·)). Firstly, the local maxima of the signal’s absolute values are identified and submitted to cubic Interpolation to produce an energy envelope *A*[*n*]. Secondly, the signal values are divided by the corresponding energy envelope values 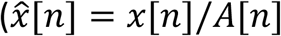, see right column panels in Figure 2). The procedure is iterated, taking as input at each iteration the output of the previous iteration, until no extremum has an absolute value higher than 1+*ε* (where *ε* represents a tolerance parameter) or until a generally small maximum number of iterations has been reached.

The obtained signal can be submitted to the Hilbert transform to obtain a quadrature signal and compute the phase. Hilbert phase, however, may show some bias in the presence of persistent modulation of the signal frequency [18]. In such cases, (and at the expense of numerical robustness) the quadrature signal can also be obtained as 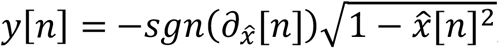, where 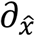 indicates the estimated derivative of the demodulated signal. We will refer to this approach as the Enhanced Hilbert-Huang transform (EHHT, HHT being commonly used to denote the same approach without demodulation). The functioning of this approach is illustrated in Figure 3, where it is applied to the signal resulting by summing the sinusoids in the left panel of Figure 1.

**Figure 3.**
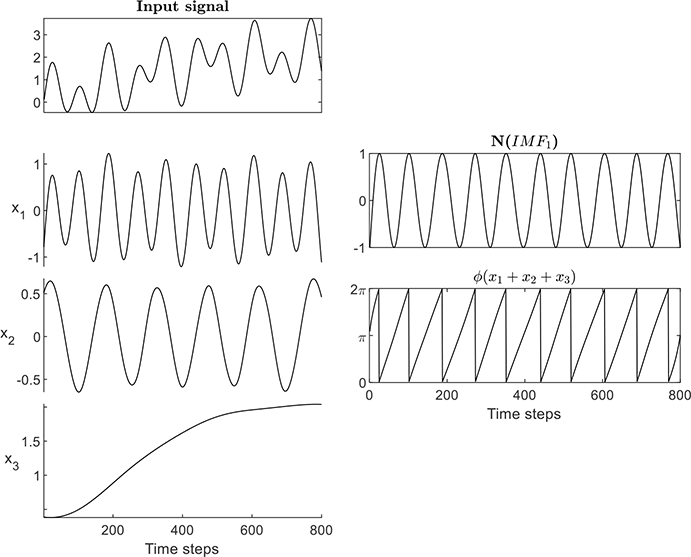
EHHT analysis of the signal resulted by summing the signals in the left panels of Figure 1, left column: observed signal (topmost panel) and IMFs from EMD (remaining panels). Right column: 1^st^ IMF after demodulation (topmost panel) and phase from the application of the direct quadrature method (bottommost panel).

#### 2.1.3. Issues with the EHHT

Being based on extrema detection, EMD is sensitive to the presence of noise and intermittent behaviour, observed when an underlying oscillatory component is not constantly active. Furthermore, EMD biases the analysis to account for variability through changes in the amplitude of the obtained analytic signal. This choice is not motivated by theoretical considerations, but by the limitations of the Hilbert transform approach, which requires a signal with a very slowly changing amplitude. In EMD, such a signal is obtained by iterating the sifting operation, which reshapes its input again and again and gradually makes it similar to a sinusoid ([19, 20]). Consequently, local accelerations and deceleration patterns can be gradually smoothed out.

Figure 4 illustrates how the phase estimation procedure is affected by the addition of a weak, fast-frequency component with intermittent behaviour to the signal analysed in Figure 3. The input signal (displayed in the topmost panels of both the leftmost and middle columns) has been obtained by scaling and adding the signal in the second panel from the top of the leftmost column to the signals displayed in the remaining panels of the same column. Although the intermittent component is barely visible in the input signal at the magnification level adopted (the scaling coefficient being 0.2), its impact on the decomposition procedure is dramatic. Due to its intermittent nature, the additional component characterizes part of the first IMF. That leaves the space for portions of the second component to appear in the remaining parts of that same IMF (second panel from the top of the middle column). Other portions of the second component however characterize the second IMF. Consequently, the true phase of the second component of the input signal, which is the fastest visible component (and which is displayed in the topmost panel of the rightmost column), is captured by different IMFs during different time intervals (as displayed by the remaining panels of the same column).

**Figure 4.**
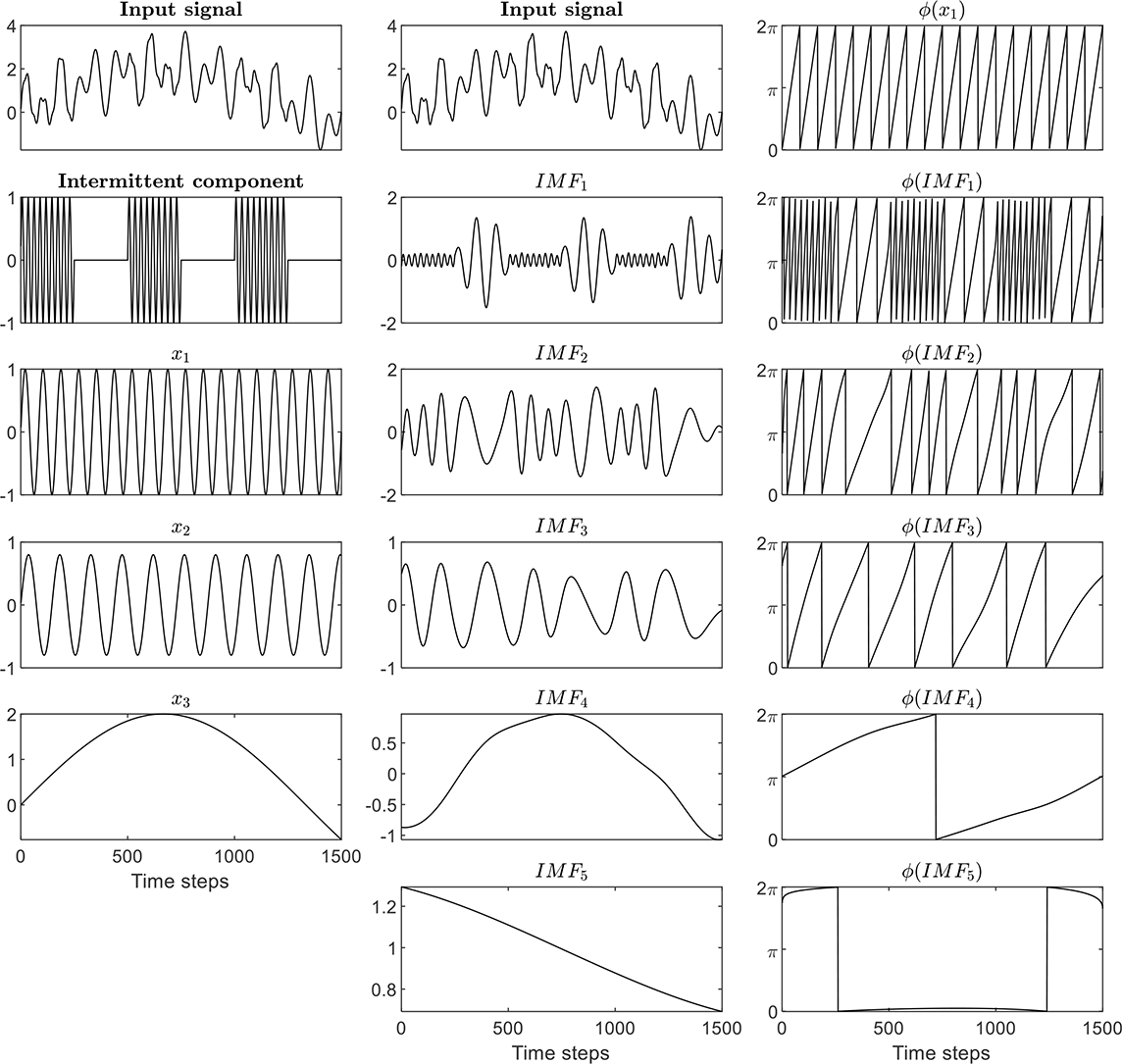
EMD of a longer stretch of the signal in Figure 3 (displayed in the topmost panels of the first and second columns) added to an intermittent oscillation (displayed in the second panel from the top of the leftmost column) multiplied by 0.2. The remaining panels of the leftmost column display the other components of the signal. The central column displays the same input signal in the topmost panel and the IMFs obtained by applying EMD in the remaining panels. The topmost panel of the third column displays the true phase of the second component of the input signal, while the following panels of the same column display the phase values extracted from each IMF after submitting it to the demodulation procedure proposed in [5].

A further limitation of the EHHT is related to the implementation of the demodulation schema based on cubic spline interpolation. According to [5], due to the inherent smoothness of the adopted basis functions, rapid changes of amplitude may not be captured by the obtained amplitude envelope. This produces negative local maxima and positive local minima whose presence may introduce phase jumps and/or intervals characterised by negative frequencies (see Figure 5).

**Figure 5.**
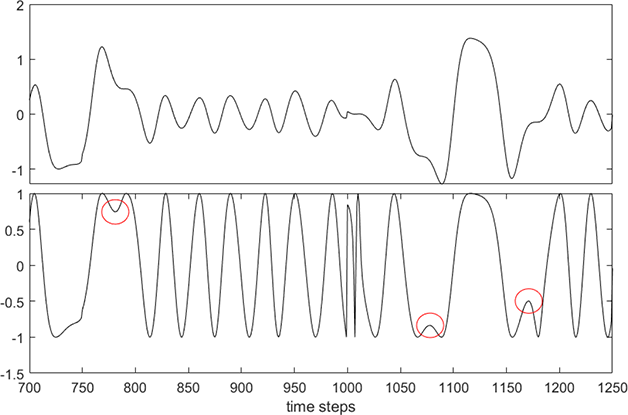
Local extrema escaping the demodulation introduced in [5]. Top panel: The original signal. Bottom panel: The signal after demodulation. Circles indicate positive local minima and negative local maxima.

### 2.2. Our proposal

The proposed approach is based on masked sifting [21; 22], a variant of the sifting process that leverages the knowledge of the frequency of oscillation of the oscillatory component of interest to factor out the effects of slower components. In order to remove the effects of sparse or weak fast components, the frequency value used to drive the sifting is obtained from the analysis of a centred version of the input signal, after application of a modified demodulation procedure that guarantees the absence of negative local maxima and positive local minima. The signal obtained by the application of masked sifting is then submitted to the modified demodulation procedure and used to construct an analytic signal from which the phase can be computed.

#### 2.2.1. First part 1: centring

The first part of the algorithm is aimed at estimating the dominant oscillatory frequency of the input signal *x*[*n*] (whit *n* ∈ {1, …, *N*}). The first step consists in centring *x*[*n*], which is obtained through one iteration of the sifting process 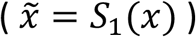. This early stopping is expected to reduce the effect of weak high-frequency component, which are less likely to be captured at the first iteration.

#### 2.2.2. Improved demodulation

In the EHHT, some extrema escape demodulation because they are extracted from the signal’s absolute values. Indeed, the absolute value of a signal containing positive local minima or negative local maxima will not display any local maximum at these locations (see Figure 6 caption and panels in the leftmost column). The envelope obtained does not reflect the presence of these local extrema, which therefore are not affected when the input signal is divided by the envelope. This issue can be easily avoided by adopting a refined demodulation procedure (henceforth referred to as *𝓍*(·)) in which we select local minima and local maxima separately in the input signal, we compute their absolute values, and we obtain the signal’s envelope via interpolation of these values (see the right column panels of Figure 6).

**Figure 6.**
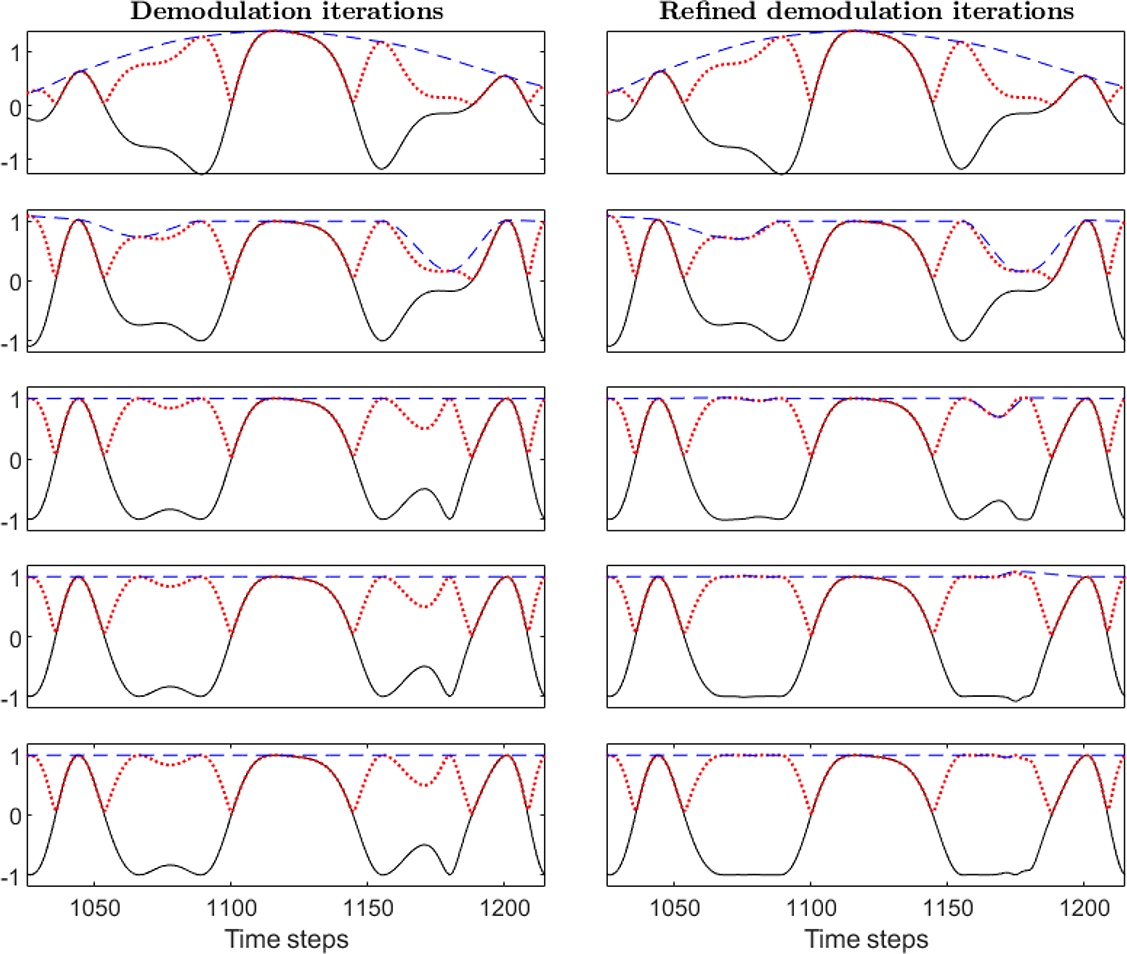
Five iterations of the demodulation based on the local maxima of the absolute values of the input signal (left column panels) and based on the local extrema of the input signal (right column panels). Each panel displays the input signal (continuous black line), the signal of its absolute values (dotted red line), and the envelope obtained by interpolating the extrema identified (dashed blue line). Signals from the first iteration are displayed in the topmost panels. With both approaches, one local maximum smaller than 0 emerges at the second iteration around time step 1070, and another at the third iteration around time step 1180. These extrema do not affect the envelopes displayed in the left column panels because the points do not correspond to local maxima of the absolute values signal. However, the peaks are captured by the envelopes in the right column panels, which are based on the locations of all the extrema of the input signal.

#### 2.2.3. First part 3: Frequency estimation

We first compute *ϕ*[*n*] through application of the Hilbert transform 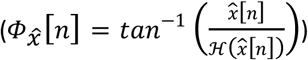. The use of the Hilbert transform to obtain the quadrature signal required for the computation of 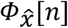 is motivated by its numerical stability and by the fact that the bias characterising the phase obtained in this way does not affect average frequencies computed over sufficiently long time windows. In order to increase the robustness of frequency computation, the instantaneous phase is unwrapped so as to increase monotonically over time and 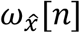 is estimated via the application of a Savitzky-Golay differentiator 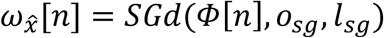, order: *o*_*sg*_ = 5, and length: *l*_*sg*_ = 16 time steps).

**Tableau 1.**
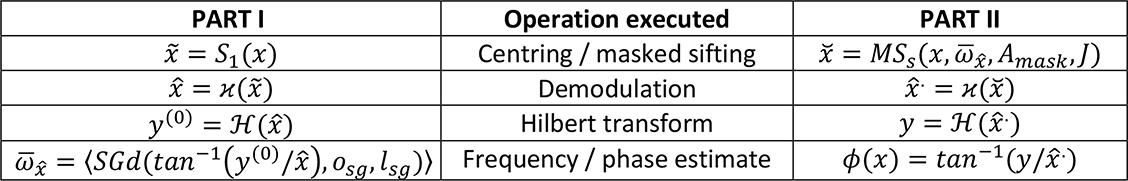
Processing steps of the two parts of our proposed approach. Leftmost and rightmost columns display the formulas implementing the operations described in the centre column. The slash separates operations applied at corresponding steps in the two parts. In the second part, the centred signal is obtained via masked sifting based on the average instantaneous frequency obtained as output of the first part of the algorithm.

#### 2.2.4. Second part I: masked sifting

In the second part of our algorithm, we use the frequency 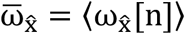 (with hooks representing averaging) to process the signal *x*[*n*] via masked sifting [21; 22]. Masked-sifting (henceforth *MS*_*s*_(*r, ω*_*mask*_, *A*_*mask*_, *J*)) takes as input a signal *x*[*n*] containing *N* values collected with sampling period ∆*t*, a frequency value *ω*_*mask*_, an amplitude value *A*_*mask*_ and a positive integer *J*. The index *s* represents the maximum allowed number of iterations in the sifting process. First, *J* phase-shifted copies of a masking sinusoidal signal are created:

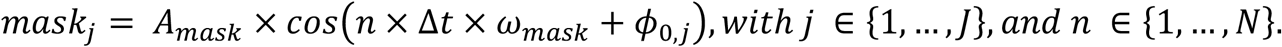

The obtained *J* signals will have the same amplitude *A*_*mask*_ and frequency *ω*_*mask*_ but different initial phases *Φ*_*0*,*j*_. The input signal is independently summed with each *mask*_*j*_ and submitted to the sifting operator. Partial IMFs are obtained by subtracting the masking signals from the results of the sifting operations (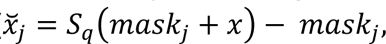 ∀*j* ∈ {1, …, *N*}). Finally, the partial IMFs are averaged to produce an independent mode function 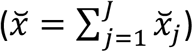.

The presence of the masking signals prevents the sifting operation from capturing oscillations with frequencies smaller than *ω*_*mask*_ and thus effectively removes the effects of low-frequency components. The amplitude of the masking signal *A*_*mask*_ is set at a fixed proportion of the signal’s standard deviation (*A*_*mask*_ = *C*_*a*_*σ*(*x*), with *σ*(*x*) indicating the signal’s standard deviation).

#### 2.2.5. Second part II: Demodulation, quadrature signal and phase estimation

The oscillatory pattern obtained by application of the masked sifting operator to the input signal 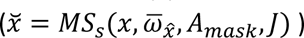, is submitted to refined demodulation 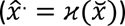. The approach to be adopted for the computation of the quadrature signal should probably depend on the application. Considerations of numerical robustness would suggest the use of the Hilbert transform (but see [23] for robustness improvements). However, rapid changes of the signal’s instantaneous frequency may introduce local biases in phase estimates [18]. If these changes are due to the presence of non-linearities in the underlying systems, their effects on the measured phase are reduced (see simulations in [5]). Moreover, if the aim of the analysis is that of measuring the variability of phase lags between signals, this issue may be less critical, because the effect of nonlinearities on the signal’s shape will be consistently captured across cycles by Hilbert transform phase.

### 2.3. Processing noisy signals via masked EMD filtering

The approach described so far deals with smooth signals, such as those obtained by low-pass filtering the data. In order to deal with signals containing high-frequency noise, we propose an adaptive filtering step based on the one introduced by [17]. Originally, the approach is based on the application of EMD to the input signal and on the identification of the modes dominated by random behaviour. The denoised signal is then obtained by summing the remaining IMFs. In our implementation, we substitute EMD with masked EMD (a variant of EMD that uses masked sifting instead of classical sifting [24, 25]), which we modified to benefit from the frequency estimation procedure proposed in sections 2.2.1-2.2.3.

#### 2.3.1. Masked EMD

At iteration *k* of the masked EMD procedure adopted in our work, the input signal (*x*), or the residual of the previous iteration (*r*_*k*−1_), is submitted to centring (*S*_*q*_(·)) and the results is amplitude normalized (through the application of *𝓍*(·)). A frequency value is computed based on the application of Hilbert transform and used to drive masked sifting (*MS*_*s*_(·)), as illustrated in section 2.2.4. The output of masked sifting at the *k*^*th*^ iteration is stored as an IMF (*d*_*k*_ = *MS*_*s*_(*r*_*k*−1_)) and the residual (*r*_*k*_ = *r*_*k*−1_ − *d*_*k*_) is submitted as input to the next iteration. The process is stopped when the residual obtained does not display local extrema or when a predetermined number of modes has been extracted.

The amplitude of the masking signal *A*_*mask*_ is set at a fixed proportion of the first mode’s standard deviation (*A*_*mask*_ = *C*_*a*_*σ*(*d*_1_)), or, at the first iteration, of the centred input signal. The choice of taking the first estimated mode as a reference for the amplitude of the masking signal is aimed at making this amplitude comparable to that of the signal’s random content, which by hypothesis changes faster than its deterministic content. If this is the case, the first extracted IMF is expected to contain mainly noise [17].

#### 2.3.2. EMD based denoising

The denoising procedure starts by estimating the Hurst exponent *H*_*1*_ of the first IMF obtained from the application of masked EMD (which should reflect the features of the signal’s random content, under the hypothesis that the random content of the signal changes faster than its deterministic content [17]). Then for each mode *d*_*k*_ obtained from the application of EMD to the observed signal we test the hypothesis that its energy is larger than that of the mode 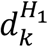 obtained from the application of EMD to a random signal with the Hurst exponent *H*_*1*_. If the null hypothesis that the energies of the two modes are not different cannot be excluded, the mode is flagged as random. To this aim, the energy of each IMF *d*_*k*_, is compared to the energy values of the corresponding IMFs obtained by submitting *r* random signals with Hurst exponent equal to *H*_*1*_ to EMD. The clean signal is finally obtained by summing the modes not flagged as random.

### 2.4. Parametrization

The following parameters were adopted in the analyses reported in Section 3. Demodulation tolerance: *ε* = *e*^−10^; Savitsky Golay differntiator order: *o*_*sg*_ = 5 and length: *l*_*sg*_ = 16; amplitude coefficient of the masking signal: *C*_*A*_ = 8 (which, under the hypothesis that the signal is normally distributed should roughly amount to twice the signal’s range); number of masking signals: *J* = 22; number of random signals generated to estimate the confidence intervals in the denoising algorithm: *r* = 1000 (but results did not changed with *r* = 100); number of masked sifting iterations: *q* = 10 (as suggested in [10]). This choice was adopted also for the sifting operation in applications of the EHHT. Quadrature signals were computed via application of the Hilbert transform.

## 3. Results

### 3.1. Synthetic data

Figure 7 displays the phase obtained by the application of our algorithm to the signal analysed in Figure 4. The topmost panel displays the analysed signal, the second panel from the top displays the first estimation of the signal phase via the Hilbert transform approach after centring (*S*_1_(·)), and refined demodulation (*𝓍*(·)). The third panel from the top displays the output of our algorithm while the bottommost panel displays the phase obtained through the EHHT (based on several iterations of the centring operation) by selecting the first IMF and submitting it to demodulation (*D*(·)).

**Figure 7.**
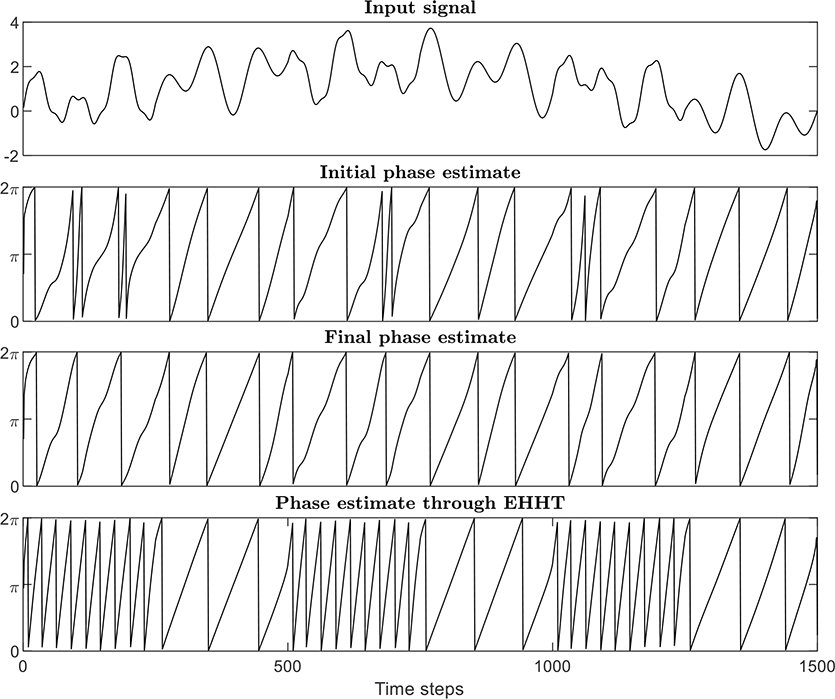
Application of our approach to the analysis of the signal in Figure 4 (here displayed in the topmost panel). Second panel from top: initial estimate of the signal phase after application of centring and refined demodulation. Third panel from top: final estimate of the signal’s phase. Bottommost panel: phase estimated with the approach in [5].

Only the final output of our algorithm permits recovering the oscillatory cycles visible in the signal. However, the comparison between the second and last panels from the top shows that limiting the sifting operator to one iteration of the centring algorithm reduces the sensitivity of the analysis to the small perturbations induced by the weak intermittent component.

### 3.2. Knee flexion data

Figure 8 displays the analysis of a signal representing knee joint angles recorded while participants in standing position had to move rhythmically their torso up and down by flexing and extending their knees and keeping their feet still [26]. The observed cycles, low-pass filtered at 7Hz, have a quite sinusoidal shape. Moreover, the changes in cycles’ amplitudes are relatively slow when compared to their durations. As shown in the second panel from top even the phase obtained by applying the Hilbert transform to the z-scored signal permits the correct identification of all the cycles. However, this signal presents a potentially challenging feature because its values appear to be quantised; therefore, it contains discontinuities all over its duration (not visible in the figure due to insufficient magnification level). Methods based on extrema detection (as those discussed so far) are expected to break down in presence of such discontinuities. Although this issue can be addressed by filtering the signal (as done in computing the phase values displayed in Figure 8), the presence of the discontinuities permits appreciating how sensitive the different methods are to the presence of small non-random fluctuations. Indeed, we can gradually increase the cut-off frequency of the low-pass filter applied to the knee angle signal, count the number of cycles detected in the phase signals obtained through the different approaches, and observe how this number depends on the filter cut-off frequency. We can estimate the number of produced cycles via: 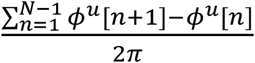, where *N* is the length of the time series and *ϕ*^*u*^[*n*] are the unwrapped phase values.

**Figure 8.**
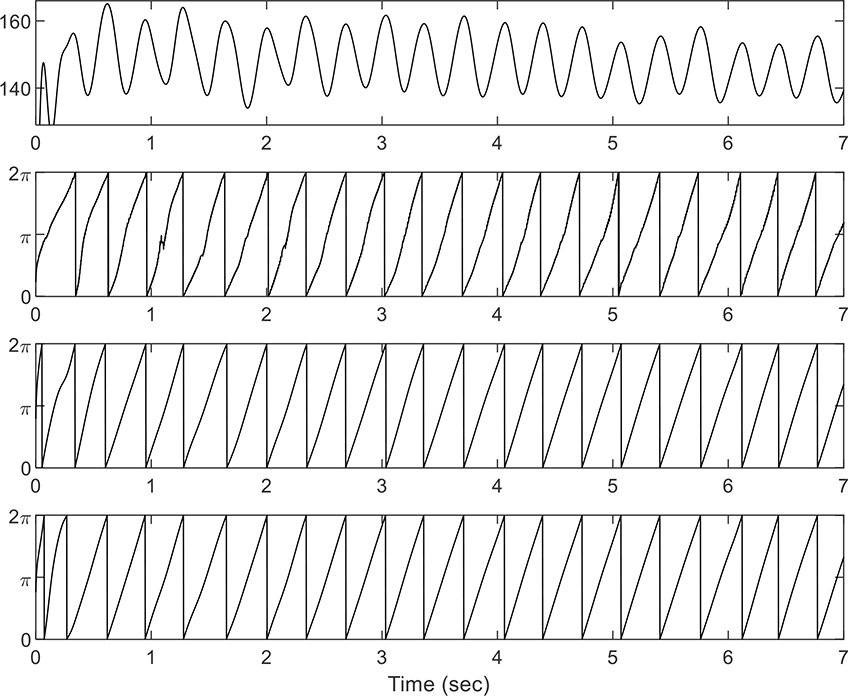
Knee angle data observed during a flexion / extension task. The observed time series is displayed in the topmost panel while the remaining panels display its phase values computed through application of z-score and Hilbert transforms (second panel from top), through the EHHT (third panel from top), and through the method proposed in this paper (bottommost panel).

The cycle counts displayed in Figure 9 were obtained by using 10 integer valued cut-off frequencies uniformly distributed over the interval [7, 120]Hz. While our approach stably approximates the correct number of cycles over a wide range of cut-off frequencies, the approach based on the EHHT starts deviating from the correct behaviour already at the first change of the cut-off frequency value.

**Figure 9.**
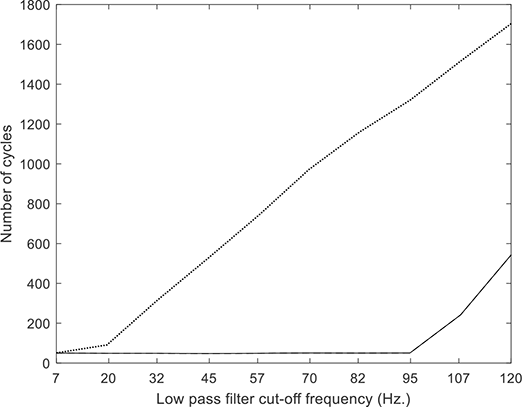
Estimates of number of cycles based on the instantaneous phase values as a function of the cut off frequency of the low pass filter applied to the signal during pre-processing. The continuous line displays the results obtained through the application of our algorithm. The dotted line displays the results obtained through the adoption of EHHT. Both methods identify the correct cycle number when the cut-of freq. is set at 7Hz.

### 3.3. Low pass filtered foot position during gait

The topmost panel of the Figure 10 represents the vertical movement of a sensor attached on the Achille’s tendon during walking. The signals were made available by the authors of the original publication ([27]) low pass-filtered with cut-off frequency at 6Hz. The second panel from the top shows the phase obtained by submitting the z-scored time series to the Hilbert transform. Note that this approach permits capturing the slowing down of phase dynamics observed at low heights. The third and fourth panels from the top display the phase of the first and second IMFs obtained through EMD (and submitted to demodulation). The second IMF captures the cycles, but information about the phase dynamics is lost. The topmost panel of the second column displays the results of masked sifting, and the following panel displays the phase obtained after the application of refined demodulation. The weak cycles appearing in the output of masked sifting are ignored and the phase dynamics can be recovered.

**Figure 10.**
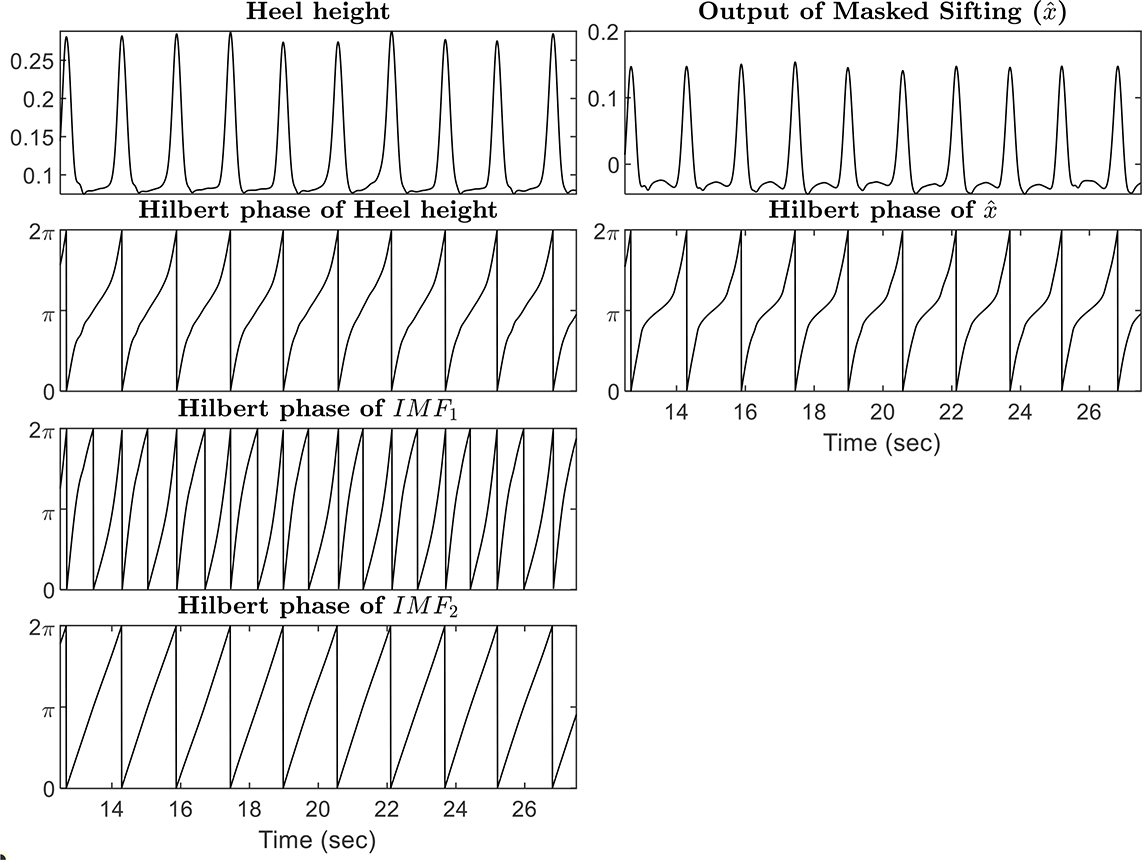
Processing of foot elevation data recorded during gait. Topmost panel of left column input signal, remaining panels from top to bottom: Phase of the z-scored input signal, of the first IMF from application of EMD, of the second IMF. Topmost panel of the right column: input signal submitted to masked sifting, bottom panel: phase obtained after application of refined demodulation.

### 3.4. Tip of the tongue, lower lip and jaw motion in repeated speech production

The signals in the panels on the left side of Figure 11 represent the vertical movement of sensors attached to the tip of the tongue (TTIP, topmost panel), of the jaw (JAW, middle panel) and of the lower lip (LLIP, bottommost panel) collected in [28] via 2D electromagnetic articulography during the repetition of the utterance ‘tapa’ (where the vowel is pronounced like in the word ‘staff’). Speakers were asked to repeat the two syllables without interruption during 10 secs simultaneously to the cues provided by a visual metronome. The metronome frequency increased during the first half of the trial and then gradually came back to its initial value during the second half. Before the analysis, signals were low-pass filtered with a cut-off freq. of 20 Hz to remove measurement noise.

**Figure 11.**
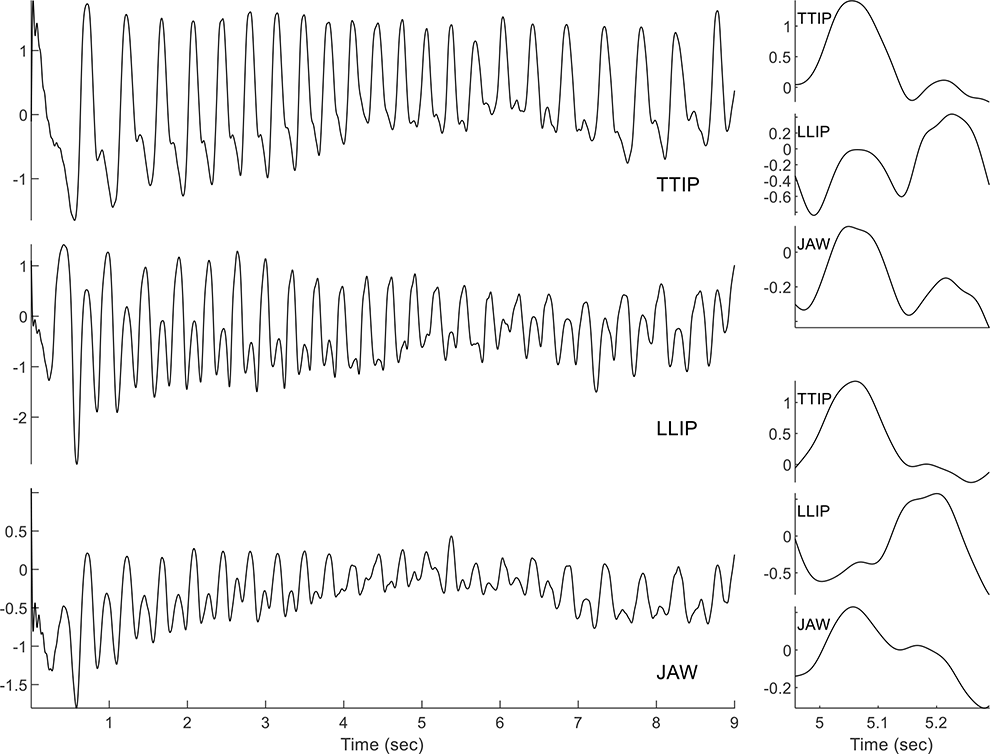
Articulator vertical motion during the repeated production of ‘tapa’ with increasing-decreasing speech rate. The panels on the left display the vertical movement of the tip of the tongue sensor (TTIP: topmost panel), of the lower lip sensor (LLIP: middle panel) and of the jaw sensor (JAW: bottommost panel). The panels of the right column display the trajectories of these three articulators during the production of an utterance at slow speech rate (first three panels from top) and at fast speech rate (last three panels from top).

The panels on the right side of the figure display one slow repetition of the utterance (first three panels from top) and one fast repetition (last three panels from top). At slow speech rate, we can clearly see one main cycle of tongue motion (to produce the consonant ‘t’) followed by a much smaller one, and one main cycle of vertical lower lip motion (to produce the ‘p’) preceded by a slightly smaller one. This pattern is observed because while each articulator organ is activated to produce either ‘t’ or ‘p’, its movement shows the trace of the other articulator producing the other consonant. This occurs because both articulators sit on the jaw, which rotates to favour the upward movement of each articulator and, in doing that, transmits the movement of one articulator to the other. This also explains why the jaw produces two cycles per utterance. Still, the movement of the tip of the tongue seems to have a larger impact on the movement of the other articulators than the cycle of vertical lip motion [28]. At fast speech rate, this difference becomes even larger and the effect of lower lip displacement on the movement of the jaw appears as a perturbation of a larger cycle corresponding to the upward movement of the tip of the tongue. This is consistent with the observation that in this task at fast speech rate the coupling between tip of the tongue and jaw becomes stronger than the coupling between the lower lip and the jaw [29].

The signals displayed in Figure 12 show how the phase values computed through our approach reflect this situation. The topmost panel of the figure displays vertical jaw motion to provide a reference. The phase of the tip of the tongue (second panel from top) captures one cycle per utterance practically all along the sequence. The lower lip phase (third panel from top) displays one cycle per utterance only at fast speech rate. Finally, the jaw (bottommost panel) produces two cycles per utterances during longer portions of the sequence with respect to the lower lip, but as the lower lip, at fast speech rate it displays one cycle per utterance. Note also that at fast speech rate the tip of the tongue and the lower lip are in anti-phase, as it is expected due the alternation of the two consonants. When all articulators display one cycle per utterance, the jaw is in phase with the tip of the tongue. This is expected if the coupling between these two articulator organs is stronger than that between the jaw and the lower lip, as proposed in [29].

**Figure 12.**
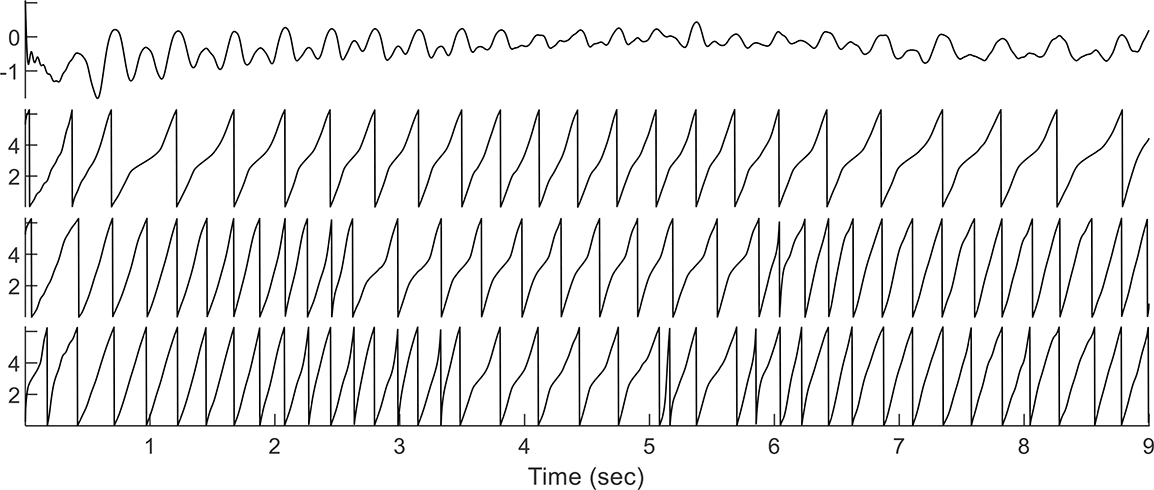
Phase of articulator motion recorded during the repeated production of ‘tapa’. Topmost panel: vertical movement of the jaw. Middle panel: tip of the tongue’s phase. Third panel from top: lower lip’s phase. Bottommost panel: jaw’s phase.

### 3.5. Unfiltered speech articulators data

The odd panels of Figure 13 starting from top display the articulator movements analysed in Section 3.4 before the application of the low-pass filter. The even panels display the phase obtained after the application of the denoising algorithm illustrated in Section 2.30. The results obtained by substituting the low pass filter with the adaptive filtering algorithm closely match those described in Section 3.4.

**Figure 13.**
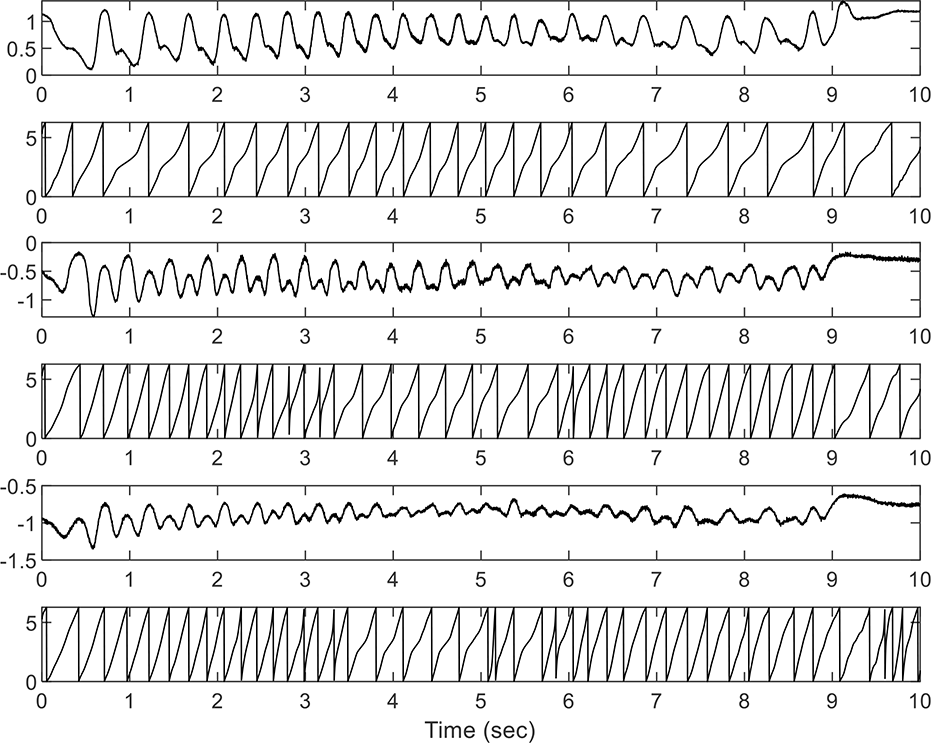
Processing of speech articulator unfiltered data. Odd panels display from top to bottom the upward movements of the jaw, the lower lip and the tip of the tongue. Even panels display the phase angles extracted (see main text for details)

### 3.6. Vocal folds vibration cycles

The signal in Figure 14 represents the vibration of the vocal folds during the production of the phrase “ma mine” from a female French speaker. More precisely, the signal represents a measure of the contact area between the vocal folds, recorded through an electroglottograph. The shape of the opening and closing cycles, characterising the production of voiced sounds such as the vowels and the nasal consonants composing our signal, displays substantial deviation from the shape of a sinusoid (see also the topmost panel of Figure 15). This feature constitutes a problem for the construction of an analytic signal without the application of a decomposition method. Moreover, contrarily to the signals analysed so far, speakers cannot control the instant-by-instant behaviour of glottal opening (although they have some degree of control over the global shape of the cycles, which is associated to different voice qualities).

**Figure 14.**
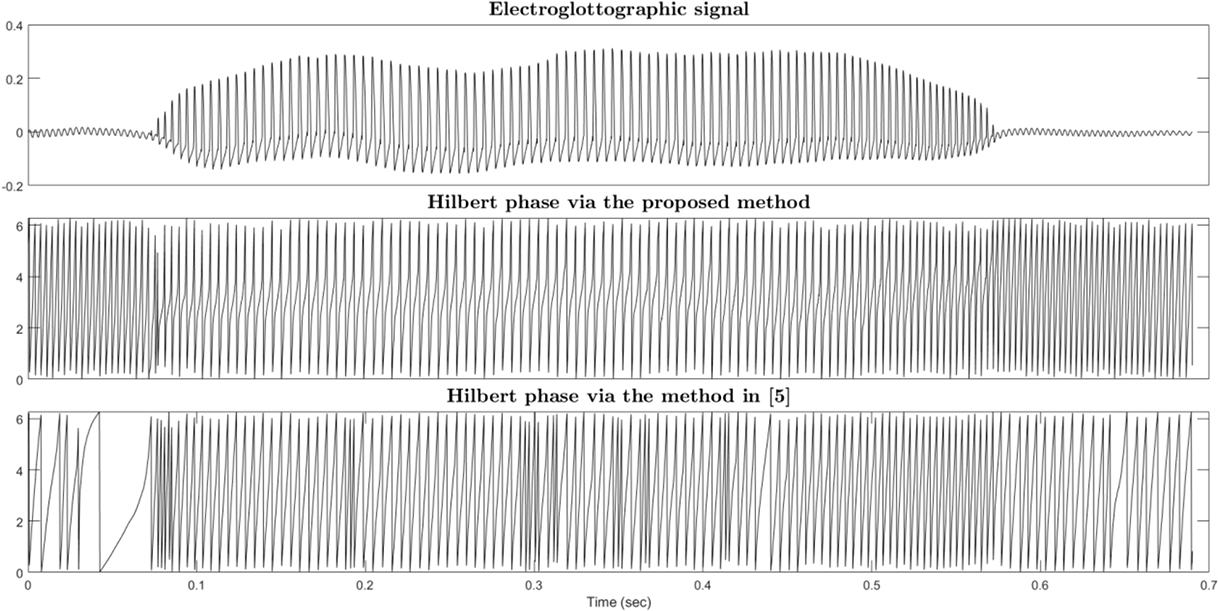
Vocal fold vibration cycles. Topmost panel: estimate of vocal folds’ contact area as captured through an electroglottograph. Second panel from top: Phase of the vocal fold vibration cycles obtained through our algorithm. Third panel from top: Phase obtained through the application of the EHHT by selecting the second IMF.

**Figure 15.**
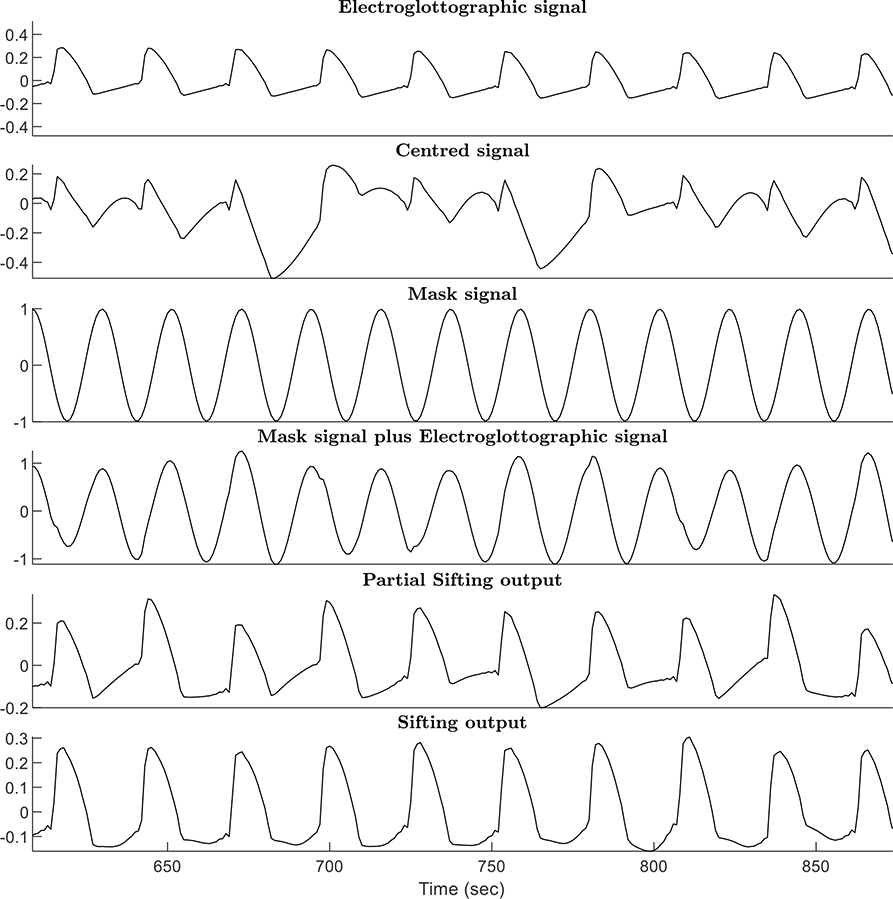
Signals produced in the analysis of a portion of the vocal folds vibration signal in Figure 14. Topmost signal: Vocal folds’ vibration. Second signal from top: vocal folds’ vibration centred on zero. Third signal from top: masking signal with ϕ_0_ = 0. Fourth signal from top: vocal folds’ vibration plus masking signal. Fifth signal from top: partial output of the masked sifting process after removal of the masking signal. Bottommost signal: final output of masked sifting.

Still, the signal displays one main peak per cycle and our algorithm is able to capture the main component of the vocal folds vibration cycles, as seen in the second panel from the top in Figure 14. The result obtained through the EHHT are much less robust due to mode splitting. Note, that even using variants of EMD (e.g.: [30]) that with this kind of signals prevent the occurrence of mixing/splitting phenomena, would not solve the problem of selecting the correct IMF, given that this may change from one chunk of signal to the other.

Figure 15 shows the signals produced in the process of extracting the phase through our algorithm. The topmost panel displays a portion of the signal in Figure 14. The second panel from the top shows the centred signal, featuring two kinds of cycles: a smoother one, reflecting the slow varying portions of the vocal fold vibration cycles, and a shaper one reflecting the faster changing portions.

The estimated frequency of this signal will be higher or equal to that of the vocal folds’ vibration cycle, but not faster than the local instantaneous frequency observed around the maximum contact instants. One masking signal produced with the observed frequency is shown in the third panel from top of Figure 15 and the signal submitted to the sifting operation in the panel just below. This signal displays the cycles of the masking signal, deformed by the input signal. Slow changes of the input signal result in changes of amplitude of the summed signal. As such, they are captured by the envelope computed during the sifting process and removed from the signal (see the second panel from the bottom in Figure 15). Variability in the partial IMF due to the phase difference between the masking signal and the input signal is canceled by averaging between the results obtained by using different initial phase values for the masking signals (see the bottommost panel of Figure 15).

## 4. Discussion and conclusions

The method proposed in the present paper permits extracting a time continuous estimate of the phase from oscillatory signals whose behaviour deviate from that of a narrowband signal with relatively slow changing amplitude. Moreover, it permits the analysis of patterns of activity deviating to some extent from a sinusoidal shape.

The example in Section 3.1 show how one iteration of sifting permits a more robust estimate of the signal phase with respect to a full sifting in presence of weak or sparse oscillations faster than the oscillations of interest. This explains the robustness of the obtained phase values to the presence weak components with frequencies higher than that of the target oscillation observed in section 3.2. The examples in section 3.4 show how the proposed approach permits isolating a dominant component in the presence of a weaker or more sparse deterministic component. So the tip of the tongue movement always shows one cycle per utterance because the effect of the lower lip movement on the TTIP sensor is relatively weak. This is not the case for the effect of the tip of the tongue movement on the position of the LLIP or of the JAW sensors, which is relatively strong at slow speech rate, and therefore induces the method to detect two cycles per utterance in the movement of these sensors. In Section 3.5, we showed that the application of adaptive noise removal based on masked EMD permits obtaining the same results in the presence of measurement noise. The example in section 3.1, illustrating the behaviour of the proposed method with synthetic data, deserves some additional comment. Indeed the main component of the observed signal (displayed in the third panel from top in the leftmost column of Figure 4) does not display the variability in phase dynamics observed in the output of our method (third panel from top of Figure 7). This occurs because the deformations of the main cycle, that in the synthetic signal are due to the intermittent weak activity of the fastest component, are mainly considered as due to local changes in amplitude or frequency of the dominant oscillatory patterns. Despite in the case illustrated in Section 3.1, this interpretation leads to mix features of two different components in one sequence of phase values, it permits a correct characterization of shapes that deviate from oscillatory behaviour as in the case of the pulse signal analysed in Section 3.2 or of the vibration signal analysed in Section 3.6. Importantly, when the task under study is defined by a pattern of behaviour of an observed quantity, variability in the temporal features of the observed quantity affects the task performance and therefore it should be reflected in the behaviour of the computed phase. Even when the perturbations may be due to an underlying component oscillating faster than the dominating component (as it is the case for the signal analysed in Section 3.1).

The results of our work have implications that extend beyond its scope and should be considered in the context of decomposition strategies. More specifically, our approach lead to a revised version of the masked EMD strategy [24, 25] in which the dominant frequency of each component needs to be estimated before the application of the masked sifting operator. We propose to estimate this frequency from the signal obtained by application of one iteration of sifting and of the refined demodulation operation. Future work will be devoted to investigate the benefits this approach for the decomposition of general multicomponent signals.

## 5. Acknowledgments

This work, carried out within the Institute of Convergence ILCB, was supported by grants from *France 2030* (ANR-16-CONV-0002) and the Excellence Initiative of Aix-Marseille University (A*MIDEX) and by grant ANR-21-CE28-0015-01 of the French National Research Agency.

## Notes

https://github.com/LeonardoLancia/Get_Phase_Tools

### Competing Interest Statement

The authors have declared no competing interest.

### Summary of Updates

Code avialability on GitHub and consistency check between Matlab and Python versions

## References

[1] Schöner G, Kelso JA. A synergetic theory of environmentally-specified and learned patterns of movement coordination: I. Relative phase dynamics. Biological cybernetics. 1988 Jan;58(2):71–80.

[2] Kelso JS. Dynamic patterns: The self-organization of brain and behaviour. MIT press; 1995.

[3] Longo G. Confusing biological rhythms and physical clocks Today’s ecological relevance of Bergson-Einstein debate on time. Eintein vs Bergson. An enduring quarrel of time. 2021.

[4] Pikovsky A, Rosenblum, M, Kurths J. Synchronization: a universal concept in nonlinear science. Cambridge University Press; 2003

[5] Huang NE, Wu Z, Long SR, Arnold KC, Chen X, Blank K. On instantaneous frequency. Advances in adaptive data analysis. 2009 Apr;1(02):177–229.

[6] Rosenblum M, Pikovsky A, Kurths J, Schäfer C, Tass PA. Phase synchronization: from theory to data analysis. In Handbook of biological physics 2001 Jan 1 (Vol. 4, pp. 279–321). North-Holland.

[7] Mörtl A, Lorenz T, Hirche S. Rhythm patterns interaction-synchronization behaviour for human-robot joint action. PloS one. 2014 Apr 21;9(4):e95195.

[8] Lancia L, Chaminade T, Nguyen N, Prevot L. Studying the link between inter-speaker coordination and speech imitation through human-machine interactions. InInterspeech 2017 2017 Aug 20.

[9] Tuller B, Kelso JS. The timing of articulatory gestures: Evidence for relational invariants. The Journal of the Acoustical Society of America. 1984 Oct 1;76(4):1030–6.

[10] Chavez M, Besserve M, Adam C, Martinerie J. Towards a proper estimation of phase synchronization from time series. Journal of Neuroscience methods. 2006 Jun 30;154(1-2):149–60.

[11] Huang NE, Shen Z, Long SR, Wu MC, Shih HH, Zheng Q, Yen NC, Tung CC, Liu HH. The empirical mode decomposition and the Hilbert spectrum for nonlinear and non-stationary time series analysis. Proceedings of the Royal Society of London. Series A: mathematical, physical and engineering sciences. 1998 Mar 8;454(1971):903–95,

[12] Wang D, Narayanan SS. Robust speech rate estimation for spontaneous speech. IEEE Transactions on Audio, Speech, and Language Processing. 2007 Oct 15;15(8):2190–201.

[13] Goldstein L. The role of temporal modulation in sensorimotor interaction. Frontiers in Psychology. 2019 Dec 6;10:2608.

[14] Goswami U. Speech rhythm and language acquisition: an amplitude modulation phase hierarchy perspective. Annals of the New York Academy of Sciences. 2019 Oct;1453(1):67–78.

[15] James EG. Nonstationarity of stable states in rhythmic bimanual coordination. Motor Control. 2014 Apr 1;18(2):184–98.

[16] Melanson A, Mejias JF, Jun JJ, Maler L, Longtin A. Nonstationary stochastic dynamics underlie spontaneous transitions between active and inactive behavioural states. Eneuro. 2017 Mar 1;4(2).

[17] Flandrin P, Goncalves P, Rilling G. Detrending and denoising with empirical mode decompositions. In2004 12th European signal processing conference 2004 Sep 6 (pp. 1581–1584). IEEE.

[18] Varlet M, Richardson MJ. Computation of continuous relative phase and modulation of frequency of human movement. Journal of biomechanics. 2011 Apr 7;44(6):1200–4.

[19] Wu Z, Huang NE. On the filtering properties of the empirical mode decomposition. Advances in Adaptive Data Analysis. 2010 Oct;2(04):397–414.

[20] Wang G, Chen XY, Qiao FL, Wu Z, Huang NE. On intrinsic mode function. Advances in Adaptiv Data Analysis. 2010 Jul;2(03):277–93.

[21] Deering R, Kaiser JF. The use of a masking signal to improve empirical mode decomposition. InProceedings.(ICASSP’05). IEEE International Conference on Acoustics, Speech, and Signal Processing, 2005. 2005 Mar 23 (Vol. 4, pp. iv–485). IEEE.

[22] Bueno-López M, Giraldo E, Molinas M, Fosso OB. The Mode Mixing Problem and its Influence in the Neural Activity Reconstruction. IAENG International Journal of Computer Science. 2019 Aug 1.

[23] Sandoval S, De Leon PL. Advances in empirical mode decomposition for computing instantaneous amplitudes and instantaneous frequencies. In2017 IEEE International Conference on Acoustics, Speech and Signal Processing (ICASSP) 2017 Mar 5 (pp. 4311–4315). IEEE.

[24] Fabus MS, Quinn AJ, Warnaby CE, Woolrich MW. Automatic decomposition of electrophysiological data into distinct nonsinusoidal oscillatory modes. Journal of Neurophysiology. 2021 Nov 1;126(5):1670–84.

[25] Wang YH, Hu K, Lo MT. Uniform phase empirical mode decomposition: An optimal hybridization of masking signal and ensemble approaches. Ieee Access. 2018 Jun 15;6:34819–33.

[26] Miyata K, Kudo K. Mutual stabilization of rhythmic vocalization and whole-body movement. PloS one. 2014 Dec 12;9(12):e115495.

[27] Hebenstreit F, Leibold A, Krinner S, Welsch G, Lochmann M, Eskofier BM. Effect of walking speed on gait sub phase durations. Human movement science. 2015 Oct 1;43:118–24.

[28] Rochet-Capellan A, Schwartz JL. An articulatory basis for the labial-to-coronal effect:/pata/seems a more stable articulatory pattern than/tapa. The Journal of the Acoustical Society of America. 2007 Jun 1;121(6):3740–54.

[29] Lancia L, Rosenbaum B. Coupling relations underlying the production of speech articulator movements and their invariance to speech rate. Biological Cybernetics. 2018 Jun;112:253–76.

[30] Colominas MA, Schlotthauer G, Torres ME. Improved complete ensemble EMD: A suitable tool for biomedical signal processing. Biomedical Signal Processing and Control. 2014 Nov 1;14:19–29.

