## Appendix for "Instantaneous phase of rhythmic behaviour under volitional control"

### Appendix: implementation details

##### A.1. Sifting

The sifting operator (indicated by $S_q(x)$) takes as argument x: a real valued signal of length N and sampling frequency $sr$ and, in the notation adopted here, it is parameterized by the positive integer $q$.

Let $maxs(\cdot)$ be the function that takes as argument a real valued unidimensional signal and returns its local maxima.

Let $interp(t_0, v, t_1)$ represent the operator that interpolates at time points $t_1$ the values in $v$ observed at time points $t_0$. by using monotone piecewise cubic interpolation [1].

Let $isIMF(\cdot)$ represent the operator that accepts as argument a real valued signal and outputs one if it fulfils the criteria that define an IMF according to [2] and 0 otherwise.

| |
|---|
| $r = x$ |
| $t = \{1, \dots, N\} \times \frac{1}{sr}$ |
| $stopIt = 0$ |
| $nIter = 0$ |
| while st$opIT == 0$ and $nIter \leq q$ |
| set $le^{+} = \arg maxs(r)$ |
| set $le^{-} = \arg maxs(-r)$ |
| $minEnv = interp(t[le^{-}], x[le^{-}], t)$ |
| $maxEnv = interp(t[le^{+}], x[le^{+}], t)$ |
| $\tilde{r} = \mathrm{r} - (minEnv + maxEnv)/2$ |
| if $isIMF(\tilde{r})$ |
| $stopIt = 1$ |
| else |
| $r = \tilde{r}$ |
| end if |
| end while |

###### A.1.1. Stopping criteria and processing of boundary conditions.

Concerning the specific criteria adopted to determine if $\tilde{r}$ is an IMF, we followed [2] who proposes a method based on two thresholds $\vartheta_1$ and $\vartheta_2$ and one tolerance parameter $\alpha$. The sifting is interrupted when:

$$\left|\frac{meanEnv[n]}{maxEnv[n]-minEnv[n]}\right| < \theta_2, \forall n \in \{1, \dots, N\},$$

and $\left\langle \left|\frac{meanEnv[n]}{maxEnv[n]-minEnv[n]}\right| < \theta_1 \right\rangle > \alpha$.

Where $meanEnv = (maxEnv + minEnv)/2$.

Another implementational aspect of sifting, as well of all the other operations involving the interpolation of local extrema, concerns the processing of the boundary conditions. In order to obtain the signal values that precede the first local extremum and follow the last one, we again followed [2] who propose mirroring the extrema at the boundaries (we chose to mirror the first/last two extrema).

The Matlab code provided in the supplementary material is based on the 2007 implementation of these criteria, available at the following address: http://perso.ens-lyon.fr/patrick.flandrin/emd.html. The Python port of this code (used in the Python implementation of our algorithm) is available in the package pyEMD (https://github.com/laszukdawid/PyEMD ) through the function 'EMD_matlab.py'.

##### A.2. Refined Demodulation

Let sort() be the function that takes as input a vector of real values and sorts them in ascending order.

| |
|---|
| set $StopIt = 0$ |
| while $nIter < maxIterN$ and $stopIt == 0$ |
| $el^{+} = \arg maxs\ (x)$ |
| $el^{-} = \arg maxs\ (-r)$ |
| $el = sort(el^{-} \cup\ el^{+})$ |
| $Z = length(el))$ |
| $z \in \{1,2, \ldots, Z\}$ |
| $e = interp(t[el], \lvert x[el] \rvert, t)$ |
| $r = \ r/e$ |
| if $\lvert r[el[z]] \rvert \leq 1 - \epsilon, \forall z \in\ \{1,2, \ldots, Z\}$ |
| $stopIt = 1$ |
| end if |
| end while |

##### A.3. Masked sifting

Masked sifting takes as arguments $x$: a real valued signal of length $N$ and sampling frequency $sr$; $\omega_{mask}\ \in R^{+} | \omega_{mask} < sr/2$; $A_{mask} \in R^{+}$; and $n_{masks} \in N^{+}$; and it is indicated as $MS_q(x, \omega_{mask}, A_{mask}, n_{masks})$, where the index $q$ refers to the maximum number of iterations of the sifting process $S_q(\cdot)$.

First, the partial sifts of the signal modified by the different masking signal are computed and the masking signals subtracted from the results:

$\breve{x}_j = S_q(mask_j + x)$- $mask_j$ , $\forall j\ \in \{1, .., J\}$ and $\forall n\ \in \{1, .., N\}$

with $mask_j[n] =\ A_{mask} cos(n \Delta t \omega_{mask} + \phi_{0,j})$. Then the obtained signals are averaged:

$\breve{x} = \frac{1}{J} \sum_{j=1}^{J} \breve{x}_j$.

##### A.4. Adaptive denoising algorithm

To clean the input signal from high frequency random noise, we first apply Masked EMD in which classical sifting is substituted by masked sifting.

###### A.4.1. Masked EMD

Let $r_d$ represent the input signal at $d^{th}$ iteration of the algorithm ($r_1 = x$).

Let $S_h(\cdot)$ represent the sifting operator, taking as argument a real valued signal of length N, with $h$ indicating the maximum number of iteration of the sifting process.

Let $\varkappa_k(\cdot)$ represent the demodulation procedure in A.2, with $k$ indicating the maximum number of iterations of the normalization operation.

Let $MS_q(\cdot, \omega_{mask}, A_{mask}, n_{masks})$ represent the masked sifting operation accepting as arguments a real valued signal, a frequency value $\omega_{mask}$, a relative amplitude value $A_{mask}$, and an integer representing the number of masking signals to be applied $n_{masks}$, with $q$ indicating the maximum number of iterations of the of the centring operation.

Let $length(\cdot)$ represent the operator that gives the number of elements of its argument.

The following pseudocode illustrates the masked EMD algorithm. Note that the free parameter $Ac$ determines the amplitude of the masking signal as a proportion of the range of the input signal after the application of the centering operator.

| |
|---|
| stopIt=0 |
| d=1 |
| $r_d = x$ |
| while stopIt =0 |
| $\tilde{r}_d = S_1(r_d)$ |
| $\hat{r}_d = \varkappa_5(\tilde{r}_d)$ |
| $y_d = \mathcal{H}(\hat{r}_d)$ |
| $\omega_{\hat{r}_d} = \langle SGd(tan^{-1}\left(\frac{y_d}{\hat{r}_d}\right), o_{sg}, l_{sg})\rangle$ |
| If d==1 |
| $A_{\hat{r}} = Ac(\max(\tilde{r}_d) - \min(\tilde{r}_d))$ |
| end if |
| $\check{r}_d = MS_{10}(r_d, \omega_{\hat{r}_d}, A_{\hat{r}}, nMasks)$ |
| $\hat{r}^{\cdot}_d = \varkappa_5(\check{r}_d)$ |
| $y^{\cdot}_d = \mathcal{H}(\hat{\hat{r}}^{\cdot}_d)$ |
| $\omega_{\hat{r}^{\cdot}_d} = \langle SGd\left(tan^{-1}\left(\frac{y^{\cdot}_d}{\hat{r}^{\cdot}_d}\right), o_{sg}, l_{sg}\right)\rangle$ |
| $r_{d+1} = r_d - \check{r}_d$ |
| $e^+ = maxs(x)$ |
| $e^- = maxs(-x)$ |
| If length($e^+ \cup e^-$)<1 |
| stopIt=1 |
| end if |
| end while |
| nIMF=d |

A.4.2. IMFs significance test and reconstruction of the clean signal

We estimate the Hurst exponent of the first IMF. We check that the energy of the obtained modes is significantly different from that of the modes obtained by decomposing random signals with the Hurst exponent computed. Finally we average the IMFs that passed the test.

Let $Hurst(\cdot)$ be the operator that gives the Hurst exponent of its argument.

Let $getHurstCI_k(H_0, n_H, a_H)$ be the operator that:

1) generates $n_H$ realizations of a random process with Hurst exponent $H_0$ and amplitude $a_H$;
2) extracts the $k^{th}$ IMF from each realization;
3) computes the boundaries delimiting the 95% confidence interval of the energy values observed across realizations of that IMF.

The following lines illustrate the remaining steps of the denoising procedure.

| |
|---|
| $H_0 = husrt(\check{r}_1)$ |
| for $k$ $in$ $\{1, \ldots, nIMF\}$ |
| $lowCI, hiCI = getHurstCI_k(H0, nh, ah)$ |
| if $\sqrt{\frac{\sum_{n=1}^{N} \check{r}_d[n]}{N}} \leq hiCI$ |
| $flag_d = 0$ |
| else |
| $flag_d = 1$ |
| end if |
| end for |
| y=$\sum_{d=1}^{N} flag_d \check{r}_d$ |

#### A.5. References

[1] Fritsch FN, Butland J. A method for constructing local monotone piecewise cubic interpolants. SIAM journal on scientific and statistical computing. 1984 Jun;5(2):300-4

[2] Rilling G, Flandrin P, Goncalves P. On empirical mode decomposition and its algorithms. InIEEE-EURASIP workshop on nonlinear signal and image processing 2003 Jun 8 (Vol. 3, No. 3, pp. 8-11). Grado: IEEE.
