## Supplementary material for "Instantaneous phase of rhythmic behaviour under volitional control": Code and data files: get_closest_Vals_Idxs.m

```
function [mapping, mapIdxs]=get_closest_Vals_Idxs(target,inventory)
%given two sets of time points it creates the vector 'mapping' by substituting each value in
%the first set with the nearest value in the second set. the vector mapIdxs
%contains the indexes that the values included in 'mapping' had in the second set

% Leonardo Lancia 14/10/2023
% mailto:

targShape=size(target);
invShape=size(inventory);
[~,lenDimTarg]=max(targShape);
[~,lenDimInv]=max(invShape);
if lenDimTarg~=lenDimInv && max(targShape)~=2 && lenDimInv>1
   error('target and inventory must have same orientation and length')
end
if min(targShape)>1
    if diff(size(inventory))<0
        newTarget=target(:)';
    else
        newTarget=target';
        newTarget=newTarget(:)';
    end
elseif diff(targShape)<0
    newTarget=target';
else
    newTarget=target;
end

if diff(size(inventory))<0
    newInventory=inventory';
else
    newInventory=inventory;
end
myDist=distance(newTarget,newInventory);
[~,mapIdxs]=min(myDist);
mapping=inventory(mapIdxs)';
mapIdxs=mapIdxs';
if diff(targShape)<0
    mapping=reshape(mapping,targShape);
    mapIdxs=reshape(mapIdxs,targShape);
else
    mapping=reshape(mapping,fliplr(targShape))';
    mapIdxs=reshape(mapIdxs,fliplr(targShape))';
end
```
