## Supplementary material for "Instantaneous phase of rhythmic behaviour under volitional control": Code and data files: get_envelopes.m

```
function [maxEnv, minEnv,meanEnv]= get_envelopes(TS0)
% computes the minimum, maximum and mean envelopes of a time series

    [myPckVs,myPcks]=findpeaks(TS0);%find maxima
    [myValVs,myVals]=findpeaks(-TS0);%find minima
    myValVs=-myValVs;%reverse minima

    lenTS=length(TS0);
    % copy first and last valleys to initial and final time series positions
    % this avoids mess at the edges with interpolation
    [myValVs,myVals]=SetBoundCond(myValVs,myVals,lenTS);
    % copy first and last peaks to initial and final time series positions
    [myPckVs,myPcks]=SetBoundCond(myPckVs,myPcks,lenTS);
    
    idxPV=1:lenTS;% vector with all locations of time series values
    maxEnv=interp1(myPcks',myPckVs',idxPV,'pchip');% maximum envelope
    minEnv=interp1(myVals',myValVs',idxPV,'pchip');% minimum envelope
    meanEnv=(maxEnv+minEnv)./2;%mean envelope
end
```
