## Supplementary material for "Instantaneous phase of rhythmic behaviour under volitional control": Code and data files: get_mfcc.m

```
function [MFCCs FBEs] = get_mfcc (s,sr,shift)
% just a wrapper for mfcc.m
    
%parameters setting
Tw = 25;     % window length (ms)
Ts = shift;  % window hop size (ms)
alpha = 0.97;% preemphasis 
M = 20;      % number of filterbank channels 
C = 12;      % number of cepstral coefficient submitted to differentiation
L = 22;      % cepstral sine lifter parameter
LF = 300;    % smallest spectral freq. considered (Hz)
HF = 3700;   % largest spectral freq. considered (Hz)

[ MFCCs, FBEs, frames ] = mfcc( s, sr, Tw, Ts, alpha, @hamming, [LF HF], M, C+1, L );
```
