## Supplementary material for "Instantaneous phase of rhythmic behaviour under volitional control": Code and data files: get_PLV.m

```
function [PLV,PLVidx, n_theta1,m_theta2]=get_PLV(theta1,theta2,winLen,winStep,maxord)
% given two unidimensional time series representing phase angles varying over time,
% it gets the phase locking value on partially overlapping chunks of time series
% 
% Input:
% theta1: phase angles of the first time series
% theta2: phase angles of the second time series
% winLen: window length (if winStep>0, winLen==0 is interpreted as winLen=length(theta1)
% winStep: window step (if winLen>0, winStep==0 is interpreted as a timeStep to big to get two measures)
% maxord: maximum order of the multiplicative coefficient to be used in the
%         computation of the gerenalize phase difference (if maxOrd==1,
%         theta1 and theta2 are hypothesized to have the same frequency, 
%         if two values are provided m and n will be fixed at these two values )
% output:
% PLV: phase locking value
% PLVtStep: time staps (in number of frames corresopnding to the middle points
%       of the analysis window)
% n_theta1: optimal multicative coefficients n (one per analysis window)
% m_theta2: optimal multicative coefficients y (one per analysis window)
% 
% Trick (yet to be tested):
% to get arbitrary time windows (with variable duration and time step)
% winLen and winStep must be two sequences of integer values with the same
% length H representing the starting frame and the end frame of each time window.
% Of course winLen(h)<winStep(h) for each h in {1,...,N} 

% Leonardo Lancia 14/10/2023
% mailto:


N=length(theta1);
if length(theta2)~=N
   error('the two angles vectors must have the same length') 
   return
end
if length(winLen)==1 && length(winStep)==1
    if winLen==0 | winStep==0
        winLen=length(theta1);
        winStep=winLen;
    end
    startPts=1:winStep:N;
    endPts=winLen:winStep:N;
%     PLVidx=floor(winLen/2):winStep:N;
    minLen=min([length(startPts),length(endPts)]);
    PLVidx=floor((startPts(1:minLen)+endPts(1:minLen))./2);
    PLV=nan(length(PLVidx),1);
    n_theta1=nan(length(PLVidx),1);
    m_theta2=nan(length(PLVidx),1);
    startPt=1:winStep:N;
    endPt=winLen:winStep:N;
    if length(startPt)>length(endPt)
        startPt(length(endPt)+1:end)=[];
    end
elseif length(winLen)>1 && length(winStep)>1
    if length(winLen)~=length(winStep)
            error('either third and fourth arguments are both vector and have same length, or they are both scalars');
    end
    if any(winLen>winStep)
        error('if third and fourth arguments are sequences each element of the third arg must be smaller than the corresponding element of the fourth argument')
    end
    startPt=winLen;
    endPt=winStep;
    PLVidx=floor(mean([startPt(:),endPt(:)],2));

else
    error('either third and fourth arguments are both vector and have same length, or they are both scalars');
end   
    
    
for i=1:length(PLVidx)
%     endPt=min(N,ceil(startPt(i)+winLen(i)));
    thisTheta1=theta1(max([1,floor(startPt(i))]):endPt(i));
    thisTheta2=theta2(max([1,floor(startPt(i))]):endPt(i));
    if length(maxord)==1
        PLV(i)=0;
        for n = 1 : maxord
            for m = 1 : maxord
                tmpPLV=abs(mean(exp(1i*( n*thisTheta1 - m*thisTheta2) )));%co_sync(thisTheta1, thisTheta2, n, m);  % Computing the matrix of synchronizations indices
                if tmpPLV > PLV(i)
                    PLV(i)=tmpPLV; n_theta1(i)=n; m_theta2(i)=m;
                end
            end
        end      
%         [~, PLV(i), n_theta1(i), m_theta2(i)]=co_maxsync(thisTheta1, thisTheta2, maxord);
    else
        PLV(i)=abs(mean(exp(1i*( maxord(1)*thisTheta1 - maxord(2)*thisTheta2) )));%co_sync(thisTheta1, thisTheta2,maxord(1), maxord(2));  
        n_theta1(i)=maxord(1);
        m_theta2(i)=maxord(2);
    end
%     startPt=startPt+winStep;
end
```
