## Supplementary material for "Instantaneous phase of rhythmic behaviour under volitional control": Code and data files: interp_NAN.m

```
function [newX,justnans]=interp_NAN(X,method)
%interpolate nan values with the interpolation method in 'method'

% Leonardo Lancia 14/10/2023
% mailto:


if nargin<2 || isempty(method)
    method='linear';
end
% if diff(size(X))>0
%     X=X';
%     turnit=1;
% else
%     turnit=0;
% end
firstnonNan=find(~any(isnan(X),2),1,'first');
lastnonNan=find(~any(isnan(X),2),1,'last');
if lastnonNan<length(X)
    X(end)=X(firstnonNan);
end
if firstnonNan>1
    X(1)=X(firstnonNan);
end
if strcmp(method,'linear')
    Xtrim=X;
    mynans=isnan(Xtrim);
    newXtrim=Xtrim;
    justnans=nan(size(Xtrim));
    justnans(mynans) = interp1(find(~mynans), Xtrim(~mynans), find(mynans), method,'extrap');

    newXtrim(mynans) = justnans(mynans);

    newX=[nan(firstnonNan-1,size(X,2)); newXtrim ; nan(size(X,1)-lastnonNan,size(X,2))];
    justnans=[nan(firstnonNan-1,size(X,2));justnans ; nan(size(X,1)-lastnonNan,size(X,2))];

else
    newX=X;
	mynans=isnan(X);
    justnans=nan(length(X),1);
    justnans(mynans) = interp1(find(~mynans), X(~mynans), find(mynans), method,'extrap');
    newX(mynans) = justnans(mynans);
end

% if turnit==1
%    newX=newX';
%    justnans=justnans';
% end
```
