## Supplementary material for "Instantaneous phase of rhythmic behaviour under volitional control": Code and data files: normalize_env_peaks.m

```
function normSig=normalize_env_peaks(sig,threshNorm,maxIterN, symN)
%    normalize envelope peaks iteratively following Huang et al. (2009) and using pchip
%    interpolation to obtain the envelope connecting the abs. values of local extrema.
%    Boundary conditions of the local maxima sequence are computed following 
%    Rilling 2007 (see function in '.\utils\boundary_conditions.m').
%     input:
%         sig: the input signal
%         threshNorm: threshold to determine if peaks are different from 1/-1
%         maxIterN: maximum number of iterations
%         symN: number of values to be added at the beginning and at the
%               end before interpolating.
%     output:
%         normalized envelope
% 

if nargin <2 || isempty(threshNorm) || isnan(threshNorm)
    threshNorm=1e-10;
end
if nargin<3 || isempty(maxIterN)
    maxIterN=3;
end
if nargin<4 || isempty(symN)
    symN=2;
end
normSig=sig;
newIdxs=1:length(normSig); %input signal's time steps 
iter_val=1; %intitialize error value
niter=0;% current iteration number

while niter<maxIterN && iter_val>threshNorm
    niter=niter+1;%update iteration number
    [~,pksLoc]=findpeaks(abs(normSig)); %find local maxima 
    [~,valLoc]=findpeaks(-abs(normSig));% local minima are only computed for the sake to apply the boundary_conditions function
	%set boundary conditions via Rilling's function

    [tMin,tMax,sigMin,sigMax] = boundary_conditions(valLoc',pksLoc',newIdxs,abs(normSig)',abs(normSig)',symN);

    if isempty(tMax)% if no maxima break and return
        normSig=normSig/range(normSig);
        return 
    end

    refSig= interp1(tMax,sigMax,newIdxs, 'pchip'); %interpolate local maxima
    normSig=normSig./refSig'; %normalize signal's amplitude
    
	iter_val = abs(sum(refSig) - max(size(refSig))) ;% compute mean difference between envelopes values from last two iterations
end
```
