## Supplementary material for "Instantaneous phase of rhythmic behaviour under volitional control": Code and data files: sample_advance.m

```
%%%% function to time-advance signal, preserve buffer length and pad with externally defined value
function op=sample_advance(ip, nshift, padding)
if nargin<3
    padding=1e-7;
end
[M,N]=size(ip);
if N>M
    transl=1;
    ip=ip';
else
    transl=0;
end
siglen=size(ip,1);
ip(1:siglen-nshift,:) = ip(nshift+1:siglen,:); %%%%% shift the bulk
ip(1+siglen-nshift:siglen,:) = padding; %%% obliterate bit at end with externally-defined fill
if transl==1
    op = ip';
else
    op = ip;
end
```
