## Supplementary material for "Instantaneous phase of rhythmic behaviour under volitional control": Code and data files: SetBoundCond.m

```
function [vals,locs]= SetBoundCond(vals, locs,lenTS)
%set boundary conditions of a sequence of extrema values (local max or min)
%by repeating the first/last observed peaks/valleys
%     
%input:
%   vals: extrema values
%   myValLocs: extrema locations
%   locs: length of the time series from which the extrema were extracted
%         
%output:
%	vals : new sequence of extrema values
%   locs : new sequence of extrema location

if locs(1)>1% if first extremum is not at initial position
    locs=[1;locs];
    vals=[vals(1);vals];
end

if locs(end)<lenTS % if last extremum is not at final position
    locs=[locs;lenTS];
    vals=[vals;vals(end)];
end

end
```
