## Supplementary material for "Instantaneous phase of rhythmic behaviour under volitional control": Code and data files: subplot_tight.m

```
function vargout=subplot_tight(m, n, p, margins, varargin)
%% subplot_tight
% A subplot function substitude with margins user tunabble parameter.
%
%% Syntax
%  h=subplot_tight(m, n, p);
%  h=subplot_tight(m, n, p, margins);
%  h=subplot_tight(m, n, p, margins, subplotArgs...);
%
%% Description
% Our goal is to grant the user the ability to define the margins between neighbouring
%  subplots. Unfotrtunately Matlab subplot function lacks this functionality, and the
%  margins between subplots can reach 40% of figure area, which is pretty lavish. While at
%  the begining the function was implememnted as wrapper function for Matlab function
%  subplot, it was modified due to axes del;etion resulting from what Matlab subplot
%  detected as overlapping. Therefore, the current implmenetation makes no use of Matlab
%  subplot function, using axes instead. This can be problematic, as axis and subplot
%  parameters are quie different. Set isWrapper to "True" to return to wrapper mode, which
%  fully supports subplot format.
%
%% Input arguments (defaults exist):
%   margins- two elements vector [vertical,horizontal] defining the margins between
%        neighbouring axes. Default value is 0.04
%
%% Output arguments
%   same as subplot- none, or axes handle according to function call.
%
%% Issues & Comments
%  - Note that if additional elements are used in order to be passed to subplot, margins
%     parameter must be defined. For default margins value use empty element- [].
%  - 
%
%% Example
% close all;
% img=imread('peppers.png');
% figSubplotH=figure('Name', 'subplot');
% figSubplotTightH=figure('Name', 'subplot_tight');
% nElems=17;
% subplotRows=ceil(sqrt(nElems)-1);
% subplotRows=max(1, subplotRows);
% subplotCols=ceil(nElems/subplotRows);
% for iElem=1:nElems
%    figure(figSubplotH);
%    subplot(subplotRows, subplotCols, iElem);
%    imshow(img);
%    figure(figSubplotTightH);
%    subplot_tight(subplotRows, subplotCols, iElem, [0.0001]);
%    imshow(img);
% end
%
%% See also
%  - subplot
%
%% Revision history
% First version: Nikolay S. 2011-03-29.
% Last update:   Nikolay S. 2012-05-24.
%
% *List of Changes:*
% 2012-05-24
%  Non wrapping mode (based on axes command) added, to deal with an issue of disappearing
%     subplots occuring with massive axes.
%% Default params
isWrapper=false;
if (nargin<4) || isempty(margins)
    margins=[0.04,0.04]; % default margins value- 4% of figure
end
if length(margins)==1
    margins(2)=margins;
end
%note n and m are switched as Matlab indexing is column-wise, while subplot indexing is row-wise :(
[subplot_col,subplot_row]=ind2sub([n,m],p);  
height=(1-(m+1)*margins(1))/m; % single subplot height
width=(1-(n+1)*margins(2))/n;  % single subplot width
% note subplot suppors vector p inputs- so a merged subplot of higher dimentions will be created
subplot_cols=1+max(subplot_col)-min(subplot_col); % number of column elements in merged subplot 
subplot_rows=1+max(subplot_row)-min(subplot_row); % number of row elements in merged subplot   
merged_height=subplot_rows*( height+margins(1) )- margins(1);   % merged subplot height
merged_width= subplot_cols*( width +margins(2) )- margins(2);   % merged subplot width
merged_bottom=(m-max(subplot_row))*(height+margins(1)) +margins(1); % merged subplot bottom position
merged_left=min(subplot_col)*(width+margins(2))-width;              % merged subplot left position
pos=[merged_left, merged_bottom, merged_width, merged_height];
if isWrapper
   h=subplot(m, n, p, varargin{:}, 'Units', 'Normalized', 'Position', pos);
else
   h=axes('Position', pos, varargin{:});
end
if nargout==1
   vargout=h;
end
```
