## Supplementary material for "Instantaneous phase of rhythmic behaviour under volitional control": Code and data files: get_omega.m

```
function outFreq=get_omega(PHIc,sr,m,g)
% get median of input signal derivative 
% input:
%       signal: input signal
%       sr: sampling rate

%       m: (positive real; default=16): length of the Savitzky-Golay differentiator
%       g: (matrix of reals) coefficients of the differentiatior. If not
%       provided it is computed with polinomial order equal to 5
% output:
%       allFreqs: frequency values
%       Ts: vector of time steps

% Leonardo Lancia 14/10/2023
% mailto:

if nargin<3 || isempty(m)% initialize m
    m=16;
end
if nargin<4 || isempty(g)% initialize g
    g = SG_calc(m,5, 1)'; %compute coeffs for the sgolay differentiator
end


p=1; % index of the relevant S.G. coefficients
h=1/sr; % sampling period

myKernel=factorial(p)/(-h)^p * g(:,p+1); % cernel of the differentiator


PHIu=unwrap(PHIc); % unwrap phase
dPHI = filter(myKernel,1,PHIu); % differentiation
dPHI=-[dPHI(m/2+1:end)]./sr;% correct for filter delay without padding with zeros
outFreq=median(dPHI./(2*pi));%get median frequency
```
