## Supplementary material for "Instantaneous phase of rhythmic behaviour under volitional control": Code and data files: getPHImask.m

```
function [newPHI,IMF1,PHI,centeredSig,mask ]=getPHImask(signal,sr,m,n,nMasks,ampCoeff,quadMethod,threshs)
% input: 
% signal: input signal
% sr: (positive integer) sampling rate
% m: (positive integer; optional, default: 16) number of filtered points for Savitzky-Golay differentiator 
% n: (positive integer; optional, default: 5) polinomial order of the differentiator
% nMasks: (positive integer; optional, default 22)number of masks used to extract the independent mode function via masked EMD
% ampCoeff:(positive real; optional, default: 2) coefficient determining the amplitude of the masks as a
%            proportion of 4*std(signal), which is meant to be a rough estimate of the signal's 
%           range from the std if the signal's values are normally distributed.
% quadMethod: (a string or a cell of two strings, default: {'h','h'}). MEthod to be use in the 
%            computation of the quadrature signal 'h' stands for Hilbert and 'q'. 
%            If two strings are provided a different method will be adopted in in the first 
%            or the second part of the algorithm.
% threshs:(scalar or vector of two positive real values close to zero, default: [1E-10, 1E-10]) threshold for refined 
%          amplitude normalization. If to values, different thresholds will be used in the two parts 
%          of the algorithm.

% output:
% newPHI: instantaneous phase
% IMF1: output of masked sifting
% PHI: initial phase estimte
% centredSig: centred signal,
% mask: mask signal with inital phase set to 0

% Leonardo Lancia 14/10/2023
% mailto:

if diff(size(signal)) >0% be sure that the signal is a column vector
    signal=signal';
end
if nargin<3 || isempty(m)
    m=16;% number of filtered points (FIR order; keep it even if you do not want warnings).
end
if nargin<4 || isempty(n)
    n=5;% approximation polynomial order
end
if nargin<5 || isempty(nMasks)
    nMasks=22;
end
if nargin<6 || isempty(ampCoeff)
    ampCoeff=[];
end
if nargin<7 || isempty(quadMethod)
    quadMethod={'h','h'};
elseif iscell(quadMethod) && length(quadMethod)==1
    quadMethod={quadMethod{1},quadMethod{1}};
elseif ischar(quadMethod)
    quadMethod={quadMethod,quadMethod};
end
if nargin<8 || isempty(threshs)
    threshs=[NaN,NaN];
elseif length(threshs)==1
    threshs=[threshs,threshs];
end

origLen=length(signal); %original length of the signal

M = (m+1)/2;      % output point , M = (m+1)/2; % middle point for odd m
h=1/sr;
g = SG_calc(m,n, 1)'; %compute coeffs for the sgolay differentiator
p=1;

% centring (sifting with one iteration)
options.MAXITERATIONS=1;
options.MAXMODES=1;
options.FIX=1;
[centeredSig,~]=emd(signal,options);
centeredSig=centeredSig(1,:)';

normSig=normalize_cycle_amp(centeredSig,[],threshs(1),5);%refined amplitude normalization

if strcmpi(quadMethod{1},'q')
    PHI=quadAngle(normSig,0); % direct quadrature phase estimation 
elseif strcmpi(quadMethod{1},'hq')
    PHI=quadAngle(normSig,1); % hilbert quadrature phase estimation 
elseif strcmpi(quadMethod{1},'h')
    PHI=wrapTo2Pi(unwrap(angle(hilbert(normSig)))); % hilbert  estimation 
else
    error('quadMethod can only be ''q'',''h'' or ''hq''')
end

newPHI=PHI;
%estimation of local frequancies
sigFreq=get_omega(newPHI,sr,m,g);
t=[1:length(newPHI)]./sr; %time stamps of the phase signal
if ~isempty(ampCoeff)
    ampCoeff=ampCoeff.*4.*std(signal);
end
[IMF1,~,mask]=mask_sift(signal,sigFreq,[],nMasks,ampCoeff);%get signal via masked sifting
IMF=normalize_cycle_amp(IMF1,[],threshs(2),5);%refined amplitude normalizatin algo

if strcmpi(quadMethod{2},'q')
	newPHI=quadAngle(IMF,0); % direct quadrature phase estimation 
elseif strcmpi(quadMethod{2},'hq')
    newPHI=quadAngle(IMF,1); % hilbert quadrature phase estimation 
elseif strcmpi(quadMethod{2},'h')
    newPHI=wrapTo2Pi(unwrap(angle(hilbert(IMF)))); % hilbert  estimation 
else
    error('quadMethod can only be ''q'',''h'' or ''hq''')
end
```
