## Supplementary material for "Instantaneous phase of rhythmic behaviour under volitional control": Code and data files: mask_sift.m

```
function [IMF,maskOmega,mask]=mask_sift(x,maskOmega,frqEst,nPhases,maskAmpCoeff)

% performs masked sifting of the input signal x with oscillatory frequency
% maskOmega in radians per sample.

%input: 
%       x: input signal
%       maskOmega (positive real; optional, by default it is computed from the signal): 
%               mask signals frequency in radians per sample (optional).
%               If not provided, empty or equal to zero, the mask frequency  
%               is computed with the method defined by the third argument
%       frqEst (string among 'phi' and 'zc', optional, default 'phi')frequency estimation method . 
%               If 'phi' the phase is computed and the frequency is obtained from its median
%               derivative. Otherwise the frequency is computed as the reciprocal of the median 
%               distance between consecutive in time zero crossing points
%       nPhases (positive integer, optional, default = 22) number of masking signals used 
%       maskAmpCoeff (optional, default=2): coefficient determining the amplitude of the masks as a
%            proportion of 4*std(signal), which is meant to be a rough estimate of the signal's 
%            range from the std if the signal's values are normally distributed.
%
%output:
%      IMF: sifted signal
%      maskOmega: frequency of the mask signal
%      mask: mask signal with initial phase equal to zero 

% Leonardo Lancia 14/10/2023
% mailto:

if diff(size(x))>0 % be sure that the input signal is a column vector
    x=x';
end
if nargin<3 || isempty(frqEst) % initialize freq. estimation method
    frqEst='phi';
end

if nargin<4 || isempty(nPhases) % initialize num of mask signals
    nPhases=22;
end

if nargin<5 || isempty(maskAmpCoeff) % initialize mask amplitude
    maskAmpCoeff=2*4*std(x);
end

% compute mask frequency if required
if nargin<2 || isempty(maskOmega) || (length(maskOmega)==1 && maskOmega==0)
    % center signal via one sifting iteration
    options.MAXITERATIONS=1;
    options.MAXMODES=1;
    [IMF0,~]=emd(x,options);
    IMF0=IMF0(1,:)';
 
    IMF0= normalize_cycle_amp(IMF0);     % refined amplitude normalization
    if strcmp(frqEst,'phi')              % if frequency is to be estimated via phase derivative
        IPH=angle(hilbert(IMF0));        % phase
        Ifreq=diff(unwrap(IPH));         
        IPH(Ifreq<0)=NaN;
        maskOmega=nanmean(diff(unwrap(IPH))/(2*pi));%normalized freq      
    elseif strcmp(frqEst,'zc')% if frequency is to be estimated via reciprocal of zero-crossing distance
         zci = @(v) find(v(:).*circshift(v(:), [-1 0]) <= 0);
         maskOmega=ceil(length(zci(IMF0))/2)/length(IMF0);%normalized freq
    end
end

phases=linspace(0,2*pi,nPhases+1); % initial phase values 
phases=phases(1:end-1);
if length(maskOmega)==1 % if constant max frequency
    masks=maskAmpCoeff.*sin([0:length(x)-1].*maskOmega*2*pi+phases')'; % synthetize nask synewave
else
    error('maskOmega must be a scalar real value')
end
options.MAXITERATIONS=10;% parametrize EMD function to do max 10 sifting iteration
options.MAXMODES=1;
tmpIMFS=zeros(size(masks)); % build matrix to store partial IMFs
for i=1:size(masks,2)
    tmpSig=x+masks(:,i); % add mask to input signal
    [thisImf,~]=emd(tmpSig,options);% do sifting
    tmpIMFS(:,i)=thisImf(1,:)'-masks(:,i); % remove mask
end
IMF=sum(tmpIMFS,2)./nPhases;% compute average IMF
mask=masks(:,1); % get output mask 
```
