## Supplementary material for "Instantaneous phase of rhythmic behaviour under volitional control": Code and data files: maskEMD.m

```
function [outIMFs,IMFfreqs]=maskEMD(signal,sr, maxIMFn,nMasks,ampCoeff,m,n,stdRatioThresh)
% performs EMD via Masked Sifting

% input:

% signal: input signal
% sr: (positive real): sampling frequency in Hz
% maxIMFn: maxIMFn (positive integer; optional, default=10): maximum number of IMFs.
%         If equal to 0, the procedure will be stopped when the obtained 
%         IMF contains no extrema.
% nMasks (positive integer; optional, default= 22 ): number of mask signal used in sifting 
% ampCoeff (positive real; optional, default=2): coefficient determining the amplitude of the masks as a
%            proportion of 4*std(signal), which is meant to be a rough estimate of the signal's 
%            range from the std if the signal's values are normally distributed.
% m (positive integer; optional, default= 16): length of the Savitzky-Golay differentiator.  
% n (positive integer; optional, default= 5): order of the Savitzky-Golay differentiator.  
% stdRatioThresh (positive real; optional, default= 1E-6): threshold for EMD convergence

% 
% output:
% outIMFs: the extracted IMFs
% IMFfreqs: the estimated frequencies of the extracted IMFs 
% Leonardo Lancia 14/10/2023
% mailto:

if nargin<3 || isempty(maxIMFn) || isnan(maxIMFn) || maxIMFn==0 %initialize maxIMF
    maxIMFn=Inf;
end
if nargin<4 || isempty(nMasks) || isnan(nMasks)  || nMasks==0 %initialize nMasks
    nMasks=22;
end
if nargin<5 || isempty(ampCoeff) || isnan(ampCoeff)  || ampCoeff==0 %initialize ampCoeff
    ampCoeff=[];
end
if nargin<6 || isempty(m) || isnan(m)  || m==0 %initialize m
    m=16;
end
if nargin<7 || isempty(n) || isnan(n)  || n==0 %initialize n
    n=5;
end
if nargin<8 || isempty(stdRatioThresh)
    stdRatioThresh=1e-6;
end
SG_coef = SG_calc(m,n,1)'; %compute coeffs for the sgolay differentiator

nPks=1; %initialize variable containing the unmber of extrema in the current IMF
resid=signal;  %initialize input signal for masked sifting 
outIMFs=zeros(length(signal),maxIMFn);  %initialize matrix containing IMFs
IMFfreqs=zeros(maxIMFn,1); %initialize vector containing IMFs frequencies
nn=0; % number of iterations 
options.MAXITERATIONS=1;% set sifting iterations number
options.MAXMODES=1; % set number of mode to extract when calling the emd function 
stdRatio=1;
while nPks>0 && nn < maxIMFn && stdRatio>stdRatioThresh
    nn =nn+1; %update num of iteration
    
    [myMode,~]=emd(resid,options); % center signal via one iteration of sifting
    myMode=myMode(1,:)';

    if ~isempty(ampCoeff) && nn==1
        thisCoeff=ampCoeff.*4*std(myMode); % set amplitude of mask signal (it is based on the first IMF for denoising purposes)
    
    elseif isempty(ampCoeff)
        thisCoeff=[]; % the empty values triggers the use of the default value in mask_sift
                      % this corresponds to twice the range of the input signal
    end
    
    myModeN=normalize_cycle_amp(myMode); % apply refined amplitude normalization
    
    PHIc=angle(hilbert(myModeN));%hilbert phase (the first estimate is used to infer local frequancy)

    sigFreq=get_omega(PHIc,sr,m,SG_coef);
    
    [outIMFs(:,nn),~]=mask_sift(resid,sigFreq,[],nMasks,thisCoeff); % do masked sifting
    warning off % remove warnings due to NaN ignored during interpolation 
    myModeN=normalize_cycle_amp(outIMFs(:,nn),[],[],10); % apply refined amplitude normalization 
    warning on
    PHINew=angle(hilbert(myModeN));%hilbert phase (better suited than quadr. to infer local frequancy)
  
    PHIu=unwrap(PHINew); % unwrap phase
    medF=median((diff(PHIu)/(2*pi))); % compute median frequency

    IMFfreqs(nn)=medF; % store median frequency
    thisStd=std(resid);
    resid=resid-outIMFs(:,nn); % update input signal for next application of masked sifting 
    nPks=length(findpeaks(resid)); % count number of peaks
    nPks=nPks+length(findpeaks(-resid)); % get number of extrema
    stdRatio=std(resid)/thisStd;
    if ~isempty(ampCoeff) && nn==1
        thisCoeff=ampCoeff.*4.*std(outIMFs(:,nn));
    end
end

outIMFs=outIMFs(:,1:nn);
IMFfreqs=IMFfreqs(1:nn);
```
