## Supplementary material for "Instantaneous phase of rhythmic behaviour under volitional control": Code and data files: mEMDdenoise.m

```
function [filtered,imf,imfF,filteredidx,noiseStd]=mEMDdeNoise(data,sr,nMasks,ampCoeff,alpha,nReps,m,n)
% remove noise from signal in Data sampled at freq. sr 
% input:
%
% data: input signal
% sr: sampling rate
% nMasks(positive integer; optional, default: 8): number of mask signal for masked EMD application .  
% ampCoeff (positive real; optional, default: 2): amplitude gain of the mask signals 
%       (it is multiplied by 4sd(IMF1), where IMF1 is the first IMF or the centered data at the first iteration ).  
% nReps(positive integer; optional, default: 100): number of repetitions of simulated random processes .  
% m (positive integer; optional, default: 16): length of the Savitzky-Golay differentiator.  
% n (positive integer; optional, default: 5): order of the Savitzky-Golay differentiator.  

%
% output:
%       filtered: filtered signal
%       imf: all signal's IMFs
%       imfF: frequencies of extracted IMFs
%       filteredidx: indexes of IMFs composing the filtered signal
%       noiseStd: standard deviation of the estimated noise (sum of the random components).

% Leonardo Lancia 14/10/2023
% mailto:

if diff(size(data))>0 % be sure that the input signal is a column vector
    data=data';
end
data=zscore(data);

% apply masked EMD  to get the signal's IMFs
[imf,imfF]=maskEMD(data,sr,10,nMasks,ampCoeff,m,n);

He = hurst(zscore(imf(:,1)'));% compute Hurst exponenet of first IMF

nObs = size(data, 1); % signal length
nImf = size(imf, 2); % number of IMFs

energy = zeros(nImf, 1);% build storage for IMFs' energy values

NoiseStd=std(data); %estimate standard deviation of the signal
% produce nReps random signals with Hurst exponent set He and std set to NoiseStd
f = ffgn(1,max([abs(realmin),He]),nReps,length(imf(:,1)),1);
noiseLev=nan(nReps,10); % build storage for energy of the IMFs extracted from the random signals

% N=size(imf,1);
options.MAXITERATIONS=10;
options.MAXMODES=10;% set maximum number of modes
for i =1:length(noiseLev)
   [rndImfs,~]=emd(f(i,:),options); % apply EMD
   %   [rndImfs,NB_ITERATIONS]=emdc_fix([],f(i,:),[],10); %if Rilling's c
   %        implementation avialable, things get much faster if this line
   %        substitutes the preceding one
   rndImfs=rndImfs';
   noiseLev(i,1:size(rndImfs,2))=sum(rndImfs.^2, 1)./nObs;% compute energy of ectracted IMFs
end
myQuants=quantile(noiseLev, [alpha, 1-alpha]);% compute quantiles of energy distributions

for i=1:nImf % compute the energy of each IMF extraced from real data
    energy(i, 1) = sum(imf(:, i).^2, 1)/nObs;
end
filteredidx = zeros(1, nImf);% build storage for chosen IMFs' indexes
filtered = zeros(nObs, 1);% initialize filtered signal

for i=2:nImf

    if(energy(i, 1) >= myQuants(2,i))% if test passed 
        filteredidx(1, i) = i; % store the index of the selected mode
        filtered = filtered + imf(:, i); % add the selected mode to the fitlered signal
    end
end
noiseStd=std(data-filtered); % compute std of the random component

% indexes of statistically significant IMFs
filteredidx = nonzeros(filteredidx)';
```
