## Supplementary material for "Instantaneous phase of rhythmic behaviour under volitional control": Code and data files: normalize_cycle_amp.m

```
function sigIn=normalize_cycle_amp(sigIn,symmetric,threshNorm,maxIterN, symN)
%    demodulate envelope peaks iteratively and using pchip interpolation.
%    At each iteration, max and min are individuated, their absolute values are 
%    interpolated to obtain an envelope. The signal is then divided by the envelope.
%    If even one peak of the obtained signal is larger than 0, this is sent to a new iteration.
%    Otherwise the procedure is stopped. Boundary conditions of the local extrema sequence 
%    are computed following Rilling 2007 (see function in '.\utils\boundary_conditions.m').
%
%    input:
%    
%    symmetric (binary, optional): if it is 0 the signal's cycles are not centered around 0. 
%           This is solved by applying one iteration of sifting.
%    threshNorm (positive real, optional): threshold to determine if peaks are different from 1/-1
%    maxIterN (positive integer, optional): maximum number of iterations
%    symN (positive integer, optional): number of extrtema to be added at the beginning and at the
%           end to regularize boundary conditions before interpolation.
%
%    output:
%     
%    normalized envelope
%    sigIn=copy.copy(sig)

% Leonardo Lancia 14/10/2023
% mailto:

if diff(size(sigIn))>0 % be sure that the input signal is a column vector
    sigIn=sigIn';
end

if nargin <2 || isempty(symmetric)
    symmetric=1;
end
if nargin <3 || isempty(threshNorm) || isnan(threshNorm)
    threshNorm=1e-10;
end
if nargin<4 || isempty(maxIterN)
    maxIterN=5;
end

if nargin<5 || isempty(symN)
    symN=2;
end
if symmetric==0 %if the signal is not centred on 0apply centring (one sifting iteration)
    options.MAXITERATIONS=1;
    options.MAXMODES=1;
    [sigIn,~]=emd(sigIn,options);
    sigIn=sigIn(1,:)';
end


newIdxs=[1:length(sigIn)]'; %input signal's time steps 

iter_val=1; %intitialize error value
niter=0; % current iteration number
while niter<maxIterN && iter_val>threshNorm
    niter=niter+1; %update iteration number
    
    [~,pksLoc]=findpeaks(sigIn); % find local maxima
    [~,valLoc]=findpeaks(-sigIn); %find local minima
    if length(pksLoc)+length(valLoc)<3 %if less than three extrema break and return
        sigIn=sigIn/range(sigIn);
        return 
    end    
    %set boundary conditions via Rilling's function
    [tMin,tMax,sigMin,sigMax] = boundary_conditions(valLoc',pksLoc',newIdxs',sigIn',sigIn',symN);
    %find local extrema order and sort their locations
    [tExtr,extrOrd]=sort([tMin,tMax]);
    allExtr=[sigMin,sigMax];
    allExtr=abs(allExtr(extrOrd));%sort local extrema absolute values according ot their order of appearence

    refSig= interp1(tExtr,allExtr,newIdxs, 'pchip'); %interpolate local extrema absolute values

	iter_val = abs(sum(refSig) - max(size(refSig))) ; % compute mean difference between envelopes values from last two iterations
    sigIn=sigIn./refSig;% normalize signal's amplitude

end
```
