## Supplementary material for "Instantaneous phase of rhythmic behaviour under volitional control": Code and data files: boundary_conditions.m

```
function [tmin,tmax,zmin,zmax] = boundary_conditions(indmin,indmax,t,x,z,nbsym)
% this is just a cut and paste of the subfunction boundary_conditions from 'emd.m' by g. Rilling	
% 
	lx = length(x);
	
	if (length(indmin) + length(indmax) < 3)
		error('not enough extrema')
	end

    % boundary conditions for interpolations :

	if indmax(1) < indmin(1)
    	if x(1) > x(indmin(1))
			lmax = fliplr(indmax(2:min(end,nbsym+1)));
			lmin = fliplr(indmin(1:min(end,nbsym)));
			lsym = indmax(1);
		else
			lmax = fliplr(indmax(1:min(end,nbsym)));
			lmin = [fliplr(indmin(1:min(end,nbsym-1))),1];
			lsym = 1;
		end
	else

		if x(1) < x(indmax(1))
			lmax = fliplr(indmax(1:min(end,nbsym)));
			lmin = fliplr(indmin(2:min(end,nbsym+1)));
			lsym = indmin(1);
		else
			lmax = [fliplr(indmax(1:min(end,nbsym-1))),1];
			lmin = fliplr(indmin(1:min(end,nbsym)));
			lsym = 1;
		end
	end
    
	if indmax(end) < indmin(end)
		if x(end) < x(indmax(end))
			rmax = fliplr(indmax(max(end-nbsym+1,1):end));
			rmin = fliplr(indmin(max(end-nbsym,1):end-1));
			rsym = indmin(end);
		else
			rmax = [lx,fliplr(indmax(max(end-nbsym+2,1):end))];
			rmin = fliplr(indmin(max(end-nbsym+1,1):end));
			rsym = lx;
		end
	else
		if x(end) > x(indmin(end))
			rmax = fliplr(indmax(max(end-nbsym,1):end-1));
			rmin = fliplr(indmin(max(end-nbsym+1,1):end));
			rsym = indmax(end);
		else
			rmax = fliplr(indmax(max(end-nbsym+1,1):end));
			rmin = [lx,fliplr(indmin(max(end-nbsym+2,1):end))];
			rsym = lx;
		end
	end
    
	tlmin = 2*t(lsym)-t(lmin);
	tlmax = 2*t(lsym)-t(lmax);
	trmin = 2*t(rsym)-t(rmin);
	trmax = 2*t(rsym)-t(rmax);
    
	% in case symmetrized parts do not extend enough
	if tlmin(1) > t(1) || tlmax(1) > t(1)
		if lsym == indmax(1)
			lmax = fliplr(indmax(1:min(end,nbsym)));
		else
			lmin = fliplr(indmin(1:min(end,nbsym)));
		end
		if lsym == 1
			error('bug')
		end
		lsym = 1;
		tlmin = 2*t(lsym)-t(lmin);
		tlmax = 2*t(lsym)-t(lmax);
	end   
    
	if trmin(end) < t(lx) || trmax(end) < t(lx)
		if rsym == indmax(end)
			rmax = fliplr(indmax(max(end-nbsym+1,1):end));
		else
			rmin = fliplr(indmin(max(end-nbsym+1,1):end));
		end
	if rsym == lx
		error('bug')
	end
		rsym = lx;
		trmin = 2*t(rsym)-t(rmin);
		trmax = 2*t(rsym)-t(rmax);
	end 
          
	zlmax =z(lmax); 
	zlmin =z(lmin);
	zrmax =z(rmax); 
	zrmin =z(rmin);
     
	tmin = [tlmin t(indmin) trmin];
	tmax = [tlmax t(indmax) trmax];
	zmin = [zlmin z(indmin) zrmin];
	zmax = [zlmax z(indmax) zrmax];
end
```
