## Supplementary material for "Instantaneous phase of rhythmic behaviour under volitional control": Code and data files: emd.m

```
%EMD  computes Empirical Mode Decomposition
%
%
%   Syntax
%
%
% IMF = EMD(X)
% IMF = EMD(X,...,'Option_name',Option_value,...)
% IMF = EMD(X,OPTS)
% [IMF,ORT,NB_ITERATIONS] = EMD(...)
%
%
%   Description
%
%
% IMF = EMD(X) where X is a real vector computes the Empirical Mode
% Decomposition [1] of X, resulting in a matrix IMF containing 1 IMF per row, the
% last one being the residue. The default stopping criterion is the one proposed
% in [2]:
%
%   at each point, mean_amplitude < THRESHOLD2*envelope_amplitude
%   &
%   mean of boolean array {(mean_amplitude)/(envelope_amplitude) > THRESHOLD} < TOLERANCE
%   &
%   |#zeros-#extrema|<=1
%
% where mean_amplitude = abs(envelope_max+envelope_min)/2
% and envelope_amplitude = abs(envelope_max-envelope_min)/2
% 
% IMF = EMD(X) where X is a complex vector computes Bivariate Empirical Mode
% Decomposition [3] of X, resulting in a matrix IMF containing 1 IMF per row, the
% last one being the residue. The default stopping criterion is similar to the
% one proposed in [2]:
%
%   at each point, mean_amplitude < THRESHOLD2*envelope_amplitude
%   &
%   mean of boolean array {(mean_amplitude)/(envelope_amplitude) > THRESHOLD} < TOLERANCE
%
% where mean_amplitude and envelope_amplitude have definitions similar to the
% real case
%
% IMF = EMD(X,...,'Option_name',Option_value,...) sets options Option_name to
% the specified Option_value (see Options)
%
% IMF = EMD(X,OPTS) is equivalent to the above syntax provided OPTS is a struct 
% object with field names corresponding to option names and field values being the 
% associated values 
%
% [IMF,ORT,NB_ITERATIONS] = EMD(...) returns an index of orthogonality
%                       ________
%         _  |IMF(i,:).*IMF(j,:)|
%   ORT = \ _____________________
%         /
%         �        || X ||�
%        i~=j
%
% and the number of iterations to extract each mode in NB_ITERATIONS
%
%
%   Options
%
%
%  stopping criterion options:
%
% STOP: vector of stopping parameters [THRESHOLD,THRESHOLD2,TOLERANCE]
% if the input vector's length is less than 3, only the first parameters are
% set, the remaining ones taking default values.
% default: [0.05,0.5,0.05]
%
% FIX (int): disable the default stopping criterion and do exactly <FIX> 
% number of sifting iterations for each mode
%
% FIX_H (int): disable the default stopping criterion and do <FIX_H> sifting 
% iterations with |#zeros-#extrema|<=1 to stop [4]
%
%  bivariate/complex EMD options:
%
% COMPLEX_VERSION: selects the algorithm used for complex EMD ([3])
% COMPLEX_VERSION = 1: "algorithm 1"
% COMPLEX_VERSION = 2: "algorithm 2" (default)
% 
% NDIRS: number of directions in which envelopes are computed (default 4)
% rem: the actual number of directions (according to [3]) is 2*NDIRS
% 
%  other options:
%
% T: sampling times (line vector) (default: 1:length(x))
%
% MAXITERATIONS: maximum number of sifting iterations for the computation of each
% mode (default: 2000)
%
% MAXMODES: maximum number of imfs extracted (default: Inf)
%
% DISPLAY: if equals to 1 shows sifting steps with pause
% if equals to 2 shows sifting steps without pause (movie style)
% rem: display is disabled when the input is complex
%
% INTERP: interpolation scheme: 'linear', 'cubic', 'pchip' or 'spline' (default)
% see interp1 documentation for details
%
% MASK: masking signal used to improve the decomposition according to [5]
%
%
%   Examples
%
%
%X = rand(1,512);
%
%IMF = emd(X);
%
%IMF = emd(X,'STOP',[0.1,0.5,0.05],'MAXITERATIONS',100);
%
%T=linspace(0,20,1e3);
%X = 2*exp(i*T)+exp(3*i*T)+.5*T;
%IMF = emd(X,'T',T);
%
%OPTIONS.DISLPAY = 1;
%OPTIONS.FIX = 10;
%OPTIONS.MAXMODES = 3;
%[IMF,ORT,NBITS] = emd(X,OPTIONS);
%
%
%   References
%
%
% [1] N. E. Huang et al., "The empirical mode decomposition and the
% Hilbert spectrum for non-linear and non stationary time series analysis",
% Proc. Royal Soc. London A, Vol. 454, pp. 903-995, 1998
%
% [2] G. Rilling, P. Flandrin and P. Gon�alves
% "On Empirical Mode Decomposition and its algorithms",
% IEEE-EURASIP Workshop on Nonlinear Signal and Image Processing
% NSIP-03, Grado (I), June 2003
%
% [3] G. Rilling, P. Flandrin, P. Gon�alves and J. M. Lilly.,
% "Bivariate Empirical Mode Decomposition",
% Signal Processing Letters (submitted)
%
% [4] N. E. Huang et al., "A confidence limit for the Empirical Mode
% Decomposition and Hilbert spectral analysis",
% Proc. Royal Soc. London A, Vol. 459, pp. 2317-2345, 2003
%
% [5] R. Deering and J. F. Kaiser, "The use of a masking signal to improve 
% empirical mode decomposition", ICASSP 2005
%
%
% See also
%  emd_visu (visualization),
%  emdc, emdc_fix (fast implementations of EMD),
%  cemdc, cemdc_fix, cemdc2, cemdc2_fix (fast implementations of bivariate EMD),
%  hhspectrum (Hilbert-Huang spectrum)
%
%
% G. Rilling, last modification: 3.2007
%


function [imf,ort,nbits] = emd(varargin)

[x,t,sd,sd2,tol,MODE_COMPLEX,ndirs,display_sifting,sdt,sd2t,r,imf,k,nbit,NbIt,MAXITERATIONS,FIXE,FIXE_H,MAXMODES,INTERP,mask] = init(varargin{:});

if display_sifting
  fig_h = figure;
end


%main loop : requires at least 3 extrema to proceed
while ~stop_EMD(r,MODE_COMPLEX,ndirs) && (k < MAXMODES+1 || MAXMODES == 0) && ~any(mask)

  % current mode
  m = r;

  % mode at previous iteration
  mp = m;

  %computation of mean and stopping criterion
  if FIXE
    [stop_sift,moyenne] = stop_sifting_fixe(t,m,INTERP,MODE_COMPLEX,ndirs);
  elseif FIXE_H
    stop_count = 0;
    [stop_sift,moyenne] = stop_sifting_fixe_h(t,m,INTERP,stop_count,FIXE_H,MODE_COMPLEX,ndirs);
  else
    [stop_sift,moyenne] = stop_sifting(m,t,sd,sd2,tol,INTERP,MODE_COMPLEX,ndirs);
  end

  % in case the current mode is so small that machine precision can cause
  % spurious extrema to appear
  if (max(abs(m))) < (1e-10)*(max(abs(x)))
    if ~stop_sift
      warning('emd:warning','forced stop of EMD : too small amplitude')
    else
      disp('forced stop of EMD : too small amplitude')
    end
    break
  end


  % sifting loop
  while ~stop_sift && nbit<MAXITERATIONS

    if(~MODE_COMPLEX && nbit>MAXITERATIONS/5 && mod(nbit,floor(MAXITERATIONS/10))==0 && ~FIXE && nbit > 100)
      disp(['mode ',int2str(k),', iteration ',int2str(nbit)])
      if exist('s','var')
        disp(['stop parameter mean value : ',num2str(s)])
      end
      [im,iM] = extr(m);
      disp([int2str(sum(m(im) > 0)),' minima > 0; ',int2str(sum(m(iM) < 0)),' maxima < 0.'])
    end

    %sifting
    m = m - moyenne;

    %computation of mean and stopping criterion
    if FIXE
      [stop_sift,moyenne] = stop_sifting_fixe(t,m,INTERP,MODE_COMPLEX,ndirs);
    elseif FIXE_H
      [stop_sift,moyenne,stop_count] = stop_sifting_fixe_h(t,m,INTERP,stop_count,FIXE_H,MODE_COMPLEX,ndirs);
    else
      [stop_sift,moyenne,s] = stop_sifting(m,t,sd,sd2,tol,INTERP,MODE_COMPLEX,ndirs);
    end

    % display
    if display_sifting && ~MODE_COMPLEX
      NBSYM = 2;
      [indmin,indmax] = extr(mp);
      [tmin,tmax,mmin,mmax] = boundary_conditions(indmin,indmax,t,mp,mp,NBSYM);
      envminp = interp1(tmin,mmin,t,INTERP);
      envmaxp = interp1(tmax,mmax,t,INTERP);
      envmoyp = (envminp+envmaxp)/2;
      if FIXE || FIXE_H
        display_emd_fixe(t,m,mp,r,envminp,envmaxp,envmoyp,nbit,k,display_sifting)
      else
        sxp=2*(abs(envmoyp))./(abs(envmaxp-envminp));
        sp = mean(sxp);
        display_emd(t,m,mp,r,envminp,envmaxp,envmoyp,s,sp,sxp,sdt,sd2t,nbit,k,display_sifting,stop_sift)
      end
    end

    mp = m;
    nbit=nbit+1;
    NbIt=NbIt+1;

    if(nbit==(MAXITERATIONS-1) && ~FIXE && nbit > 100)
      if exist('s','var')
        warning('emd:warning',['forced stop of sifting : too many iterations... mode ',int2str(k),'. stop parameter mean value : ',num2str(s)])
      else
        warning('emd:warning',['forced stop of sifting : too many iterations... mode ',int2str(k),'.'])
      end
    end

  end % sifting loop
  imf(k,:) = m;
  if display_sifting
    disp(['mode ',int2str(k),' stored'])
  end
  nbits(k) = nbit;
  k = k+1;


  r = r - m;
  nbit=0;


end %main loop

if any(r) && ~any(mask)
  imf(k,:) = r;
end

ort = io(x,imf);

if display_sifting
  close
end
end

%---------------------------------------------------------------------------------------------------
% tests if there are enough (3) extrema to continue the decomposition
function stop = stop_EMD(r,MODE_COMPLEX,ndirs)
if MODE_COMPLEX
  for k = 1:ndirs
    phi = (k-1)*pi/ndirs;
    [indmin,indmax] = extr(real(exp(i*phi)*r));
    ner(k) = length(indmin) + length(indmax);
  end
  stop = any(ner < 3);
else
  [indmin,indmax] = extr(r);
  ner = length(indmin) + length(indmax);
  stop = ner < 3;
end
end

%---------------------------------------------------------------------------------------------------
% computes the mean of the envelopes and the mode amplitude estimate
function [envmoy,nem,nzm,amp] = mean_and_amplitude(m,t,INTERP,MODE_COMPLEX,ndirs)
NBSYM = 2;
if MODE_COMPLEX
  switch MODE_COMPLEX
    case 1
      for k = 1:ndirs
        phi = (k-1)*pi/ndirs;
        y = real(exp(-i*phi)*m);
        [indmin,indmax,indzer] = extr(y);
        nem(k) = length(indmin)+length(indmax);
        nzm(k) = length(indzer);
        [tmin,tmax,zmin,zmax] = boundary_conditions(indmin,indmax,t,y,m,NBSYM);
        envmin(k,:) = interp1(tmin,zmin,t,INTERP);
        envmax(k,:) = interp1(tmax,zmax,t,INTERP);
      end
      envmoy = mean((envmin+envmax)/2,1);
      if nargout > 3
        amp = mean(abs(envmax-envmin),1)/2;
      end
    case 2
      for k = 1:ndirs
        phi = (k-1)*pi/ndirs;
        y = real(exp(-i*phi)*m);
        [indmin,indmax,indzer] = extr(y);
        nem(k) = length(indmin)+length(indmax);
        nzm(k) = length(indzer);
        [tmin,tmax,zmin,zmax] = boundary_conditions(indmin,indmax,t,y,y,NBSYM);
        envmin(k,:) = exp(i*phi)*interp1(tmin,zmin,t,INTERP);
        envmax(k,:) = exp(i*phi)*interp1(tmax,zmax,t,INTERP);
      end
      envmoy = mean((envmin+envmax),1);
      if nargout > 3
        amp = mean(abs(envmax-envmin),1)/2;
      end
  end
else
  [indmin,indmax,indzer] = extr(m);
  nem = length(indmin)+length(indmax);
  nzm = length(indzer);
  [tmin,tmax,mmin,mmax] = boundary_conditions(indmin,indmax,t,m,m,NBSYM);
  envmin = interp1(tmin,mmin,t,INTERP);
  envmax = interp1(tmax,mmax,t,INTERP);
  envmoy = (envmin+envmax)/2;
  if nargout > 3
    amp = mean(abs(envmax-envmin),1)/2;
  end
end
end

%-------------------------------------------------------------------------------
% default stopping criterion
function [stop,envmoy,s] = stop_sifting(m,t,sd,sd2,tol,INTERP,MODE_COMPLEX,ndirs)
try
  [envmoy,nem,nzm,amp] = mean_and_amplitude(m,t,INTERP,MODE_COMPLEX,ndirs);
  sx = abs(envmoy)./amp;
  s = mean(sx);
  stop = ~((mean(sx > sd) > tol | any(sx > sd2)) & (all(nem > 2)));
  if ~MODE_COMPLEX
    stop = stop && ~(abs(nzm-nem)>1);
  end
catch
  stop = 1;
  envmoy = zeros(1,length(m));
  s = NaN;
end
end

%-------------------------------------------------------------------------------
% stopping criterion corresponding to option FIX
function [stop,moyenne]= stop_sifting_fixe(t,m,INTERP,MODE_COMPLEX,ndirs)
try
  moyenne = mean_and_amplitude(m,t,INTERP,MODE_COMPLEX,ndirs);
  stop = 0;
catch
  moyenne = zeros(1,length(m));
  stop = 1;
end
end

%-------------------------------------------------------------------------------
% stopping criterion corresponding to option FIX_H
function [stop,moyenne,stop_count]= stop_sifting_fixe_h(t,m,INTERP,stop_count,FIXE_H,MODE_COMPLEX,ndirs)
try
  [moyenne,nem,nzm] = mean_and_amplitude(m,t,INTERP,MODE_COMPLEX,ndirs);
  if (all(abs(nzm-nem)>1))
    stop = 0;
    stop_count = 0;
  else
    stop_count = stop_count+1;
    stop = (stop_count == FIXE_H);
  end
catch
  moyenne = zeros(1,length(m));
  stop = 1;
end
end

%-------------------------------------------------------------------------------
% displays the progression of the decomposition with the default stopping criterion
function display_emd(t,m,mp,r,envmin,envmax,envmoy,s,sb,sx,sdt,sd2t,nbit,k,display_sifting,stop_sift)
subplot(4,1,1)
plot(t,mp);hold on;
plot(t,envmax,'--k');plot(t,envmin,'--k');plot(t,envmoy,'r');
title(['IMF ',int2str(k),';   iteration ',int2str(nbit),' before sifting']);
set(gca,'XTick',[])
hold  off
subplot(4,1,2)
plot(t,sx)
hold on
plot(t,sdt,'--r')
plot(t,sd2t,':k')
title('stop parameter')
set(gca,'XTick',[])
hold off
subplot(4,1,3)
plot(t,m)
title(['IMF ',int2str(k),';   iteration ',int2str(nbit),' after sifting']);
set(gca,'XTick',[])
subplot(4,1,4);
plot(t,r-m)
title('residue');
disp(['stop parameter mean value : ',num2str(sb),' before sifting and ',num2str(s),' after'])
if stop_sift
  disp('last iteration for this mode')
end
if display_sifting == 2
  pause(0.01)
else
  pause
end
end

%---------------------------------------------------------------------------------------------------
% displays the progression of the decomposition with the FIX and FIX_H stopping criteria
function display_emd_fixe(t,m,mp,r,envmin,envmax,envmoy,nbit,k,display_sifting)
subplot(3,1,1)
plot(t,mp);hold on;
plot(t,envmax,'--k');plot(t,envmin,'--k');plot(t,envmoy,'r');
title(['IMF ',int2str(k),';   iteration ',int2str(nbit),' before sifting']);
set(gca,'XTick',[])
hold  off
subplot(3,1,2)
plot(t,m)
title(['IMF ',int2str(k),';   iteration ',int2str(nbit),' after sifting']);
set(gca,'XTick',[])
subplot(3,1,3);
plot(t,r-m)
title('residue');
if display_sifting == 2
  pause(0.01)
else
  pause
end
end

%---------------------------------------------------------------------------------------
% defines new extrema points to extend the interpolations at the edges of the
% signal (mainly mirror symmetry)
function [tmin,tmax,zmin,zmax] = boundary_conditions(indmin,indmax,t,x,z,nbsym)
	
	lx = length(x);
	
	if (length(indmin) + length(indmax) < 3)
		error('not enough extrema')
	end

    % boundary conditions for interpolations :

	if indmax(1) < indmin(1)
    	if x(1) > x(indmin(1))
			lmax = fliplr(indmax(2:min(end,nbsym+1)));
			lmin = fliplr(indmin(1:min(end,nbsym)));
			lsym = indmax(1);
		else
			lmax = fliplr(indmax(1:min(end,nbsym)));
			lmin = [fliplr(indmin(1:min(end,nbsym-1))),1];
			lsym = 1;
		end
	else

		if x(1) < x(indmax(1))
			lmax = fliplr(indmax(1:min(end,nbsym)));
			lmin = fliplr(indmin(2:min(end,nbsym+1)));
			lsym = indmin(1);
		else
			lmax = [fliplr(indmax(1:min(end,nbsym-1))),1];
			lmin = fliplr(indmin(1:min(end,nbsym)));
			lsym = 1;
		end
	end
    
	if indmax(end) < indmin(end)
		if x(end) < x(indmax(end))
			rmax = fliplr(indmax(max(end-nbsym+1,1):end));
			rmin = fliplr(indmin(max(end-nbsym,1):end-1));
			rsym = indmin(end);
		else
			rmax = [lx,fliplr(indmax(max(end-nbsym+2,1):end))];
			rmin = fliplr(indmin(max(end-nbsym+1,1):end));
			rsym = lx;
		end
	else
		if x(end) > x(indmin(end))
			rmax = fliplr(indmax(max(end-nbsym,1):end-1));
			rmin = fliplr(indmin(max(end-nbsym+1,1):end));
			rsym = indmax(end);
		else
			rmax = fliplr(indmax(max(end-nbsym+1,1):end));
			rmin = [lx,fliplr(indmin(max(end-nbsym+2,1):end))];
			rsym = lx;
		end
	end
    
	tlmin = 2*t(lsym)-t(lmin);
	tlmax = 2*t(lsym)-t(lmax);
	trmin = 2*t(rsym)-t(rmin);
	trmax = 2*t(rsym)-t(rmax);
    
	% in case symmetrized parts do not extend enough
	if tlmin(1) > t(1) || tlmax(1) > t(1)
		if lsym == indmax(1)
			lmax = fliplr(indmax(1:min(end,nbsym)));
		else
			lmin = fliplr(indmin(1:min(end,nbsym)));
		end
		if lsym == 1
			error('bug')
		end
		lsym = 1;
		tlmin = 2*t(lsym)-t(lmin);
		tlmax = 2*t(lsym)-t(lmax);
	end   
    
	if trmin(end) < t(lx) || trmax(end) < t(lx)
		if rsym == indmax(end)
			rmax = fliplr(indmax(max(end-nbsym+1,1):end));
		else
			rmin = fliplr(indmin(max(end-nbsym+1,1):end));
		end
	if rsym == lx
		error('bug')
	end
		rsym = lx;
		trmin = 2*t(rsym)-t(rmin);
		trmax = 2*t(rsym)-t(rmax);
	end 
          
	zlmax =z(lmax); 
	zlmin =z(lmin);
	zrmax =z(rmax); 
	zrmin =z(rmin);
     
	tmin = [tlmin t(indmin) trmin];
	tmax = [tlmax t(indmax) trmax];
	zmin = [zlmin z(indmin) zrmin];
	zmax = [zlmax z(indmax) zrmax];
end
    
%---------------------------------------------------------------------------------------------------
%extracts the indices of extrema
function [indmin, indmax, indzer] = extr(x,t)

if(nargin==1)
  t=1:length(x);
end

m = length(x);

if nargout > 2
  x1=x(1:m-1);
  x2=x(2:m);
  indzer = find(x1.*x2<0);

  if any(x == 0)
    iz = find( x==0 );
    indz = [];
    if any(diff(iz)==1)
      zer = x == 0;
      dz = diff([0 zer 0]);
      debz = find(dz == 1);
      finz = find(dz == -1)-1;
      indz = round((debz+finz)/2);
    else
      indz = iz;
    end
    indzer = sort([indzer indz]);
  end
end

d = diff(x);

n = length(d);
d1 = d(1:n-1);
d2 = d(2:n);
indmin = find(d1.*d2<0 & d1<0)+1;
indmax = find(d1.*d2<0 & d1>0)+1;


% when two or more successive points have the same value we consider only one extremum in the middle of the constant area
% (only works if the signal is uniformly sampled)

if any(d==0)

  imax = [];
  imin = [];

  bad = (d==0);
  dd = diff([0 bad 0]);
  debs = find(dd == 1);
  fins = find(dd == -1);
  if debs(1) == 1
    if length(debs) > 1
      debs = debs(2:end);
      fins = fins(2:end);
    else
      debs = [];
      fins = [];
    end
  end
  if length(debs) > 0
    if fins(end) == m
      if length(debs) > 1
        debs = debs(1:(end-1));
        fins = fins(1:(end-1));

      else
        debs = [];
        fins = [];
      end
    end
  end
  lc = length(debs);
  if lc > 0
    for k = 1:lc
      if d(debs(k)-1) > 0
        if d(fins(k)) < 0
          imax = [imax round((fins(k)+debs(k))/2)];
        end
      else
        if d(fins(k)) > 0
          imin = [imin round((fins(k)+debs(k))/2)];
        end
      end
    end
  end

  if length(imax) > 0
    indmax = sort([indmax imax]);
  end

  if length(imin) > 0
    indmin = sort([indmin imin]);
  end

end
end

%---------------------------------------------------------------------------------------------------

function ort = io(x,imf)
% ort = IO(x,imf) computes the index of orthogonality
%
% inputs : - x    : analyzed signal
%          - imf  : empirical mode decomposition

n = size(imf,1);

s = 0;

for i = 1:n
  for j =1:n
    if i~=j
      s = s + abs(sum(imf(i,:).*conj(imf(j,:)))/sum(x.^2));
    end
  end
end

ort = 0.5*s;
end
%---------------------------------------------------------------------------------------------------

function [x,t,sd,sd2,tol,MODE_COMPLEX,ndirs,display_sifting,sdt,sd2t,r,imf,k,nbit,NbIt,MAXITERATIONS,FIXE,FIXE_H,MAXMODES,INTERP,mask] = init(varargin)

x = varargin{1};
if nargin == 2
  if isstruct(varargin{2})
    inopts = varargin{2};
  else
    error('when using 2 arguments the first one is the analyzed signal X and the second one is a struct object describing the options')
  end
elseif nargin > 2
  try
    inopts = struct(varargin{2:end});
  catch
    error('bad argument syntax')
  end
end

% default for stopping
defstop = [0.05,0.5,0.05];

opt_fields = {'t','stop','display','maxiterations','fix','maxmodes','interp','fix_h','mask','ndirs','complex_version'};

defopts.stop = defstop;
defopts.display = 0;
defopts.t = 1:max(size(x));
defopts.maxiterations = 2000;
defopts.fix = 0;
defopts.maxmodes = 0;
defopts.interp = 'spline';
defopts.fix_h = 0;
defopts.mask = 0;
defopts.ndirs = 4;
defopts.complex_version = 2;

opts = defopts;


if(nargin==1)
  inopts = defopts;
elseif nargin == 0
  error('not enough arguments')
end


names = fieldnames(inopts);
for nom = names'
  if ~any(strcmpi(char(nom), opt_fields))
    error(['bad option field name: ',char(nom)])
  end
  if ~isempty(eval(['inopts.',char(nom)])) % empty values are discarded
    eval(['opts.',lower(char(nom)),' = inopts.',char(nom),';'])
  end
end

t = opts.t;
stop = opts.stop;
display_sifting = opts.display;
MAXITERATIONS = opts.maxiterations;
FIXE = opts.fix;
MAXMODES = opts.maxmodes;
INTERP = opts.interp;
FIXE_H = opts.fix_h;
mask = opts.mask;
ndirs = opts.ndirs;
complex_version = opts.complex_version;

if ~isvector(x)
  error('X must have only one row or one column')
end

if size(x,1) > 1
  x = x.';
end

if ~isvector(t)
  error('option field T must have only one row or one column')
end

if ~isreal(t)
  error('time instants T must be a real vector')
end

if size(t,1) > 1
  t = t';
end

if (length(t)~=length(x))
  error('X and option field T must have the same length')
end

if ~isvector(stop) || length(stop) > 3
  error('option field STOP must have only one row or one column of max three elements')
end

if ~all(isfinite(x))
  error('data elements must be finite')
end

if size(stop,1) > 1
  stop = stop';
end

L = length(stop);
if L < 3
  stop(3)=defstop(3);
end

if L < 2
  stop(2)=defstop(2);
end


if ~ischar(INTERP) || ~any(strcmpi(INTERP,{'linear','cubic','spline'}))
  error('INTERP field must be ''linear'', ''cubic'', ''pchip'' or ''spline''')
end

%special procedure when a masking signal is specified
if any(mask)
  if ~isvector(mask) || length(mask) ~= length(x)
    error('masking signal must have the same dimension as the analyzed signal X')
  end

  if size(mask,1) > 1
    mask = mask.';
  end
  opts.mask = 0;
  imf1 = emd(x+mask,opts);
  imf2 = emd(x-mask,opts);
  if size(imf1,1) ~= size(imf2,1)
    warning('emd:warning',['the two sets of IMFs have different sizes: ',int2str(size(imf1,1)),' and ',int2str(size(imf2,1)),' IMFs.'])
  end
  S1 = size(imf1,1);
  S2 = size(imf2,1);
  if S1 ~= S2
    if S1 < S2
      tmp = imf1;
      imf1 = imf2;
      imf2 = tmp;
    end
    imf2(max(S1,S2),1) = 0;
  end
  imf = (imf1+imf2)/2;

end


sd = stop(1);
sd2 = stop(2);
tol = stop(3);

lx = length(x);

sdt = sd*ones(1,lx);
sd2t = sd2*ones(1,lx);

if FIXE
  MAXITERATIONS = FIXE;
  if FIXE_H
    error('cannot use both ''FIX'' and ''FIX_H'' modes')
  end
end

MODE_COMPLEX = ~isreal(x)*complex_version;
if MODE_COMPLEX && complex_version ~= 1 && complex_version ~= 2
  error('COMPLEX_VERSION parameter must equal 1 or 2')
end


% number of extrema and zero-crossings in residual
ner = lx;
nzr = lx;

r = x;

if ~any(mask) % if a masking signal is specified "imf" already exists at this stage
  imf = [];
end
k = 1;

% iterations counter for extraction of 1 mode
nbit=0;

% total iterations counter
NbIt=0;
end
%---------------------------------------------------------------------------------------------------
```
