## Supplementary material for "Instantaneous phase of rhythmic behaviour under volitional control": Code and data files: ffgn.m

```
% Generates *exact* paths of Fractional Gaussian Noise by using
% circulant embedding (for 1/2<H<1) and Lowen's method (for 0<H<1/2).  
%
% Input: 
%   sigma <- standard deviation of the time series
%   H     <- Hurst exponent
%   n     <- number of independent paths
%   N     <- the length of the time series
%   force <- if *1*, then works with the smallest integer power of 2,
%            greater than or equal to N.
%
% Output:
%   f     <- a (nxN) matrix.  Each row contains an independently generated
%            path of FGN.
% 
% Usage:
%   f = ffgn(sigma,H,n,N,force);
%
% Written jointly by Yingchun Zhou (Jasmine), 
% and Stilian Stoev,, September, 2005.
%
% Example:
%
%  f = ffgn(1,0.8,1,2^16,0); plot(cumsum(f)); title('FBM, H=0.8');
function f = ffgn(sigma,H,n,N,force);
if force,
 N2 = 2^(fix(log2(N)));
 if N2 <N,
  N = 2*N2;
 else
 N  = N2;
 end;
end;
if (H>0.5)&(H<1),
%%
%% Use the "circulant ebedding" technique.  This method
%% works only in the case when 1/2 < H < 1.
%%
%%
 % First step: specify the covariance
 c = sigma^2/2*(([0:N]+1).^(2*H)-2*([0:N].^(2*H))+abs([0:N]-1).^(2*H));
 v = [c(1:N), c(N+1:-1:2)];
 % Second step: calculate Fourier transform of c
 g = real(fft(v));
 if min(g)<0,
  error(' Some of the g(k) are negative!');
 end;
 g = abs(g);
 % Third step: generate {z(1),...,z(2*N)}
 z(:,1)=sqrt(2)*randn(n,1);
 y(:,1)=z(:,1);
 z(:,N+1)=sqrt(2)*randn(n,1);
 y(:,N+1)=z(:,N+1);
 a=randn(n,N-1);
 b=randn(n,N-1);
 z1=a+b*i;
 z(:,2:N)=z1;
 y1=z1;
 y(:,2:N)=y1;
 y(:,N+2:2*N)=conj(y(:,N:-1:2));
 y = y.*(ones(n,1)*sqrt(g));
 % Fourth step: calculate the stationary process f
 f = real(fft(y')')/sqrt(4*N);
 f = f(:,1:N);
elseif H == 0.5,
  f = randn(n,N);
elseif (H<0.5)&(H>0),
%%
%% Use Lowen's method for this case.  Copied from the code "fftfgn.m"
%%
     G1=randn(n,N-1);  
     G2=randn(n,N-1);
     G = (G1+sqrt(-1)*G2)/sqrt(2);  
     GN = randn(n,1);
     G0 = zeros(n,1);
     H2=2*H;       
     R=(1-((1:N-1)/N).^H2);
     R=[1 R 0 R(N-1:-1:1)];
     S=ones(n,1)*(abs(fft(R,2*N)).^.5);
     X=[zeros(n,1) G, GN, conj(G(:,N-1:-1:1)) ].*S;  
     x=ifft(X',2*N)';
     y=sqrt(N)*real((x(:,1:N)-x(:,1)*ones(1,N)));
     f = sigma*N^H*[y(:,1), diff(y')'];
else
 error(' The value of the Hurst parameter H must be in (0,1).');
end;
```
