## Supplementary material for "Instantaneous phase of rhythmic behaviour under volitional control": Code and data files: hurst.m

```
function H = hurst(x)
% Vilen Abramov (2023). Hurst Exponent Estimation 
% (https://www.mathworks.com/matlabcentral/fileexchange/39069-hurst-exponent-estimation), 
% MATLAB Central File Exchange. Retrieved October 16, 2023. 

% Creates local matrix for calculations
X = tril(repmat(x, length(x), 1));
X(1,:) = [];
% Makes zeros equal NaN for calculations
X(X == 0) = NaN;
% Range and Std Dev
R = max(cumsum(X - mean(X,2,'omitnan'),2),[],2) - min(cumsum(X - mean(X,2,'omitnan'),2),[],2);
S = nanstd(X,0,2);
% Plot for visualization
% plot(log10(2:length(x)), log10((R./S)'), '-')
% Power law fitting
powerLaw = polyfit(log10(2:length(x)), log10((R./S)'), 1);
% Hurst exponent
H = powerLaw(1);
```
