## Supplementary material for "Instantaneous phase of rhythmic behaviour under volitional control": Code and data files: quadAngle.m

```
function quadAngle = quadAngle(nIMF,hilbQuad)
%
% INPUT:
%       nIMF     - matrix containing normalized IMFs to calculate analytical signal on
%       hilbQuad - toggle : 0 (default),1. If hilbQuad==1 then hilbert
%                  quadrature is used (mask = sign(hilbert(data)) otherwise direct  
%                  quadrature (mask= ((diff(data)>0) .* -2) + 1; mask(end+1) = mask(end);)
% OUTPUT:
%       quadAngle: array of angles obtained from data and quadrature
%                 signals
%

if nargin==1;
    hilbQuad=0;
end

[npt,ncol] = size(nIMF);

%flip data if needed 
flipped=0;
if (ncol > npt)
    flipped=1;
    nIMF=nIMF';
    [npt,ncol] = size(nIMF);
end

%initialize quadrature array
quadrature=zeros(size(nIMF,1),ncol);
%loop over columns
for i=1:ncol
        data = nIMF(:,i);
       %create mask
        if hilbQuad==0 %quadrature from signal direction of change
            mask = ((diff(data)>0) .* -2) + 1;
            mask(end+1) = mask(end);
        else %quadrature from sign of Hilbert transform
            P=hilbert(data);
            mask=sign(imag(P));
        end
        % quadrature value
        y = real(sqrt(1-data.^2));
    
        %flip data in 3rd & 4th quadrants
        q = y .* mask;
        
        %store column
        quadrature(:,i) = complex(data, q);
end
quadAngle=wrapTo2Pi(unwrap(angle(quadrature)));%compute angles

%flip data bask if needed 

if flipped==1
    quadAngle = quadAngle';
end
```
