## Supplementary material for "Instantaneous phase of rhythmic behaviour under volitional control": Code and data files: SG_calc.m

```
function coef = SG_calc(m,n,h,varargin)
%                            Savitzky-Golay Filter
%________________________________________________________________________________
%
%                            coef = SG_calc(m,n,h,M)
%
% The function calculates convolution coeffitients for the Savitzky-Golay filter.
% It approximates the given data by a polynomial and calculates a derivative by 
% analytical dereviation of the polynomial. 
%
% COMPULSORY INPUTS :
%
% m - number of smoothed points
% n - order of the fitting polynomial
% h - distance between the measured points
%
% OPTIONAL INPUTS :
%
% M - query point position
%
% OUTPUT:
%
% coef - (n+1 ; m ) matrix, each row contains m FIR filter coefficients for  
% j-th order differentiation, j = 0,1,2,...n. j = 0 represents a common 
% low-pass filter (zero-order derivative).
%
%
% EXAMPLE 1:
%
% - symetrical FIR filter (m can be odd or even)
% - the query point is in the middle of the interval
%
% m = 9;      % number of filtered points (FIR order)
% n = 2;      % approximation polynomial order
% h = 1e-2;   % sample step
% 
% coef = SG_calc(m,n,h);
% y   = filter(coef(1,:),1,y);    % low-pass filter
% dy  = filter(coef(2,:),1,y);    % 1st order differenciator
% ddy = filter(coef(3,:),1,y);    % 2nd order differenciator
%
% EXAMPLE 2:
% - asymetrical FIR filter (m can be odd or even)
% - the query point position is arbitryry, any nuber (even decimal) is allowed
% - Preferably, 1<=M<=m
%
%% m = 9;      % number of filtered points (FIR order)
% n = 2;      % approximation polynomial order
% h = 1e-2;   % sample step
% M = 7.5;    % asymetrical query point position
% 
% coef = SG_calc(m,n,h,M);
% y   = filter(coef(1,:),1,y);    % low-pass filter
% dy  = filter(coef(2,:),1,y);    % 1st order differenciator
% ddy = filter(coef(3,:),1,y);    % 2nd order differenciator
%
% REFERENCE
% M. Brablc, V. Sova, and R. Grepl, ?Adaptive feedforward controller for a 
% DC motor drive based on inverse dynamic model with recursive least squares 
% parameter estimation,? in Proceedings of the 2016 17th International Conference 
% on Mechatronics - Mechatronika, ME 2016, D. Maga and T. Brezina, Eds. Prague, 2017, pp. 146?150.
%
%________________________________________________________________________________
% See also SGOLAY
% Martin (2023). Savitzky-Golay Differentiation FIR filter (Generalized) (https://www.mathworks.com/matlabcentral/fileexchange/68342-savitzky-golay-differentiation-fir-filter-generalized), MATLAB Central File Exchange. Retrieved August 14, 2023. 
%% Check inputs
if nargin < 3
    error('Some input parameters are missing!')
elseif nargin == 3 % symmetrical filter (even/odd number of points)
    M = (m+1)/2;
elseif nargin == 4 % specified query point position
    M = varargin{1};
else
    error('Too many input parameters')
end
if (m ~= round(m)) || (m < 1)
    error('FIR filter order (m) has to be a positive integer!')
elseif (n ~= round(n)) || (n < 1)
    error('Approximation polynomial order (n) has to be a positive integer!')
elseif h <= 0
    error('Step size (h) has to be positive!')
end
if (M < 1) || (M > m) 
    warning('Be careful, the query point is outside the filter points (defined by m)!')
elseif (m == 1)
    warning('First order FIR filter is not a filter => no smoothing occures!')
elseif m < (n+1)
    warning('Incorect input parameters: m has to be >= n+1 !')
    error('Unable to exactly approximate the data, the polynomial has more parameters than the data points!')
elseif m == (n+1)
    warning('The approximation polynomial has the same number of parameters (n+1) as the data points (m), no smoothing!')
end
%% Savitzky-Golay FIR design
z = m-M:-1:1-M; 
J = ones(m,n+1);
for i = 1:m
    for j = 1:n
        J(i,j+1) = (z(i)*h)^j;
    end
end
scl = diag(factorial(0:n));     % scaling coefficients
coef = scl*((J'*J)\J');         % scaled LS estimate
```
