## Supplementary material for "Instantaneous phase of rhythmic behaviour under volitional control": Code and data files: downsample.m

```
% Author: Paul Kienzle <> (2007)
% This program is granted to the public domain.

% -*- texinfo -*-
% @deftypefn  {Function File} {@var{y} =} downsample (@var{x}, @var{n})
% @deftypefnx {Function File} {@var{y} =} downsample (@var{x}, @var{n}, @var{offset})
% Downsample the signal, selecting every @var{n}th element.  If @var{x}
% is a matrix, downsample every column.
%
% For most signals you will want to use @code{decimate} instead since
% it prefilters the high frequency components of the signal and
% avoids aliasing effects.
%
% If @var{offset} is defined, select every @var{n}th element starting at
% sample @var{offset}.
% @seealso{decimate, interp, resample, upfirdn, upsample}
% @end deftypefn

function y = downsample(x, n, phase)

if nargin<2 || nargin>3, print_usage; end

if nargin<3
  phase = 0;
end

if phase > n - 1
  warning('This is incompatible with Matlab (phase = 0:n-1). See octave-forge signal package release notes for details.')
end

if isvector(x)
  y = x(phase + 1:n:end);
else
  y = x(phase + 1:n:end,:);
end


%!assert(downsample([1,2,3,4,5],2),[1,3,5]);
%!assert(downsample([1;2;3;4;5],2),[1;3;5]);
%!assert(downsample([1,2;3,4;5,6;7,8;9,10],2),[1,2;5,6;9,10]);
%!assert(downsample([1,2,3,4,5],2,1),[2,4]);
%!assert(downsample([1,2;3,4;5,6;7,8;9,10],2,1),[3,4;7,8]);
```
