## Supplementary material for "Instantaneous phase of rhythmic behaviour under volitional control": Code and data files: findpeaks.m

```
function [Ypk,Xpk,Wpk,Ppk,wxPk] = findpeaks(Yin,varargin)
%FINDPEAKS Find local peaks in data
%   PKS = FINDPEAKS(Y) finds local peaks in the data vector Y. A local peak
%   is defined as a data sample which is either larger than the two
%   neighboring samples or is equal to Inf.
%
%   [PKS,LOCS]= FINDPEAKS(Y) also returns the indices LOCS at which the
%   peaks occur.
%
%   [PKS,LOCS] = FINDPEAKS(Y,X) specifies X as the location vector of data
%   vector Y. X must be a strictly increasing vector of the same length as
%   Y. LOCS returns the corresponding value of X for each peak detected.
%   If X is omitted, then X will correspond to the indices of Y.
%
%   [PKS,LOCS] = FINDPEAKS(Y,Fs) specifies the sample rate, Fs, as a
%   positive scalar, where the first sample instant of Y corresponds to a
%   time of zero.
%
%   [...] = FINDPEAKS(...,'MinPeakHeight',MPH) finds only those peaks that
%   are greater than the minimum peak height, MPH. MPH is a real valued
%   scalar. The default value of MPH is -Inf.
%
%   [...] = FINDPEAKS(...,'MinPeakProminence',MPP) finds peaks guaranteed
%   to have a vertical drop of more than MPP from the peak on both sides
%   without encountering either the end of the signal or a larger
%   intervening peak. The default value of MPP is zero.
%
%   [...] = FINDPEAKS(...,'Threshold',TH) finds peaks that are at least
%   greater than both adjacent samples by the threshold, TH. TH is real
%   valued scalar greater than or equal to zero. The default value of TH is
%   zero.
%
%   FINDPEAKS(...,'WidthReference',WR) estimates the width of the peak as
%   the distance between the points where the signal intercepts a
%   horizontal reference line. The points are found by linear
%   interpolation. The height of the line is selected using the criterion
%   specified in WR:
% 
%    'halfprom' - the reference line is positioned beneath the peak at a
%       vertical distance equal to half the peak prominence.
% 
%    'halfheight' - the reference line is positioned at one-half the peak 
%       height. The line is truncated if any of its intercept points lie
%       beyond the borders of the peaks selected by the 'MinPeakHeight',
%       'MinPeakProminence' and 'Threshold' parameters. The border between
%       peaks is defined by the horizontal position of the lowest valley
%       between them. Peaks with heights less than zero are discarded.
% 
%    The default value of WR is 'halfprom'.
%
%   [...] = FINDPEAKS(...,'MinPeakWidth',MINW) finds peaks whose width is
%   at least MINW. The default value of MINW is zero.
%
%   [...] = FINDPEAKS(...,'MaxPeakWidth',MAXW) finds peaks whose width is
%   at most MAXW. The default value of MAXW is Inf.
%
%   [...] = FINDPEAKS(...,'MinPeakDistance',MPD) finds peaks separated by
%   more than the minimum peak distance, MPD. This parameter may be
%   specified to ignore smaller peaks that may occur in close proximity to
%   a large local peak. For example, if a large local peak occurs at LOC,
%   then all smaller peaks in the range [N-MPD, N+MPD] are ignored. If not
%   specified, MPD is assigned a value of zero.
%
%   [...] = FINDPEAKS(...,'SortStr',DIR) specifies the direction of sorting
%   of peaks. DIR can take values of 'ascend', 'descend' or 'none'. If not
%   specified, DIR takes the value of 'none' and the peaks are returned in
%   the order of their occurrence.
%
%   [...] = FINDPEAKS(...,'NPeaks',NP) specifies the maximum number of peaks
%   to be found. NP is an integer greater than zero. If not specified, all
%   peaks are returned. Use this parameter in conjunction with setting the
%   sort direction to 'descend' to return the NP largest peaks. (see
%   'SortStr')
%
%   [PKS,LOCS,W] = FINDPEAKS(...) returns the width, W, of each peak by
%   linear interpolation of the left- and right- intercept points to the
%   reference defined by 'WidthReference'.
%
%   [PKS,LOCS,W,P] = FINDPEAKS(...) returns the prominence, P, of each
%   peak.
%
%   FINDPEAKS(...) without output arguments plots the signal and the peak
%   values it finds
%
%   FINDPEAKS(...,'Annotate',PLOTSTYLE) will annotate a plot of the
%   signal with PLOTSTYLE. If PLOTSTYLE is 'peaks' the peaks will be
%   plotted. If PLOTSTYLE is 'extents' the signal, peak values, widths,
%   prominences of each peak will be annotated. 'Annotate' will be ignored
%   if called with output arguments. The default value of PLOTSTYLE is
%   'peaks'.
%
%   % Example 1:
%   %   Plot the Zurich numbers of sunspot activity from years 1700-1987
%   %   and identify all local maxima at least six years apart
%   load sunspot.dat
%   findpeaks(sunspot(:,2),sunspot(:,1),'MinPeakDistance',6)
%   xlabel('Year');
%   ylabel('Zurich number');
%
%   % Example 2: 
%   %   Plot peak values of an audio signal that drop at least 1V on either
%   %   side without encountering values larger than the peak.
%   load mtlb
%   findpeaks(mtlb,Fs,'MinPeakProminence',1)
%
%   % Example 3:
%   %   Plot all peaks of a chirp signal whose widths are between .5 and 1 
%   %   milliseconds.
%   Fs = 44.1e3; N = 1000;
%   x = sin(2*pi*(1:N)/N + (10*(1:N)/N).^2);
%   findpeaks(x,Fs,'MinPeakWidth',.5e-3,'MaxPeakWidth',1e-3, ...
%             'Annotate','extents')

%   Copyright 2007-2014 The MathWorks, Inc.

%#ok<*EMCLS>
%#ok<*EMCA>
%#codegen

cond = nargin >= 1;
if ~cond
    coder.internal.assert(cond,'MATLAB:narginchk:notEnoughInputs');
end

cond = nargin <= 22;
if ~cond
    coder.internal.assert(cond,'MATLAB:narginchk:tooManyInputs');
end

% extract the parameters from the input argument list
[y,yIsRow,x,xIsRow,minH,minP,minW,maxW,minD,minT,maxN,sortDir,annotate,refW] ...
  = parse_inputs(Yin,varargin{:});

% find indices of all finite and infinite peaks and the inflection points
[iFinite,iInfite,iInflect] = getAllPeaks(y);

% keep only the indices of finite peaks that meet the required 
% minimum height and threshold
iPk = removePeaksBelowMinPeakHeight(y,iFinite,minH,refW);
iPk = removePeaksBelowThreshold(y,iPk,minT);

% indicate if we need to compute the extent of a peak
needWidth = minW>0 || maxW<inf || minP>0 || nargout>2 || strcmp(annotate,'extents');

if needWidth
  % obtain the indices of each peak (iPk), the prominence base (bPk), and
  % the x- and y- coordinates of the peak base (bxPk, byPk) and the width
  % (wxPk)
  [iPk,bPk,bxPk,byPk,wxPk] = findExtents(y,x,iPk,iFinite,iInfite,iInflect,minP,minW,maxW,refW);
else
  % combine finite and infinite peaks into one list
  [iPk,bPk,bxPk,byPk,wxPk] = combinePeaks(iPk,iInfite);
end

% find the indices of the largest peaks within the specified distance
idx = findPeaksSeparatedByMoreThanMinPeakDistance(y,x,iPk,minD);

% re-order and bound the number of peaks based upon the index vector
idx = orderPeaks(y,iPk,idx,sortDir);
idx = keepAtMostNpPeaks(idx,maxN);

% use the index vector to fecth the correct peaks.
iPk = iPk(idx);
if needWidth
  [bPk, bxPk, byPk, wxPk] = fetchPeakExtents(idx,bPk,bxPk,byPk,wxPk);
end

if nargout > 0
  % assign output variables
  if needWidth
    [Ypk,Xpk,Wpk,Ppk] = assignFullOutputs(y,x,iPk,wxPk,bPk,yIsRow,xIsRow);
  else
    [Ypk,Xpk] = assignOutputs(y,x,iPk,yIsRow,xIsRow);
  end    
else
  % no output arguments specified. plot and optionally annotate
  hAxes = plotSignalWithPeaks(x,y,iPk);
  if strcmp(annotate,'extents')
    plotExtents(hAxes,x,y,iPk,bPk,bxPk,byPk,wxPk,refW);
  end
  
  scalePlot(hAxes);
end

%--------------------------------------------------------------------------
function [y,yIsRow,x,xIsRow,Ph,Pp,Wmin,Wmax,Pd,Th,NpOut,Str,Ann,Ref] = parse_inputs(Yin,varargin)

% Validate input signal
validateattributes(Yin,{'numeric'},{'nonempty','real','vector'},...
    'findpeaks','Y');
yIsRow = isrow(Yin);
y = Yin(:);

% copy over orientation of y to x.
xIsRow = yIsRow;

% indicate if the user specified an Fs or X
hasX = ~isempty(varargin) && isnumeric(varargin{1});

if hasX
  startArg = 2;
  if isscalar(varargin{1})
    % Fs
    Fs = varargin{1};
    validateattributes(Fs,{'double'},{'real','finite','positive'},'findpeaks','Fs');
    x = (0:numel(y)-1).'/Fs;
  else
    % X
    Xin = varargin{1};
    validateattributes(Xin,{'double'},{'real','finite','vector','increasing'},'findpeaks','X');
    if numel(Xin) ~= numel(Yin)
      if coder.target('MATLAB')
        throwAsCaller(MException(message('signal:findpeaks:mismatchYX')));
      else
        coder.internal.errorIf(true,'signal:findpeaks:mismatchYX');
      end
    end
    xIsRow = isrow(Xin);
    x = Xin(:);
  end
else
  startArg = 1;
  % unspecified, use index vector
  x = (1:numel(y)).';
end

if coder.target('MATLAB')
    try %#ok<EMTC>
        % Check the input data type. Single precision is not supported.
        chkinputdatatype(y);
        chkinputdatatype(x);
    catch ME
        throwAsCaller(ME);
    end
else
    chkinputdatatype(y);
    chkinputdatatype(x);
end

M = numel(y);
cond = (M < 3);
if cond
    coder.internal.errorIf(cond,'signal:findpeaks:emptyDataSet');
end

%#function dspopts.findpeaks
defaultMinPeakHeight = -inf;
defaultMinPeakProminence = 0;
defaultMinPeakWidth = 0;
defaultMaxPeakWidth = Inf;
defaultMinPeakDistance = 0;
defaultThreshold = 0;
defaultNPeaks = [];
defaultSortStr = 'none';
defaultAnnotate = 'peaks';
defaultWidthReference = 'halfprom';

if coder.target('MATLAB')
    p = inputParser;
    addParameter(p,'MinPeakHeight',defaultMinPeakHeight);
    addParameter(p,'MinPeakProminence',defaultMinPeakProminence);
    addParameter(p,'MinPeakWidth',defaultMinPeakWidth);
    addParameter(p,'MaxPeakWidth',defaultMaxPeakWidth);
    addParameter(p,'MinPeakDistance',defaultMinPeakDistance);
    addParameter(p,'Threshold',defaultThreshold);
    addParameter(p,'NPeaks',defaultNPeaks);
    addParameter(p,'SortStr',defaultSortStr);
    addParameter(p,'Annotate',defaultAnnotate);
    addParameter(p,'WidthReference',defaultWidthReference);
    parse(p,varargin{startArg:end});
    Ph = p.Results.MinPeakHeight;
    Pp = p.Results.MinPeakProminence;
    Wmin = p.Results.MinPeakWidth;
    Wmax = p.Results.MaxPeakWidth;
    Pd = p.Results.MinPeakDistance;
    Th = p.Results.Threshold;
    Np = p.Results.NPeaks;
    Str = p.Results.SortStr;
    Ann = p.Results.Annotate;
    Ref = p.Results.WidthReference;
else
    parms = struct('MinPeakHeight',uint32(0), ...
                'MinPeakProminence',uint32(0), ...
                'MinPeakWidth',uint32(0), ...
                'MaxPeakWidth',uint32(0), ...
                'MinPeakDistance',uint32(0), ...
                'Threshold',uint32(0), ...
                'NPeaks',uint32(0), ...
                'SortStr',uint32(0), ...
                'Annotate',uint32(0), ...
                'WidthReference',uint32(0));
    pstruct = eml_parse_parameter_inputs(parms,[],varargin{startArg:end});
    Ph = eml_get_parameter_value(pstruct.MinPeakHeight,defaultMinPeakHeight,varargin{startArg:end});
    Pp = eml_get_parameter_value(pstruct.MinPeakProminence,defaultMinPeakProminence,varargin{startArg:end});
    Wmin = eml_get_parameter_value(pstruct.MinPeakWidth,defaultMinPeakWidth,varargin{startArg:end});
    Wmax = eml_get_parameter_value(pstruct.MaxPeakWidth,defaultMaxPeakWidth,varargin{startArg:end});
    Pd = eml_get_parameter_value(pstruct.MinPeakDistance,defaultMinPeakDistance,varargin{startArg:end});
    Th = eml_get_parameter_value(pstruct.Threshold,defaultThreshold,varargin{startArg:end});
    Np = eml_get_parameter_value(pstruct.NPeaks,defaultNPeaks,varargin{startArg:end});
    Str = eml_get_parameter_value(pstruct.SortStr,defaultSortStr,varargin{startArg:end});
    Ann = eml_get_parameter_value(pstruct.Annotate,defaultAnnotate,varargin{startArg:end});
    Ref = eml_get_parameter_value(pstruct.WidthReference,defaultWidthReference,varargin{startArg:end});
end

% limit the number of peaks to the number of input samples
if isempty(Np)
    NpOut = M;
else
    NpOut = Np;
end

% ignore peaks below zero when using halfheight width reference
if strcmp(Ref,'halfheight')
  Ph = max(Ph,0);
end

validateattributes(Ph,{'numeric'},{'real','scalar','nonempty'},'findpeaks','MinPeakHeight');
validateattributes(Pd,{'numeric'},{'real','scalar','nonempty','nonnegative','<',x(M)-x(1)},'findpeaks','MinPeakDistance');
validateattributes(Pp,{'numeric'},{'real','scalar','nonempty','nonnegative'},'findpeaks','MinPeakProminence');
validateattributes(Wmin,{'numeric'},{'real','scalar','finite','nonempty','nonnegative'},'findpeaks','MinPeakWidth');
validateattributes(Wmax,{'numeric'},{'real','scalar','nonnan','nonempty','nonnegative'},'findpeaks','MaxPeakWidth');
validateattributes(Pd,{'numeric'},{'real','scalar','nonempty','nonnegative'},'findpeaks','MinPeakDistance');
validateattributes(Th,{'numeric'},{'real','scalar','nonempty','nonnegative'},'findpeaks','Threshold');
validateattributes(NpOut,{'numeric'},{'real','scalar','nonempty','integer','positive'},'findpeaks','NPeaks');
Str = validatestring(Str,{'ascend','none','descend'},'findpeaks','SortStr');
Ann = validatestring(Ann,{'peaks','extents'},'findpeaks','SortStr');
Ref = validatestring(Ref,{'halfprom','halfheight'},'findpeaks','WidthReference');

%--------------------------------------------------------------------------
function [iPk,iInf,iInflect] = getAllPeaks(y)
% fetch indices all infinite peaks
iInf = find(isinf(y) & y>0);

% temporarily remove all +Inf values
yTemp = y;
yTemp(iInf) = NaN;

% determine the peaks and inflection points of the signal
[iPk,iInflect] = findLocalMaxima(yTemp);


%--------------------------------------------------------------------------
function [iPk, iInflect] = findLocalMaxima(yTemp)
% bookend Y by NaN and make index vector
yTemp = [NaN; yTemp; NaN];
iTemp = (1:numel(yTemp)).';

% keep only the first of any adjacent pairs of equal values (including NaN).
yFinite = ~isnan(yTemp);
iNeq = [1; 1 + find((yTemp(1:end-1) ~= yTemp(2:end)) & ...
                    (yFinite(1:end-1) | yFinite(2:end)))];
iTemp = iTemp(iNeq);

% take the sign of the first sample derivative
s = sign(diff(yTemp(iTemp)));

% find local maxima
iMax = 1 + find(diff(s)<0);

% find all transitions from rising to falling or to NaN
iAny = 1 + find(s(1:end-1)~=s(2:end));

% index into the original index vector without the NaN bookend.
iInflect = iTemp(iAny)-1;
iPk = iTemp(iMax)-1;

%--------------------------------------------------------------------------
function iPk = removePeaksBelowMinPeakHeight(Y,iPk,Ph,widthRef)
if ~isempty(iPk) 
  iPk = iPk(Y(iPk) > Ph);
  if isempty(iPk) && ~strcmp(widthRef,'halfheight')
    if coder.target('MATLAB')
        warning(message('signal:findpeaks:largeMinPeakHeight', 'MinPeakHeight', 'MinPeakHeight'));
    end
  end
end
    
%--------------------------------------------------------------------------
function iPk = removePeaksBelowThreshold(Y,iPk,Th)

base = max(Y(iPk-1),Y(iPk+1));
iPk = iPk(Y(iPk)-base >= Th);

%--------------------------------------------------------------------------
function [iPk,bPk,bxPk,byPk,wxPk] = findExtents(y,x,iPk,iFin,iInf,iInflect,minP,minW,maxW,refW)
% temporarily filter out +Inf from the input
yFinite = y;
yFinite(iInf) = NaN;

% get the base and left and right indices of each prominence base
[bPk,iLB,iRB] = getPeakBase(yFinite,iPk,iFin,iInflect);

% keep only those indices with at least the specified prominence
[iPk,bPk,iLB,iRB] = removePeaksBelowMinPeakProminence(yFinite,iPk,bPk,iLB,iRB,minP);

% get the x-coordinates of the half-height width borders of each peak
[wxPk,iLBh,iRBh] = getPeakWidth(yFinite,x,iPk,bPk,iLB,iRB,refW);

% merge finite and infinte peaks together into one list
[iPk,bPk,bxPk,byPk,wxPk] = combineFullPeaks(y,x,iPk,bPk,iLBh,iRBh,wxPk,iInf);

% keep only those in the range minW < w < maxW
[iPk,bPk,bxPk,byPk,wxPk] = removePeaksOutsideWidth(iPk,bPk,bxPk,byPk,wxPk,minW,maxW);


%--------------------------------------------------------------------------
function [peakBase,iLeftSaddle,iRightSaddle] = getPeakBase(yTemp,iPk,iFin,iInflect)
% determine the indices that border each finite peak
[iLeftBase, iLeftSaddle] = getLeftBase(yTemp,iPk,iFin,iInflect);
[iRightBase, iRightSaddle] = getLeftBase(yTemp,flipud(iPk),flipud(iFin),flipud(iInflect));
iRightBase = flipud(iRightBase);
iRightSaddle = flipud(iRightSaddle);
peakBase = max(yTemp(iLeftBase),yTemp(iRightBase));

%--------------------------------------------------------------------------
function [iBase, iSaddle] = getLeftBase(yTemp,iPeak,iFinite,iInflect)
% pre-initialize output base indices
iBase = zeros(size(iPeak));

% table stores the most recently encoutered peaks in order of height
peak = zeros(size(iPeak));
valley = zeros(size(iPeak));
iValley = zeros(size(iPeak));
iSaddle = zeros(size(iPeak));
n = 0;
i = 1;
j = 1;
k = 1;

% pre-initialize v for code generation
v = NaN; 
iv = 1;

while k<=numel(iPeak)
  % walk through the inflections until you reach a peak
  while iInflect(i) ~= iFinite(j) 
    v = yTemp(iInflect(i));
    iv = iInflect(i);
    if isnan(v)
      % border seen, start over.
      n = 0;
    else
      % ignore previously stored peaks with a valley larger than this one
      while n>0 && valley(n)>v;
        n = n - 1;
      end
    end
    i = i + 1;
  end
  % get the peak
  p = yTemp(iInflect(i));
  
  % keep the smallest valley of all smaller peaks
  while n>0 && peak(n) < p
    if valley(n) < v
      v = valley(n);
      iv = iValley(n);
    end
    n = n - 1;
  end

  % record "saddle" valleys in between equal-height peaks
  isv = iv;
  
  % keep seeking smaller valleys until you reach a larger peak
  while n>0 && peak(n) <= p
    if valley(n) < v
      v = valley(n);
      iv = iValley(n);
    end
    n = n - 1;      
  end
  
  % record the new peak and save the index of the valley into the base
  % and saddle
  n = n + 1;
  peak(n) = p;
  valley(n) = v;
  iValley(n) = iv;
  
  if iInflect(i) == iPeak(k)
    iBase(k) = iv;
    iSaddle(k) = isv;
    k = k + 1;
  end
  
  i = i + 1;
  j = j + 1;
end

%--------------------------------------------------------------------------
function [iPk,pbPk,iLB,iRB] = removePeaksBelowMinPeakProminence(y,iPk,pbPk,iLB,iRB,minP)
% compute the prominence of each peak
Ppk = y(iPk)-pbPk;

% keep those that are above the specified prominence
idx = find(Ppk >= minP);
iPk = iPk(idx);
pbPk = pbPk(idx);
iLB = iLB(idx);
iRB = iRB(idx);

%--------------------------------------------------------------------------
function [wxPk,iLBh,iRBh] = getPeakWidth(y,x,iPk,pbPk,iLB,iRB,wRef)
if isempty(iPk)
  % no peaks.  define empty containers
  base = zeros(size(iPk));
  iLBh = zeros(size(iPk));
  iRBh = zeros(size(iPk));
elseif strcmp(wRef,'halfheight')
  % set the baseline to zero
  base = zeros(size(iPk));

  % border the width by no more than the lowest valley between this peak
  % and the next peak
  iLBh = [iLB(1); max(iLB(2:end),iRB(1:end-1))];
  iRBh = [min(iRB(1:end-1),iLB(2:end)); iRB(end)];
  iGuard = iLBh > iPk;
  iLBh(iGuard) = iLB(iGuard);
  iGuard = iRBh < iPk;
  iRBh(iGuard) = iRB(iGuard);
else
  % use the prominence base
  base = pbPk;
  
  % border the width by the saddle of the peak
  iLBh = iLB;
  iRBh = iRB;
end

% get the width boundaries of each peak
wxPk = getHalfMaxBounds(y, x, iPk, base, iLBh, iRBh);

%--------------------------------------------------------------------------
function bounds = getHalfMaxBounds(y, x, iPk, base, iLB, iRB)
bounds = zeros(numel(iPk),2);

% interpolate both the left and right bounds clamping at borders
for i=1:numel(iPk)
  
  % compute the desired reference level at half-height or half-prominence
  refHeight = (y(iPk(i))+base(i))/2;
  
  % compute the index of the left-intercept at half max
  iLeft = findLeftIntercept(y, iPk(i), iLB(i), refHeight);
  if iLeft < iLB(i)
    xLeft = x(iLB(i));
  else
    xLeft = linterp(x(iLeft),x(iLeft+1),y(iLeft),y(iLeft+1),y(iPk(i)),base(i));
  end
  
  % compute the index of the right-intercept
  iRight = findRightIntercept(y, iPk(i), iRB(i), refHeight);
  if iRight > iRB(i)
    xRight = x(iRB(i));
  else
    xRight = linterp(x(iRight), x(iRight-1), y(iRight), y(iRight-1), y(iPk(i)),base(i));
  end

  % store result
  bounds(i,:) = [xLeft xRight];
end

%--------------------------------------------------------------------------
function idx = findLeftIntercept(y, idx, borderIdx, refHeight)
% decrement index until you pass under the reference height or pass the
% index of the left border, whichever comes first
while idx>=borderIdx && y(idx) > refHeight
  idx = idx - 1;
end

%--------------------------------------------------------------------------
function idx = findRightIntercept(y, idx, borderIdx, refHeight)
% increment index until you pass under the reference height or pass the
% index of the right border, whichever comes first
while idx<=borderIdx && y(idx) > refHeight
  idx = idx + 1;
end

%--------------------------------------------------------------------------
function xc = linterp(xa,xb,ya,yb,yc,bc)
% interpolate between points (xa,ya) and (xb,yb) to find (xc, 0.5*(yc-yc)).
xc = xa + (xb-xa) .* (0.5*(yc+bc)-ya) ./ (yb-ya);

% invoke L'Hospital's rule when -Inf is encountered. 
if isnan(xc)
  % yc and yb are guaranteed to be finite. 
  if isinf(bc)
    % both ya and bc are -Inf.
    xc = 0.5*(xa+xb);
  else
    % only ya is -Inf.
    xc = xb;
  end
end

%--------------------------------------------------------------------------
function [iPk,bPk,bxPk,byPk,wxPk] = removePeaksOutsideWidth(iPk,bPk,bxPk,byPk,wxPk,minW,maxW)

if isempty(iPk) || minW==0 && maxW == inf;
  return
end

% compute the width of each peak and extract the matching indices
w = diff(wxPk,1,2);
idx = find(minW <= w & w <= maxW);

% fetch the surviving peaks
iPk = iPk(idx);
bPk = bPk(idx);
bxPk = bxPk(idx,:);
byPk = byPk(idx,:);
wxPk = wxPk(idx,:);

%--------------------------------------------------------------------------
function [iPkOut,bPk,bxPk,byPk,wxPk] = combinePeaks(iPk,iInf)
iPkOut = union(iPk,iInf);
bPk = zeros(0,1);
bxPk = zeros(0,2);
byPk = zeros(0,2);
wxPk = zeros(0,2);

%--------------------------------------------------------------------------
function [iPkOut,bPkOut,bxPkOut,byPkOut,wxPkOut] = combineFullPeaks(y,x,iPk,bPk,iLBw,iRBw,wPk,iInf)
iPkOut = union(iPk, iInf);

% create map of new indices to old indices
[~, iFinite] = intersect(iPkOut,iPk);
[~, iInfinite] = intersect(iPkOut,iInf);

% prevent row concatenation when iPk and iInf both have less than one
% element
iPkOut = iPkOut(:);

% compute prominence base
bPkOut = zeros(size(iPkOut));
bPkOut(iFinite) = bPk;
bPkOut(iInfinite) = 0;

% compute indices of left and right infinite borders
iInfL = max(1,iInf-1);
iInfR = min(iInf+1,numel(x));

% copy out x- values of the left and right prominence base
% set each base border of an infinite peaks halfway between itself and
% the next adjacent sample
bxPkOut = zeros(size(iPkOut,1),2);
bxPkOut(iFinite,1) = x(iLBw);
bxPkOut(iFinite,2) = x(iRBw);
bxPkOut(iInfinite,1) = 0.5*(x(iInf)+x(iInfL));
bxPkOut(iInfinite,2) = 0.5*(x(iInf)+x(iInfR));

% copy out y- values of the left and right prominence base
byPkOut = zeros(size(iPkOut,1),2);
byPkOut(iFinite,1) = y(iLBw);
byPkOut(iFinite,2) = y(iRBw);
byPkOut(iInfinite,1) = y(iInfL);
byPkOut(iInfinite,2) = y(iInfR);

% copy out x- values of the width borders
% set each width borders of an infinite peaks halfway between itself and
% the next adjacent sample
wxPkOut = zeros(size(iPkOut,1),2);
wxPkOut(iFinite,:) = wPk;
wxPkOut(iInfinite,1) = 0.5*(x(iInf)+x(iInfL));
wxPkOut(iInfinite,2) = 0.5*(x(iInf)+x(iInfR));

%--------------------------------------------------------------------------
function idx = findPeaksSeparatedByMoreThanMinPeakDistance(y,x,iPk,Pd)
% Start with the larger peaks to make sure we don't accidentally keep a
% small peak and remove a large peak in its neighborhood. 

if isempty(iPk) || Pd==0
  idx = (1:numel(iPk)).';
  return
end

% copy peak values and locations to a temporary place
pks = y(iPk);
locs = x(iPk);

% Order peaks from large to small
[~, sortIdx] = sort(pks,'descend');
locs_temp = locs(sortIdx);

idelete = ones(size(locs_temp))<0;
for i = 1:length(locs_temp)
  if ~idelete(i)
    % If the peak is not in the neighborhood of a larger peak, find
    % secondary peaks to eliminate.
    idelete = idelete | (locs_temp>=locs_temp(i)-Pd)&(locs_temp<=locs_temp(i)+Pd); 
    idelete(i) = 0; % Keep current peak
  end
end

% report back indices in consecutive order
idx = sort(sortIdx(~idelete));


%--------------------------------------------------------------------------
function idx = orderPeaks(Y,iPk,idx,Str)

if isempty(idx) || strcmp(Str,'none')
  return
end

if strcmp(Str,'ascend')
  [~,s]  = sort(Y(iPk(idx)),'ascend');
else
  [~,s]  = sort(Y(iPk(idx)),'descend');
end

idx = idx(s);


%--------------------------------------------------------------------------
function idx = keepAtMostNpPeaks(idx,Np)

if length(idx)>Np
  idx = idx(1:Np);
end

%--------------------------------------------------------------------------
function [bPk,bxPk,byPk,wxPk] = fetchPeakExtents(idx,bPk,bxPk,byPk,wxPk)
bPk = bPk(idx);
bxPk = bxPk(idx,:);
byPk = byPk(idx,:);
wxPk = wxPk(idx,:);

%--------------------------------------------------------------------------
function [YpkOut,XpkOut] = assignOutputs(y,x,iPk,yIsRow,xIsRow)

% fetch the coordinates of the peak
Ypk = y(iPk);
Xpk = x(iPk);

% preserve orientation of Y
if yIsRow
  YpkOut = Ypk.';
else
  YpkOut = Ypk;
end

% preserve orientation of X
if xIsRow
  XpkOut = Xpk.';
else
  XpkOut = Xpk;
end

%--------------------------------------------------------------------------
function [YpkOut,XpkOut,WpkOut,PpkOut] = assignFullOutputs(y,x,iPk,wxPk,bPk,yIsRow,xIsRow)

% fetch the coordinates of the peak
Ypk = y(iPk);
Xpk = x(iPk);

% compute the width and prominence
Wpk = diff(wxPk,1,2);
Ppk = Ypk-bPk;

% preserve orientation of Y (and P)
if yIsRow
  YpkOut = Ypk.';
  PpkOut = Ppk.';
else
  YpkOut = Ypk;
  PpkOut = Ppk;  
end

% preserve orientation of X (and W)
if xIsRow
  XpkOut = Xpk.';
  WpkOut = Wpk.';
else
  XpkOut = Xpk;
  WpkOut = Wpk;  
end

%--------------------------------------------------------------------------
function hAxes = plotSignalWithPeaks(x,y,iPk)

% plot signal
hLine = plot(x,y,'Tag','Signal');
hAxes = ancestor(hLine,'Axes');
% turn on grid
grid on;

% use the color of the line
color = get(hLine,'Color');
hLine = line(x(iPk),y(iPk),'Parent',hAxes, ...
     'Marker','o','LineStyle','none','Color',color,'tag','Peak');

% if using MATLAB use offset inverted triangular marker
if coder.target('MATLAB')
  plotpkmarkers(hLine,y(iPk));
end
   
%--------------------------------------------------------------------------
function plotExtents(hAxes,x,y,iPk,bPk,bxPk,byPk,wxPk,refW)

% compute level of half-maximum (height or prominence)
if strcmp(refW,'halfheight')
  hm = 0.5*y(iPk);
else
  hm = 0.5*(y(iPk)+bPk);
end

% get the default color order
colors = get(0,'DefaultAxesColorOrder');

% plot boundaries between adjacent peaks when using half-height
if strcmp(refW,'halfheight')
  % plot height
  plotLines(hAxes,'Height',x(iPk),y(iPk),x(iPk),zeros(numel(iPk),1),colors(2,:));  

  % plot width
  plotLines(hAxes,'HalfHeightWidth',wxPk(:,1),hm,wxPk(:,2),hm,colors(3,:));
      
  % plot peak borders
  idx = find(byPk(:,1)>0);
  plotLines(hAxes,'Border',bxPk(idx,1),zeros(numel(idx),1),bxPk(idx,1),byPk(idx,1),colors(4,:));
  idx = find(byPk(:,2)>0);
  plotLines(hAxes,'Border',bxPk(idx,2),zeros(numel(idx),1),bxPk(idx,2),byPk(idx,2),colors(4,:));
  
else
  % plot prominence
  plotLines(hAxes,'Prominence',x(iPk), y(iPk), x(iPk), bPk, colors(2,:));  
  
  % plot width
  plotLines(hAxes,'HalfProminenceWidth',wxPk(:,1), hm, wxPk(:,2), hm, colors(3,:));
  
  % plot peak borders
  idx = find(bPk(:)<byPk(:,1));
  plotLines(hAxes,'Border',bxPk(idx,1),bPk(idx),bxPk(idx,1),byPk(idx,1),colors(4,:));
  idx = find(bPk(:)<byPk(:,2));
  plotLines(hAxes,'Border',bxPk(idx,2),bPk(idx),bxPk(idx,2),byPk(idx,2),colors(4,:));
end

if coder.target('MATLAB')
  hLine = get(hAxes,'Children');
  tags = get(hLine,'tag');
  
  legendStrs = {};
  searchTags = {'Signal','Peak','Prominence','Height','HalfProminenceWidth','HalfHeightWidth','Border'};
  for i=1:numel(searchTags)
    if any(strcmp(searchTags{i},tags))
      legendStrs = [legendStrs, ...
        {getString(message(['signal:findpeaks:Legend' searchTags{i}]))}]; %#ok<AGROW>
    end
  end
  
  if numel(hLine)==1
    legend(getString(message('signal:findpeaks:LegendSignalNoPeaks')), ...
      'Location','best');
  else
    legend(legendStrs,'Location','best');
  end  
end

%--------------------------------------------------------------------------
function plotLines(hAxes,tag,x1,y1,x2,y2,c)
% concatenate multiple lines into a single line and fencepost with NaN
n = numel(x1);
line(reshape([x1(:).'; x2(:).'; NaN(1,n)], 3*n, 1), ...
     reshape([y1(:).'; y2(:).'; NaN(1,n)], 3*n, 1), ...
     'Color',c,'Parent',hAxes,'tag',tag);

%--------------------------------------------------------------------------
function scalePlot(hAxes)

% In the event that the plot has integer valued y limits, 'axis auto' may
% clip the YLimits directly to the data with no margin.  We search every
% line for its max and minimum value and create a temporary annotation that
% is 10% larger than the min and max values.  We then feed this to "axis
% auto", save the y limits, set axis to "tight" then restory the y limits.
% This obviates the need to check each line for its max and minimum x
% values as well.

minVal = Inf;
maxVal = -Inf;

if coder.target('MATLAB')
  hLines = findall(hAxes,'Type','line');
  for i=1:numel(hLines)
    data = get(hLines(i),'YData');
    data = data(isfinite(data));
    if ~isempty(data)
      minVal = min(minVal, min(data(:)));
      maxVal = max(maxVal, max(data(:)));
    end
  end
  
  axis auto
  xlimits = xlim;
  
  % grow upper and lower y extent by 5% (a total of 10%)
  p = .05;    
  y1 = (1+p)*maxVal - p*minVal;
  y2 = (1+p)*minVal - p*maxVal;
  
  % artificially expand the data range by the specified amount
  hTempLine = line(xlimits([1 1]),[y1 y2],'Parent',hAxes);  
  
  % save the limits
  ylimits = ylim;
  delete(hTempLine);  
else
  axis auto
  ylimits = ylim;
end

% preserve expanded y limits but tighten x axis.
axis tight
ylim(ylimits);  

% [EOF]

function chkinputdatatype(varargin)
%CHKINPUTDATATYPE Check that all inputs are double

%   Copyright 2009-2013 The MathWorks, Inc.


%#ok<*EMCLS>
%#ok<*EMCA>
%#codegen

for n = 1:nargin
    if ~isa(varargin{n},'double')
        error(message('signal:chkinputdatatype:NotSupported'));
    end
end

% [EOF]

function plotpkmarkers(hLine, y)
%PLOTPKMARKERS  handler to offset peak markers by the size of the marker
%
%   This function is for internal use only.
%
%   If the figure zoom ActionPostCallback event cannot be overridden
%   warn user that we can't override and use (centered) circular markers
%   for peaks.
%
%   Otherwise, use inverted triangular markers and offset them by the size
%   of the marker, taking care to recompute the offsets upon size change
%   and figure zoom.

%   Copyright 2014 The MathWorks, Inc.

if ~isempty(y) && ~isempty(hLine) && ishghandle(hLine)
  hFig = ancestor(hLine,'figure');
  hZoom = zoom(hFig);  
  cbActionPostZoom = get(hZoom,'ActionPostCallback');
  cbZoom = createZoomCallback();
  if isempty(cbActionPostZoom) || isequal(char(cbActionPostZoom),char(cbZoom))
    % offset markers when figure zoom is invoked.
    if isempty(cbActionPostZoom)
      set(hZoom, 'ActionPostCallback', cbZoom);
    end
    
    % use solid inverted triangular markers
    set(hLine, 'Marker', 'v');
    set(hLine, 'MarkerFaceColor', get(hLine,'Color'));

    % create callback to offset the peak markers
    setappdata(hLine, 'YPos', y);
    cbSizeChange = createSizeChangeCallback(hLine);
    
    % offset markers on a SizeChange/SizeChanged event.
    e = events(hFig);
    eventName = e{strncmp(e,'SizeChange',10)};
    listener = matlab.ui.internal.createListener(hFig, eventName, cbSizeChange);
    setappdata(hLine, 'Listener', listener);

    %Offset the markers now.
    cbSizeChange(hLine);
  else
    % zoom post-action callback has already been overridden
    warning(message('signal:findpeaks:CantOffsetPeakMarkers'));
  end
end

function fcn = createSizeChangeCallback(hLine)
fcn = @(obj, evt) offsetPeakMarkers(hLine);

function fcn = createZoomCallback()
fcn = @(hFig, hAxes) offsetPeakMarkersByAxes(hFig, hAxes);

% offset all peak markers contained within an axes
function offsetPeakMarkersByAxes(~, evt)
if isfield(evt,'Axes') && ishghandle(evt.Axes)
  hLines = findall(evt.Axes,'Type','line','Tag','Peak');
  for i=1:numel(hLines)
    offsetPeakMarkers(hLines(i));
  end
end

% offset the markers in the line
function offsetPeakMarkers(hLine)
if ishghandle(hLine)
  % Fetch the data needed to compute the offset line
  y = getappdata(hLine, 'YPos');
  hAxes = ancestor(hLine, 'axes');
  yMarkerOffset = get(hLine,'MarkerSize');
  axesPos = getpixelposition(hAxes);
  yLim = get(hAxes,'YLim');

  % bump the line y data by the marker size
  yOffset = yMarkerOffset * diff(yLim) ./ axesPos(4);
  yOld = get(hLine, 'YData');
  yNew = y + yOffset;
  if ~isequaln(yOld, yNew)
    set(hLine,'YData', y + yOffset);
  end
end
```
