## Supplementary material for "Instantaneous phase of rhythmic behaviour under volitional control": Code and data files: fir1.m

```
% Copyright (C) 2000 Paul Kienzle <>
%
% This program is free software: you can redistribute it and/or modify
% it under the terms of the GNU General Public License as published by
% the Free Software Foundation, either version 3 of the License, or
% (at your option) any later version.
%
% This program is distributed in the hope that it will be useful,
% but WITHOUT ANY WARRANTY; without even the implied warranty of
% MERCHANTABILITY or FITNESS FOR A PARTICULAR PURPOSE. See the
% GNU General Public License for more details.
%
% You should have received a copy of the GNU General Public License
% along with this program; see the file COPYING. If not, see
% <https://www.gnu.org/licenses/>.

% -*- texinfo -*-
% @deftypefn  {Function File} {@var{b} =} fir1 (@var{n}, @var{w})
% @deftypefnx {Function File} {@var{b} =} fir1 (@var{n}, @var{w}, @var{type})
% @deftypefnx {Function File} {@var{b} =} fir1 (@var{n}, @var{w}, @var{type}, @var{window})
% @deftypefnx {Function File} {@var{b} =} fir1 (@var{n}, @var{w}, @var{type}, @var{window}, @var{noscale})
%
% Produce an order @var{n} FIR filter with the given frequency cutoff @var{w},
% returning the @var{n}+1 filter coefficients in @var{b}.  If @var{w} is a
% scalar, it specifies the frequency cutoff for a lowpass or highpass filter.
% If @var{w} is a two-element vector, the two values specify the edges of a
% bandpass or bandstop filter.  If @var{w} is an N-element vector, each
% value specifies a band edge of a multiband pass/stop filter.
%
% The filter @var{type} can be specified with one of the following strings:
% "low", "high", "stop", "pass", "bandpass", "DC-0", or "DC-1".  The default
% is "low" is @var{w} is a scalar, "pass" if @var{w} is a pair, or "DC-0" if
% @var{w} is a vector with more than 2 elements.
%
% An optional shaping @var{window} can be given as a vector with length
% @var{n}+1.  If not specified, a Hamming window of length @var{n}+1 is used.
%
% With the option "noscale", the filter coefficients are not normalized. The
% default is to normalize the filter such that the magnitude response of the
% center of the first passband is 1.
%
% To apply the filter, use the return vector @var{b} with the @code{filter}
% function, for example @code{y = filter (b, 1, x)}.
%
% Examples:
% @example
% freqz (fir1 (40, 0.3));
% freqz (fir1 (15, [0.2, 0.5], "stop"));  # note the zero-crossing at 0.1
% freqz (fir1 (15, [0.2, 0.5], "stop", "noscale"));
% @end example
% @seealso{filter, fir2}
% @end deftypefn

% FIXME: Consider using exact expression (in terms of sinc) for the
%        impulse response rather than relying on fir2.
% FIXME: Find reference to the requirement that order be even for
%        filters that end high.  Figure out what to do with the
%        window in these cases

function b = fir1(n, w, varargin)

  if nargin < 2 || nargin > 5
    error('wrong number of inputs');
  end

  % Assign default window, filter type and scale.
  % If single band edge, the first band defaults to a pass band to
  % create a lowpass filter.  If multiple band edges, the first band
  % defaults to a stop band so that the two band case defaults to a
  % band pass filter.  Ick.
  window  = [];
  scale   = 1;
  ftype   = (length(w)==1);

  % sort arglist, normalize any string
  for i=1:length(varargin)
    arg = varargin{i};
    if ischar(arg), arg=lower(arg);end
    if isempty(arg) continue; end  
    switch arg
      case {'low','stop','dc-1'},             ftype  = 1;
      case {'high','pass','bandpass','dc-0'}, ftype  = 0;
      case {'scale'},                         scale  = 1;
      case {'noscale'},                       scale  = 0;
      otherwise                               window = arg;
    end
  end

  % build response function according to fir2 requirements
  bands = length(w)+1;
  f = zeros(1,2*bands);
  f(1) = 0; f(2*bands)=1;
  f(2:2:2*bands-1) = w;
  f(3:2:2*bands-1) = w;
  m = zeros(1,2*bands);
  m(1:2:2*bands) = rem([1:bands]-(1-ftype),2);
  m(2:2:2*bands) = m(1:2:2*bands);

  % Increment the order if the final band is a pass band.  Something
  % about having a nyquist frequency of zero causing problems.
  if rem(n,2)==1 && m(2*bands)==1,
    warning("n must be even for highpass and bandstop filters. Incrementing.");
    n = n+1;
    if isvector(window) && isreal(window) && ~ischar(window)
      % Extend the window using interpolation
      M = length(window);
      if M == 1,
        window = [window; window];
      elseif M < 4
        window = interp1(linspace(0,1,M),window,linspace(0,1,M+1),'linear');
      else
        window = interp1(linspace(0,1,M),window,linspace(0,1,M+1),'spline');
      end
    end
  end

  % compute the filter
  b = fir2(n, f, m, [], 2, window);

  % normalize filter magnitude
  if scale == 1
    % find the middle of the first band edge
    % find the frequency of the normalizing gain
    if m(1) == 1
      % if the first band is a passband, use DC gain
      w_o = 0;
    elseif f(4) == 1
      % for a highpass filter,
      % use the gain at half the sample frequency
      w_o = 1;
    else
      % otherwise, use the gain at the center
      % frequency of the first passband
      w_o = f(3) + (f(4)-f(3))/2;
    end

    % compute |h(w_o)|^-1
    renorm = 1/abs(polyval(b, exp(-1i*pi*w_o)));

    % normalize the filter
    b = renorm*b;
  end

end

%!demo
%! freqz(fir1(40,0.3));
%!demo
%! freqz(fir1(15,[0.2, 0.5], 'stop'));  # note the zero-crossing at 0.1
%!demo
%! freqz(fir1(15,[0.2, 0.5], 'stop', 'noscale'));

%!assert(fir1(2, .5, 'low', @hanning, 'scale'), [0 1 0]);
%!assert(fir1(2, .5, 'low', "hanning", 'scale'), [0 1 0]);
%!assert(fir1(2, .5, 'low', hanning(3), 'scale'), [0 1 0]);

%!assert(fir1(10,.5,'noscale'), fir1(10,.5,'low','hamming','noscale'));
%!assert(fir1(10,.5,'high'), fir1(10,.5,'high','hamming','scale'));
%!assert(fir1(10,.5,'boxcar'), fir1(10,.5,'low','boxcar','scale'));
%!assert(fir1(10,.5,'hanning','scale'), fir1(10,.5,'scale','hanning','low'));
%!assert(fir1(10,.5,'haNNing','NOscale'), fir1(10,.5,'noscale','Hanning','LOW'));
%!assert(fir1(10,.5,'boxcar',[]), fir1(10,.5,'boxcar'));

%% Test expected magnitudes of passbands, stopbands, and cutoff frequencies

%!test
%! b = fir1 (30, 0.3);
%! h = abs (freqz (b, 1, [0, 0.3, 1], 2));
%! assert (h(1), 1, 1e-3)
%! assert (all (h(2:3) <= [1/sqrt(2), 3e-3]))

%!test
%! b = fir1 (30, 0.7, "high");
%! h = abs (freqz (b, 1, [0, 0.7, 1], 2));
%! assert (h(3), 1, 1e-3)
%! assert (all (h(1:2) <= [3e-3, 1/sqrt(2)]))

%!test
%! b = fir1 (30, [0.3, 0.7]);
%! h = abs (freqz (b, 1, [0, 0.3, 0.5, 0.7, 1], 2));
%! assert (h(3), 1, 1e-3)
%! assert (all (h([1:2, 4:5]) <= [3e-3, 1/sqrt(2), 1/sqrt(2), 3e-3]))

%!test
%! b = fir1 (50, [0.3, 0.7], "stop");
%! h = abs (freqz (b, 1, [0, 0.3, 0.5, 0.7, 1], 2));
%! assert (h(1), 1, 1e-3)
%! assert (h(5), 1, 1e-3)
%! assert (all (h(2:4) <= [1/sqrt(2), 3e-3, 1/sqrt(2)]))
```
