## Supplementary material for "Instantaneous phase of rhythmic behaviour under volitional control": Code and data files: fir2.m

```
% Copyright (C) 2000 Paul Kienzle <>
%
% This program is free software: you can redistribute it and/or modify
% it under the terms of the GNU General Public License as published by
% the Free Software Foundation, either version 3 of the License, or
% (at your option) any later version.
%
% This program is distributed in the hope that it will be useful,
% but WITHOUT ANY WARRANTY; without even the implied warranty of
% MERCHANTABILITY or FITNESS FOR A PARTICULAR PURPOSE. See the
% GNU General Public License for more details.
%
% You should have received a copy of the GNU General Public License
% along with this program; see the file COPYING. If not, see
% <https://www.gnu.org/licenses/>.

% -*- texinfo -*-
% @deftypefn  {Function File} {@var{b} =} fir2 (@var{n}, @var{f}, @var{m})
% @deftypefnx {Function File} {@var{b} =} fir2 (@var{n}, @var{f}, @var{m}, @var{grid_n})
% @deftypefnx {Function File} {@var{b} =} fir2 (@var{n}, @var{f}, @var{m}, @var{grid_n}, @var{ramp_n})
% @deftypefnx {Function File} {@var{b} =} fir2 (@var{n}, @var{f}, @var{m}, @var{grid_n}, @var{ramp_n}, @var{window})
%
% Produce an order @var{n} FIR filter with arbitrary frequency response
% @var{m} over frequency bands @var{f}, returning the @var{n}+1 filter
% coefficients in @var{b}.  The vector @var{f} specifies the frequency band
% edges of the filter response and @var{m} specifies the magnitude response
% at each frequency.
%
% The vector @var{f} must be nondecreasing over the range [0,1], and the
% first and last elements must be 0 and 1, respectively.  A discontinuous
% jump in the frequency response can be specified by duplicating a band edge
% in @var{f} with different values in @var{m}.
%
% The resolution over which the frequency response is evaluated can be
% controlled with the @var{grid_n} argument.  The default is 512 or the
% next larger power of 2 greater than the filter length.
%
% The band transition width for discontinuities can be controlled with the
% @var{ramp_n} argument.  The default is @var{grid_n}/25.  Larger values
% will result in wider band transitions but better stopband rejection.
%
% An optional shaping @var{window} can be given as a vector with length
% @var{n}+1.  If not specified, a Hamming window of length @var{n}+1 is used.
%
% To apply the filter, use the return vector @var{b} with the @code{filter}
% function, for example @code{y = filter (b, 1, x)}.
%
% Example:
% @example
% f = [0, 0.3, 0.3, 0.6, 0.6, 1]; m = [0, 0, 1, 1/2, 0, 0];
% [h, w] = freqz (fir2 (100, f, m));
% plot (f, m, ';target response;', w/pi, abs (h), ';filter response;');
% @end example
% @seealso{filter, fir1}
% @end deftypefn

function b = fir2(n, f, m, grid_n, ramp_n, window)

  if nargin < 3 || nargin > 6
    error('wrong number of inputs');
  end

  % verify frequency and magnitude vectors are reasonable
  t = length(f);
  if t<2 || f(1)~=0 || f(t)~=1 || any(diff(f)<0)
    error ('fir2: frequency must be nondecreasing starting from 0 and ending at 1');
  elseif t ~= length(m)
    error ('fir2: frequency and magnitude vectors must be the same length');
  % find the grid spacing and ramp width
  elseif (nargin>4 && length(grid_n)>1) || ...
         (nargin>5 && (length(grid_n)>1 || length(ramp_n)>1))
    error ('fir2: grid_n and ramp_n must be integers');
  end
  if nargin < 4, grid_n=[]; end
  if nargin < 5, ramp_n=[]; end

  % find the window parameter, or default to hamming
  w=[];
  if length(grid_n)>1, w=grid_n; grid_n=[]; end
  if length(ramp_n)>1, w=ramp_n; ramp_n=[]; end
  if nargin < 6, window=w; end
  if isempty(window), window=hamming(n+1); end
  if ~isreal(window) || ischar(window), window=feval(window, n+1); end
    if length(window) ~= n+1, error ('fir2: window must be of length n+1'); end

  % Default grid size is 512... unless n+1 >= 1024
  if isempty (grid_n)
    if n+1 < 1024
      grid_n = 512;
    else
      grid_n = n+1;
    end
  end

  % ML behavior appears to always round the grid size up to a power of 2
  grid_n = 2 ^ nextpow2 (grid_n);

  % Error out if the grid size is not big enough for the window
  if 2*grid_n < n+1
    error ('fir2: grid size must be greater than half the filter order');
  end

  if isempty (ramp_n), ramp_n = fix (grid_n / 25); end

  % Apply ramps to discontinuities
  if (ramp_n > 0)
    % remember original frequency points prior to applying ramps
    basef = f(:); basem = m(:);

    % separate identical frequencies, but keep the midpoint
    idx = find (diff(f) == 0);
    f(idx) = f(idx) - ramp_n/grid_n/2;
    f(idx+1) = f(idx+1) + ramp_n/grid_n/2;
    f = [f(:);basef(idx)]';

    % make sure the grid points stay monotonic in [0,1]
    f(f<0) = 0;
    f(f>1) = 1;
    f = unique([f(:);reshape(basef(idx),[],1)]');

    % preserve window shape even though f may have changed
    % JM change: MATLAB (2021b) errors if the basef points are not unique,
    % which might be often the case, add some numeric noise to basef
    for k = 1:numel(basef)-1
      if basef(k)==basef(k+1)
        basef(k) = basef(k)-eps;
        basef(k+1) = basef(k+1)+eps;
      end
    end

    m = interp1(basef, basem, f); %original

    % axis([-.1 1.1 -.1 1.1])
    % plot(f,m,'-xb;ramped;',basef,basem,'-or;original;'); pause;
  end

  % interpolate between grid points
  grid = interp1(f,m,linspace(0,1,grid_n+1)');
  % hold on; plot(linspace(0,1,grid_n+1),grid,'-+g;grid;'); hold off; pause;

  % Transform frequency response into time response and
  % center the response about n/2, truncating the excess
  if (rem(n,2) == 0)
    b = ifft([grid ; grid(grid_n:-1:2)]);
    mid = (n+1)/2;
    b = real ([ b([end-floor(mid)+1:end]) ; b(1:ceil(mid)) ]);
  else
    % Add zeros to interpolate by 2, then pick the odd values below.
    b = ifft([grid ; zeros(grid_n*2,1) ;grid(grid_n:-1:2)]);
    b = 2 * real([ b([end-n+1:2:end]) ; b(2:2:(n+1))]);
  end

  % Multiplication in the time domain is convolution in frequency,
  % so multiply by our window now to smooth the frequency response.
  % Also, for matlab compatibility, we return return values in 1 row
  b = b(:)' .* window(:)';

end

%% Test that the grid size is rounded up to the next power of 2
%% FIXME: test fails randomly on i386
%!xtest
%! f = [0 0.6 0.6 1]; m = [1 1 0 0];
%! b9  = fir2 (30, f, m, 9);
%! b16 = fir2 (30, f, m, 16);
%! b17 = fir2 (30, f, m, 17);
%! b32 = fir2 (30, f, m, 32);
%! assert ( isequal (b9,  b16))
%! assert ( isequal (b17, b32))
%! assert (~isequal (b16, b17))

%% Test expected magnitudes of passbands, stopbands, and cutoff frequencies
%!test
%! f = [0, 0.7, 0.7, 1]; m = [0, 0, 1, 1];
%! b = fir2 (50, f, m);
%! h = abs (freqz (b, 1, [0, 0.7, 1], 2));
%! assert (h(1) <= 3e-3)
%! assert (h(2) <= 1/sqrt (2))
%! assert (h(3), 1, 2e-3)

%!test
%! f = [0, 0.25, 0.25, 0.75, 0.75, 1]; m = [0, 0, 1, 1, 0, 0];
%! b = fir2 (50, f, m);
%! h = abs (freqz (b, 1, [0, 0.25, 0.5, 0.75, 1], 2));
%! assert (h(1) <= 3e-3)
%! assert (h(2) <= 1/sqrt (2))
%! assert (h(3), 1, 2e-3)
%! assert (h(4) <= 1/sqrt (2))
%! assert (h(5) <= 3e-3)

%!test
%! f = [0, 0.45, 0.45, 0.55, 0.55, 1]; m = [1, 1, 0, 0, 1, 1];
%! b = fir2 (50, f, m);
%! h = abs (freqz (b, 1, [0, 0.45, 0.5, 0.55, 1], 2));
%! assert (h(1), 1, 2e-3)
%! assert (h(2) <= 1/sqrt (2))
%! assert (h(3) <= 1e-1)
%! assert (h(4) <= 1/sqrt (2))
%! assert (h(5), 1, 2e-3)

%!demo
%! f=[0, 0.3, 0.3, 0.6, 0.6, 1]; m=[0, 0, 1, 1/2, 0, 0];
%! [h, w] = freqz(fir2(100,f,m));
%! subplot(121);
%! plot(f,m,';target response;',w/pi,abs(h),';filter response;');
%! subplot(122);
%! plot(f,20*log10(m+1e-5),';target response (dB);',...
%!      w/pi,20*log10(abs(h)),';filter response (dB);');

%!demo
%! f=[0, 0.3, 0.3, 0.6, 0.6, 1]; m=[0, 0, 1, 1/2, 0, 0];
%! plot(f,20*log10(m+1e-5),';target response;');
%! hold on;
%! [h, w] = freqz(fir2(50,f,m,512,0));
%! plot(w/pi,20*log10(abs(h)),';filter response (ramp=0);');
%! [h, w] = freqz(fir2(50,f,m,512,25.6));
%! plot(w/pi,20*log10(abs(h)),';filter response (ramp=pi/20 rad);');
%! [h, w] = freqz(fir2(50,f,m,512,51.2));
%! plot(w/pi,20*log10(abs(h)),';filter response (ramp=pi/10 rad);');
%! hold off;

%!demo
%! % Classical Jakes spectrum
%! % X represents the normalized frequency from 0
%! % to the maximum Doppler frequency
%! asymptote = 2/3;
%! X = linspace(0,asymptote-0.0001,200);
%! Y = (1 - (X./asymptote).^2).^(-1/4);
%!
%! % The target frequency response is 0 after the asymptote
%! X = [X, asymptote, 1];
%! Y = [Y, 0, 0];
%!
%! plot(X,Y,'b;Target spectrum;');
%! hold on;
%! [H,F]=freqz(fir2(20, X, Y));
%! plot(F/pi,abs(H),'c;Synthesized spectrum (n=20);');
%! [H,F]=freqz(fir2(50, X, Y));
%! plot(F/pi,abs(H),'r;Synthesized spectrum (n=50);');
%! [H,F]=freqz(fir2(200, X, Y));
%! plot(F/pi,abs(H),'g;Synthesized spectrum (n=200);');
%! hold off;
%! title('Theoretical/Synthesized CLASS spectrum');
%! xlabel('Normalized frequency (Fs=2)');
%! ylabel('Magnitude');
```
