## Supplementary material for "Instantaneous phase of rhythmic behaviour under volitional control": Code and data files: firls.m

```
% Copyright (C) 2006 Quentin Spencer <>
%
% This program is free software: you can redistribute it and/or modify
% it under the terms of the GNU General Public License as published by
% the Free Software Foundation, either version 3 of the License, or
% (at your option) any later version.
%
% This program is distributed in the hope that it will be useful,
% but WITHOUT ANY WARRANTY; without even the implied warranty of
% MERCHANTABILITY or FITNESS FOR A PARTICULAR PURPOSE. See the
% GNU General Public License for more details.
%
% You should have received a copy of the GNU General Public License
% along with this program; see the file COPYING. If not, see
% <https://www.gnu.org/licenses/>.

% -*- texinfo -*-
% @deftypefn  {Function File} {@var{b} =} firls (@var{n}, @var{f}, @var{a})
% @deftypefnx {Function File} {@var{b} =} firls (@var{n}, @var{f}, @var{a}, @var{w})
%
% FIR filter design using least squares method. Returns a length @var{n}+1
% linear phase filter such that the integral of the weighted mean
% squared error in the specified bands is minimized.
%
% The vector @var{f} specifies the frequencies of the band edges, normalized
% so that half the sample frequency is equal to 1.  Each band is specified by
% two frequencies, to the vector must have an even length.
%
% The vector @var{a} specifies the amplitude of the desired response at each
% band edge.
%
% The optional argument @var{w} is a weighting function that contains one
% value for each band that weights the mean squared error in that band.
%
% @var{a} must be the same length as @var{f}, and @var{w} must be half the
% length of @var{f}.  @var{n} must be even.  If given an odd value,
% @code{firls} increments it by 1.
%
% The least squares optimization algorithm for computing FIR filter
% coefficients is derived in detail in:
%
% I. Selesnick, "Linear-Phase FIR Filter Design by Least Squares,"
% http://cnx.org/content/m10577
% @end deftypefn

function coef = firls(N, frequencies, pass, weight, str)

if (nargin < 3 || nargin > 6)
  error('there is somethig wrong with the input arguments');
elseif (nargin == 3)
  weight = ones (1, length(pass)/2);
  str = [];
elseif (nargin == 4)
  if ischar (weight)
    str = weight;
    weight = ones (size (pass));
  else
    str = [];
  end
end
if length(frequencies) ~= length(pass)
  error('F and A must have equal lengths.');
elseif 2 * length(weight) ~= length(pass)
  error('W must contain one weight per band.');
elseif ischar(str)
  error('This feature is implemented yet');
end

% ensure both pass and frequency to be a row vector
if ~isrow(frequencies), frequencies = frequencies(:)'; end
if ~isrow(pass), pass = pass(:)'; end

N = 2*ceil(N/2);

M = N/2;
w = kron(weight(:), [-1; 1]);
omega = frequencies * pi;
i1 = 1:2:length(omega);
i2 = 2:2:length(omega);

% Generate the matrix Q
% As illustrated in the above-cited reference, the matrix can be
% expressed as the sum of a Hankel and Toeplitz matrix. A factor of
% 1/2 has been dropped and the final filter coefficients multiplied
% by 2 to compensate.
cos_ints = [omega; sin((1:N)' * omega)];
q = [1, 1./(1:N)]' .* (cos_ints * w);
Q = toeplitz(q(1:M+1)) + hankel(q(1:M+1), q(M+1:end));

% The vector b is derived from solving the integral:
%
%           _ w
%          /   2
%  b  =   /       W(w) D(w) cos(kw) dw
%   k    /    w
%       -      1
%
% Since we assume that W(w) is constant over each band (if not, the
% computation of Q above would be considerably more complex), but
% D(w) is allowed to be a linear function, in general the function
% W(w) D(w) is linear. The computations below are derived from the
% fact that:
%     _
%    /                          a              ax + b
%   /   (ax + b) cos(nx) dx =  --- cos (nx) +  ------ sin(nx)
%  /                             2                n
% -                             n
%
cos_ints2 = [omega(i1).^2 - omega(i2).^2; ...
  cos((1:M)' * omega(i2)) - cos((1:M)' * omega(i1))] ./ ...
  ([2, 1:M]' * (omega(i2) - omega(i1)));
d = [-weight .* pass(i1); weight .* pass(i2)];
b = [1, 1./(1:M)]' .* ( (kron(cos_ints2, [1, 1]) + cos_ints(1:M+1,:)) * d(:));

% Having computed the components Q and b of the  matrix equation,
% solve for the filter coefficients.
a = Q \ b;
coef = [ a(end:-1:2); 2 * a(1); a(2:end) ];
coef = coef(:)';
```
