## Supplementary material for "Instantaneous phase of rhythmic behaviour under volitional control": Code and data files: hamming.m

```
function w= hamming(M,mode)

if nargin<2
    mode='symmetric';
end
if strcmp(mode, 'periodic')
    M=M+1;
elseif strcmp(mode,'symmetric')==0;
    error('If provided, second parameter must be either periodic or symmetric')
end

w = .54 - .46*cos(2*pi*(0:M-1)'/(M-1));
```
