## Supplementary material for "Instantaneous phase of rhythmic behaviour under volitional control": Code and data files: hilbert.m

```
function y = hilberTr(x)
%Hilbert transform of input

   if nargin ~= 1
       usage("y = hilbert(x)"); 
   end
   if ~isreal(x)
       error("hilbert: requires real input vector"); 
   end
   transpose = size(x,1)==1;
   if transpose
       x=x.'; 
   end
   [r, c] = size(x);
   n=2^nextpow2(r);
   if r < n
       x = [ x ; zeros(n-r, c) ]; 
   end
   y = fft(x);
   y = ifft([y(1,:) ; 2*y(2:n/2,:) ; y(n/2+1,:) ; zeros(n/2-1,size(y,2))]);
   if r < n
       y(r+1:n,:) = []; 
   end
   if transpose
       y = y.'; 
   end
end
```
