## Supplementary material for "Instantaneous phase of rhythmic behaviour under volitional control": Code and data files: kaiser.m

```
% Copyright (C) 1995, 1996, 1997 Kurt Hornik <>
% Copyright (C) 2000 Paul Kienzle <>
%
% This program is free software; you can redistribute it and/or modify it under
% the terms of the GNU General Public License as published by the Free Software
% Foundation; either version 3 of the License, or (at your option) any later
% version.
%
% This program is distributed in the hope that it will be useful, but WITHOUT
% ANY WARRANTY; without even the implied warranty of MERCHANTABILITY or
% FITNESS FOR A PARTICULAR PURPOSE. See the GNU General Public License for more
% details.
%
% You should have received a copy of the GNU General Public License along with
% this program; if not, see <http://www.gnu.org/licenses/>.
%
% usage:  kaiser (L, beta)
%
% Returns the filter coefficients of the L-point Kaiser window with
% parameter beta.
%
% For the definition of the Kaiser window, see A. V. Oppenheim &
% R. W. Schafer, 'Discrete-Time Signal Processing'.
%
% The continuous version of width L centered about x=0 is:
%
%         besseli(0, beta * sqrt(1-(2*x/L).^2))
% k(x) =  -------------------------------------,  L/2 <= x <= L/2
%                besseli(0, beta)
%
% See also: kaiserord

function w = kaiser (L, beta)

if nargin<2
  beta = 0.5;
end


if (nargin < 1)
  help(mfilename);
elseif ~(isscalar (L) && (L == round (L)) && (L > 0))
  error ('kaiser:  L has to be a positive integer');
elseif ~(isscalar (beta) && (beta == real (beta)))
  error ('kaiser:  beta has to be a real scalar');
end % if

if (L == 1)
  w = 1;
else
  m = L - 1;
  k = (0 : m)';
  k = 2 * beta / m * sqrt (k .* (m - k));
  if m==0
    w = 1;
  else
    w = besseli (0, k) / besseli (0, beta);
  end
end % if

end

%!demo
%! % use demo('kaiserord');
```
