## Supplementary material for "Instantaneous phase of rhythmic behaviour under volitional control": Code and data files: resample.m

```
% Copyright (C) 2008 Eric Chassande-Mottin, CNRS (France) <>
%
% This program is free software: you can redistribute it and/or modify
% it under the terms of the GNU General Public License as published by
% the Free Software Foundation, either version 3 of the License, or
% (at your option) any later version.
%
% This program is distributed in the hope that it will be useful,
% but WITHOUT ANY WARRANTY; without even the implied warranty of
% MERCHANTABILITY or FITNESS FOR A PARTICULAR PURPOSE. See the
% GNU General Public License for more details.
%
% You should have received a copy of the GNU General Public License
% along with this program; see the file COPYING. If not, see
% <https://www.gnu.org/licenses/>.

% -*- texinfo -*-
% @deftypefn  {Function File} {[@var{y}, @var{h}] =} resample (@var{x}, @var{p}, @var{q})
% @deftypefnx {Function File} {@var{y} =} resample (@var{x}, @var{p}, @var{q}, @var{h})
% Change the sample rate of @var{x} by a factor of @var{p}/@var{q}.  This is
% performed using a polyphase algorithm.  The impulse response @var{h} of
% the antialiasing filter is either specified or either designed with a
% Kaiser-windowed sinecard.
%
% Ref [1] J. G. Proakis and D. G. Manolakis,
% Digital Signal Processing: Principles, Algorithms, and Applications,
% 4th ed., Prentice Hall, 2007. Chap. 6
%
% Ref [2] A. V. Oppenheim, R. W. Schafer and J. R. Buck,
% Discrete-time signal processing, Signal processing series,
% Prentice-Hall, 1999
% @end deftypefn

function  [y, h] = resample( x, p, q, h )

if nargin<3
  error('at least 3 inputs are needed');
end

if any([p q]<=0) || any([p q]~=floor([p q]))
  error('resample.m: p and q must be positive integers');
end

% simplify decimation and interpolation factors

great_common_divisor=gcd(p,q);
if (great_common_divisor>1)
  p = double (p) / double (great_common_divisor);
  q = double (q) / double (great_common_divisor);
else
  p = double (p);
  q = double (q);
end

% filter design if required

if (nargin < 4)
  
  % properties of the antialiasing filter
  
  log10_rejection = -3.0;
  stopband_cutoff_f = 1 / (2 * max (p, q));
  roll_off_width = stopband_cutoff_f / 10.0;
  
  % determine filter length
  % use empirical formula from [2] Chap 7, Eq. (7.63) p 476
  
  rejection_dB = -20.0*log10_rejection;
  L = ceil((rejection_dB-8.0) / (28.714 * roll_off_width));
  
  % ideal sinc filter
  
  t=(-L:L)';
  
  % JMS: The matlab implementation uses a firls filter
  %ideal_filter=2*p*stopband_cutoff_f*sinc(2*stopband_cutoff_f*t);
  Lx = 2*10*max(p,q) + 1; % this is according to the matlab default
  ideal_filter=firls( Lx-1, [0 2*stopband_cutoff_f 2*stopband_cutoff_f 1], [1 1 0 0]);
  
  % determine parameter of Kaiser window
  % use empirical formula from [2] Chap 7, Eq. (7.62) p 474
  
  if ((rejection_dB>=21) && (rejection_dB<=50))
    beta = 0.5842 * (rejection_dB-21.0)^0.4 + 0.07886 * (rejection_dB-21.0);
  elseif (rejection_dB>50)
    beta = 0.1102 * (rejection_dB-8.7);
  else
    beta = 0.0;
  end
  
  % JMS: beta = 5 by default in the matlab version, this overrules the
  % above
  beta = 5;
  
  % apodize ideal filter response
  
  %h=kaiser(2*L+1,beta).*ideal_filter;
  h=kaiser(Lx,beta).*ideal_filter(:);
  
end

% check if input is a row vector
isrowvector=false;
if size(x,1)==1 && size(x,2)>1
  x=x(:);
  isrowvector=true;
end

% check if filter is a vector
if ~isvector(h)
  error('resample.m: the filter h should be a vector');
end

Lx = size(x,1);
Lh = length(h);
L = ( Lh - 1 )/2.0;
Ly = ceil(Lx*p/q);

% pre and postpad filter response

nz_pre = floor(q-mod(L,q));
hpad = [zeros(nz_pre,1);h(:)];

offset = floor((L+nz_pre)/q);
nz_post = 0;
while ceil( ( (Lx-1)*p + nz_pre + Lh + nz_post )/q ) - offset < Ly
  nz_post = nz_post+1;
end
hpad = [hpad; zeros(nz_post,1)];

% filtering
if p==1 && q==1
  % no resampling needed.
  y = x;
else
  xfilt = upfirdn(x,hpad,p,q);
  y = xfilt(offset+1:offset+Ly,:);
end

if isrowvector
  y=y.';
end

%!test
%! N=512;
%! p=3; q=5;
%! r=p/q;
%! NN=ceil(r*N);
%! t=0:N-1;
%! tt=0:NN-1;
%! err=zeros(N/2,1);
%! for n = 0:N/2-1,
%!   phi0=2*pi*rand;
%!   f0=n/N;
%!   x=sin(2*pi*f0*t' + phi0);
%!   [y,h]=resample(x,p,q);
%!   xx=sin(2*pi*f0/r*tt' + phi0);
%!   t0=ceil((length(h)-1)/2/q);
%!   idx=t0+1:NN-t0;
%!   err(n+1)=max(abs(y(idx)-xx(idx)));
%! endfor;
%! rolloff=.1;
%! rejection=10^-3;
%! idx_inband=1:ceil((1-rolloff/2)*r*N/2)-1;
%! assert(max(err(idx_inband))<rejection);

%!test
%! N=512;
%! p=3; q=5;
%! r=p/q;
%! NN=ceil(r*N);
%! t=0:N-1;
%! tt=0:NN-1;
%! reject=zeros(N/2,1);
%! for n = 0:N/2-1,
%!   phi0=2*pi*rand;
%!   f0=n/N;
%!   x=sin(2*pi*f0*t' + phi0);
%!   [y,h]=resample(x,p,q);
%!   xx=sin(2*pi*f0/r*tt' + phi0);
%!   t0=ceil((length(h)-1)/2/q);
%!   idx=t0+1:NN-t0;
%!   reject(n+1)=max(abs(y(idx)));
%! endfor;
%! rolloff=.1;
%! rejection=10^-3;
%! idx_stopband=ceil((1+rolloff/2)*r*N/2)+1:N/2;
%! assert(max(reject(idx_stopband))<=rejection);

%!test
%! N=1024;
%! p=2; q=7;
%! r=p/q;
%! NN=ceil(r*N);
%! t=0:N-1;
%! tt=0:NN-1;
%! err=zeros(N/2,1);
%! for n = 0:N/2-1,
%!   phi0=2*pi*rand;
%!   f0=n/N;
%!   x=sin(2*pi*f0*t' + phi0);
%!   [y,h]=resample(x,p,q);
%!   xx=sin(2*pi*f0/r*tt' + phi0);
%!   t0=ceil((length(h)-1)/2/q);
%!   idx=t0+1:NN-t0;
%!   err(n+1)=max(abs(y(idx)-xx(idx)));
%! endfor;
%! rolloff=.1;
%! rejection=10^-3;
%! idx_inband=1:ceil((1-rolloff/2)*r*N/2)-1;
%! assert(max(err(idx_inband))<rejection);

%% Test integer-type arguments
%!test
%! N = 512;
%! f = 0.1;
%! x = sin (2*pi*f*[0:N-1]);
%! y1 = resample (x, 3, 2);
%! y2 = resample (x, uint8 (3), 2);
%! assert (y1, y2);
```
