## Supplementary material for "Instantaneous phase of rhythmic behaviour under volitional control": Code and data files: upfirdn.m

```
% Copyright (C) 2008   Sylvain Pelissier   <>
%
% This program is free software; you can redistribute it and/or modify
% it under the terms of the GNU General Public License as published by
% the Free Software Foundation; either version 2 of the License, or
% (at your option) any later version.
%
% This program is distributed in the hope that it will be useful,
% but WITHOUT ANY WARRANTY; without even the implied warranty of
% MERCHANTABILITY or FITNESS FOR A PARTICULAR PURPOSE.  See the
% GNU General Public License for more details.
%
% You should have received a copy of the GNU General Public License
% along with this program; If not, see <http://www.gnu.org/licenses/>.

% -*- texinfo -*-
% @deftypefn {Function File} {[@var{yout}] =} bohmanwin(@var{xin},@var{h}),@var{p},@var{q})
%	Upsample, filter and downsample a signal.
% @seealso{rectwin,  bartlett}
% @end deftypefn

function yout = upfirdn(xin,h,p,q)

if(nargin < 2)
  error('usage : yout = upfirdn(xin,h,p,q)');
end
	
if(nargin < 3)
	p = 1;
	q = 1;
end
	
if(nargin < 4)
	q = 1;
end
	
if(floor(p) ~= p || floor(q) ~= q || p < 1 || q < 1)
	error('p and q must be positive integer');
end
	
yout = upsample(xin,p);
yout = convn(yout, h).*p; % original was filter(h, 1, yout);
% the scaling with p is needed as per github issue 2085, causing the output
% to be scaled by the value of p, with this change, the compat/matlab
% versions will give an output that is about equal (scaled with about 0.9993)
yout = downsample(yout,q);
```
