## Supplementary material for "Instantaneous phase of rhythmic behaviour under volitional control": Code and data files: upsample.m

```
% Author: Paul Kienzle <> (2007)
% This program is granted to the public domain.

% -*- texinfo -*-
% @deftypefn  {Function File} {@var{y} =} upsample (@var{x}, @var{n})
% @deftypefnx {Function File} {@var{y} =} upsample (@var{x}, @var{n}, @var{offset})
% Upsample the signal, inserting @var{n}-1 zeros between every element.
%
% If @var{x} is a matrix, upsample every column.
%
% If @var{offset} is specified, control the position of the inserted sample in
% the block of @var{n} zeros.
% @seealso{decimate, downsample, interp, resample, upfirdn}
% @end deftypefn

function y = upsample(x, n, phase)
if nargin<2 || nargin>3, print_usage; end

if nargin<3
  phase = 0;
end

if phase > n - 1
  warning('This is incompatible with Matlab (phase = 0:n-1). See octave-forge signal package release notes for details.')
end

[nr,nc] = size(x);
if any([nr,nc]==1)
  y = zeros(n*nr*nc,1);
  y(phase + 1:n:end) = x;
  if nr==1, y = y.'; end
else
  y = zeros(n*nr,nc);
  y(phase + 1:n:end,:) = x;
end

%!assert(upsample([1,3,5],2),[1,0,3,0,5,0]);
%!assert(upsample([1;3;5],2),[1;0;3;0;5;0]);
%!assert(upsample([1,2;5,6;9,10],2),[1,2;0,0;5,6;0,0;9,10;0,0]);
%!assert(upsample([2,4],2,1),[0,2,0,4]);
%!assert(upsample([3,4;7,8],2,1),[0,0;3,4;0,0;7,8]);
```
