## Supplementary material for "Instantaneous phase of rhythmic behaviour under volitional control": Code and data files: nanmean.m

```
% NANMEAN provides a replacement for MATLAB's nanmean.
%
% For usage see MEAN.

function y = nanmean(x, dim)

if nargin<2
  N = sum(~isnan(x));
  y = nansum(x) ./ N;
else
  N = sum(~isnan(x), dim);
  y = nansum(x, dim) ./ N;
end

end % function
```
