## Supplementary material for "Instantaneous phase of rhythmic behaviour under volitional control": Code and data files: nanstd.m

```
% NANSTD provides a replacement for MATLAB's nanstd that is almost
% compatible.
%
% For usage see STD. Note that the three-argument call with FLAG is not 
% supported.

function Y = nanstd(varargin)

Y = sqrt(nanvar(varargin{:}));

end % function
```
