## Supplementary material for "Instantaneous phase of rhythmic behaviour under volitional control": Code and data files: nanvar.m

```
function y = nanvar(x,flag,dim)
% FORMAT: Y = NANVAR(X,FLAG,DIM)
% 
%    This file is intended as a drop-in replacement for Matlab's nanvar. It
%    originally forms part of the NaN suite:
%    http://www.mathworks.com/matlabcentral/fileexchange/6837-nan-suite/
%    and was modified to be compatible.

% -------------------------------------------------------------------------
%    author:      Jan Glscher
%    affiliation: Neuroimage Nord, University of Hamburg, Germany
%   
%    

if isempty(x)
	y = NaN;
	return
end

if nargin < 2 || isempty(flag)
	flag = 0;
end

if nargin < 3
	dim = find(size(x)~=1,1,'first');
	if isempty(dim)
		dim = 1; 
	end	  
end


% Find NaNs in x and nanmean(x)
nans = isnan(x);
avg = nanmean(x,dim);

% create array indicating number of element 
% of x in dimension DIM (needed for subtraction of mean)
tile = ones(1,max(ndims(x),dim));
tile(dim) = size(x,dim);

% remove mean
x = x - repmat(avg,tile);

count = size(x,dim) - sum(nans,dim);

% Replace NaNs with zeros.
x(isnan(x)) = 0; 


% Protect against a  all NaNs in one dimension
i = find(count==0);

if flag == 0
	y = sum(x.*x,dim)./max(count-1,1);
else
	y = sum(x.*x,dim)./max(count,1);
end
y(i) = i + NaN;
```
