## Supplementary material for "Instantaneous phase of rhythmic behaviour under volitional control": Code and data files: range.m

```
function range=range(x,dim)
% computes the range of the first argument along the dimension specified by the second argument
% x: input data sample
% dim: (integer scalar, optional; default:1)

if nargin < 2
    range = max(x) - min(x);
else
    range = max(x,[],dim) - min(x,[],dim);
end
```
