## Supplementary material for "Instantaneous phase of rhythmic behaviour under volitional control": Code and data files: wrapTo2Pi.m

```
function lambda = wrapTo2Pi(lambda)
%wrapTo2Pi Wrap angle in radians to [0 2*pi]
%
%   lambdaWrapped = wrapTo2Pi(LAMBDA) wraps angles in LAMBDA, in radians,
%   to the interval [0 2*pi] such that zero maps to zero and 2*pi maps
%   to 2*pi. (In general, positive multiples of 2*pi map to 2*pi and
%   negative multiples of 2*pi map to zero.)
%
%   See also wrapToPi, wrapTo180, wrapTo360.

% Copyright 2007-2008 The MathWorks, Inc.
% $Revision: 1.1.6.1 $  $Date: 2008/12/22 23:50:53 $

positiveInput = (lambda > 0);
lambda = mod(lambda, 2*pi);
lambda((lambda == 0) & positiveInput) = 2*pi;
```
