## Supplementary material for "Instantaneous phase of rhythmic behaviour under volitional control": Code and data files: wrapToPi.m

```
function a = wraptopi(a, a_center )
% function a = wrap(a,a_center)
%
% Wraps angles to a range of 2*pi.
% Inverse of Matlab's "unwrap", and better than wrapToPi ( which has
% redundant [-pi,pi])
% Optional input "a_center" defines the center angle.  Default is 0, giving
% angles from (-pi,pi], chosen to match angle(complex(-1,0)).  Maximum
% possible value is pi.

% T.Hilmer, UH
% 2010.10.18 version 2
%   removed code from version 1. Have not bug-checked second input
%   "a_center"

if nargin < 2, a_center = 0; end

% new way
a = mod(a,2*pi); % [0 2pi)

% shift
j = a > pi - a_center;
a(j) = a(j) - 2*pi;
j = a < a_center - pi;
a(j) = a(j) + 2*pi;
```
