## Supplementary material for "Instantaneous phase of rhythmic behaviour under volitional control": Code and data files: getPhaseToolsM.py

```
# -*- coding: utf-8 -*-
"""
Created on Fri Aug 18 15:15:00 2023

@author: L. lancia

mailto:
"""

import numpy as np
import copy
from PyEMD import EMD_matlab as EMDm
from scipy import signal, interpolate
from scipy.stats import zscore
from stochastic.processes.noise import FractionalGaussianNoise
from hurst import compute_Hc

def normalize_cycle_amp(sig,symmetric=1,threshNorm=1e-10,maxIterN=5,symN=2):
    """
    demodulate envelope peaks iteratively and using pchip interpolation.
    At each iteration, max and min are individuated, their absolute values are 
    interpolated to obtain an envelope. The signal is then divided by the envelope.
    If even one peak of the obtained signal is larger than 0, this is sent to a new iteration.
    Otherwise the procedure is stopped.
    
    input:
        sig: input signal
        symmetric (binary, optional): if it is 0 the signal's cycles are not centered around 0. 
            This is solved by applying one iteration of sifting.
        threshNorm (positive real, optional): threshold to determine if peaks are different from 1/-1
        maxIterN (positive integer, optional): maximum number of iterations
        symN (positive integer, optional): number of extrtema to be added at the beginning and at the
              end to regularize boundary conditions before interpolation.
    output:
        demodulated signal
    """
    normSig=copy.copy(sig)

    if symmetric==0:
        # emdObj = EMDm.EMD(DTYPE=np.float32,MAX_ITERATION=2,FIXE = 1 )
        emdObj = EMDm.EMD()
        emdObj.MAX_ITERATION=2
        emdObj.FIXE = 1
        normSig=emdObj.emd(normSig,maxImf=2)[0]
        # normSig,_=emd.sift.get_next_imf(normSig,
        #         stop_method='rilling',max_iters=1000)
        #tmpIMFs=emdObj(resid+m[:, nn])
        normSig=normSig[0]#[:,0]
        
    newIdxs=np.arange(np.size(normSig)) #input signal's time steps 

    iter_val=1 #intitialize error value
    niter=0 #current iteration number
    
    while niter<maxIterN and iter_val>threshNorm:
        niter=niter+1

        pksLoc,_=signal.find_peaks(normSig)
        valLoc,_=signal.find_peaks(-normSig)
        
        if len(pksLoc)+len(valLoc)<3:
            return normSig/np.ptp(normSig)
        
        max_pos, max_val = pksLoc, normSig[pksLoc]
        min_pos, min_val = valLoc, normSig[valLoc]
        
        emdObj1=EMDm.EMD()
        # max_extrema, min_extrema= emdObj1._prepare_points_simple(newIdxs, normSig, 
        #                        max_pos, max_val, min_pos, min_val)
        max_extrema, min_extrema= emdObj1.preparePoints( normSig, newIdxs,
                                    max_pos, max_val, min_pos, min_val)
        extrIdxs=np.hstack((min_extrema[0],max_extrema[0]))
        extrVals=np.hstack((min_extrema[1],max_extrema[1]))
        
        extrSortIdxs=np.argsort(extrIdxs)
        extrIdxs=extrIdxs[extrSortIdxs]
        extrVals=extrVals[extrSortIdxs]
        
        refSigf=interpolate.PchipInterpolator(extrIdxs, np.abs(extrVals))
        refSig=refSigf(newIdxs)
    
        iter_val = np.abs(refSig.sum() - refSig.shape[0])

        
        normSig=normSig/ refSig 
        
    
    return normSig

def do_mask_sift(resid,CF=None,nMasks=4,ampCoeff=None,frqEst='phi',emdObj=None):

    """
    do masked sift to obtain one IMF from the input signal
     input:
         resid (optional, real valued 1D signal): input signal
         CF (optional, positive real): normalized (freq/sampling rate) masking frequency
         nMasks (optional, positive integer): number of masking signals to be used
         ampCoeff (optional, positive real): multiplicative mask amplitude coefficient 
         emdObj (optional, initialized emd object): Object of the EMD class parameterized 
                 to do the sifting (default: MAX_ITERATION=10) 

     output:
         IMF: (dimensionless numpy array): obtained signal
         CF: (positive real): frequency used for masking
         m[:, 0] (one dimensional np array): first masking signal used
         
    """

    if ampCoeff is None:
       ampCoeff=2*4*np.std(resid)
    if emdObj is None:
        # emdObj=EMDm.EMD(DTYPE=np.float32,MAX_ITERATION=10,FIXE = 1 )
        emdObj = EMDm.EMD()
        emdObj.MAX_ITERATION=10
        
    if CF is None or (np.size(CF)==1 and CF==0):
        # emdObj1= EMDm.EMD(DTYPE=np.float32,MAX_ITERATION=2,FIXE = 1 )
        emdObj1 = EMDm.EMD()
        emdObj1.MAX_ITERATION=1
        
        IMF0=emdObj1.emd(resid,maxImf=2)[0][0]
        IMF0=normalize_cycle_amp(IMF0) # refined amplitude normalization
        if frqEst=='phi':           # if frequency is to be estimated via phase derivative
            IPH=np.angle(signal.hilbert(IMF0)) # phase
            Ifreq=np.diff(np.unwrap(IPH));         
            IPH[Ifreq<0]=np.nan
            CF=np.nanmean(np.diff(np.unwrap(IPH))/(2*np.pi))   # normalized freq
        else:
            def zci(v): # define function to get zero crossings
                return np.argwhere((v[:]*np.roll(v[:],-1))<=0)
            CF=np.ceil((np.size(zci(IMF0))/2)/np.size(IMF0) ) # normalized freq
                
    # Create time matrix including mask phase-shifts
    t = np.repeat(np.arange(resid.shape[0])[:, np.newaxis], nMasks, axis=1)
    phases = np.linspace(0, (2*np.pi), nMasks+1)[:nMasks]
    # Create masks
    zf = CF * 2 * np.pi
    m =ampCoeff * np.sin(zf * t + phases)

        
    IMFs=np.zeros((resid.shape[0],nMasks))
    
    for nn in np.arange(nMasks):
        # tmpIMFs,_=emd.sift.get_next_imf(resid+m[:, nn],
        #         stop_method='rilling', rilling_thresh=[0.05,0.5,0.05],max_iters=1000)
        tmpIMFs=emdObj.emd(resid+m[:, nn],maxImf=2)[0]
        IMFs[:,nn]=tmpIMFs[0]#[:,0]#
        
    return np.sum(IMFs - m,axis=1)/nMasks, CF, m[:, 0]

def quadAngle(IMFin,hilbQuad=0):
    """ 
    computes the instantaneous phase of the columns of IMFin via the direct quadrature method
    Input:
        IMFin: array of signals (each column is a different signal)
        hilbQuad: toggle 0/1: if 1, hilbert quadrature (mask = sign(hilbert(data))),
            otherwise direct quadrature (mask= ((np.diff(data)>0) * -2) + 1;mask =np.append(mask,mask[-1]))
    Output:
        array of phase values
    """
    if len(np.shape(IMFin))==1:
        npt=np.size(IMFin)
        ncol=1
    else:
        npt=np.shape(IMFin)[0]
        ncol=np.shape(IMFin)[1]
        
    #1----- Initialize and flip data if needed 
    flipped=0;
    if (ncol > npt):
        flipped=1
        IMFin=IMFin.T
        npt= np.shape(IMFin)[0]
        ncol=np.shape(IMFin)[1]
        
    
    #2.Calculate the quadrature value--loop start
     
    #quadrature=np.complex((np.shape(IMFin)[0],ncol))
    quadrature = np.empty((np.shape(IMFin)[0],ncol)).astype(complex)
    for i in np.arange(ncol):
        if ncol==1:
            data=IMFin
        else:
            data = IMFin[:,i]
           
    #3.add mask-for positive/negative signal    
            #creates a mask for the signal 
            #the 1st & 2nd Q = 1 
            #and the 3rd & 4th Q = -1
        if hilbQuad==0:  
            mask = ((np.diff(data)>0) * -2) + 1;
            mask = np.append(mask,mask[-1])
        else:
            P=signal.hilbert(data)
            mask=np.sign(np.imag(P));
        
    #4.the calculation of quadrature value
        y = np.real(np.sqrt(1-data**2))
        
    #5.multiplying by mask flips data in 3rd & 4th quadrature
        q = y * mask
        quadrature[:,i] = data+1j* q
    
    quadAngle=np.angle(quadrature)
    if flipped==1:
        quadAngle = quadAngle.T;
        
    return quadAngle

def get_omega(PHIin,sr,m=16,n=5):
    """
    given an input sequence of phase values it computes the median frequency 
    
     input:
        PHIin: input phase signal
        sr: phase sampling rate
        m: (positive integer; optional, default: 16): length of the Savitzky-Golay differentiator.  
        n: (positive integer; optional, default: 5): order of the Savitzky-Golay differentiator.  
        
     output:
        outFreq: median frequency
    """
           
    PHIu=np.unwrap(PHIin); # unwrap phase
    
    # get freqs via differentiation
    Freqs=signal.savgol_filter(
        PHIu, m, polyorder=n, deriv=1, delta= 1/sr,axis=0, mode='interp')
    # here there is a difference with the Matlab version 
    # (where the output of the filtering is truncated) because
    # the truncated output of filtering is extended (to match 
    # the input length) via polinomial fitting (see help of savgol_filter)
        
    return np.median((Freqs/sr)/(2*np.pi))


def maskEMD(sigIn,sr, maxIMFn=10,nMasks=22,ampCoeff=2,m=16,n=5,stdRatioThresh=1e-6):
    """
    masked EMDm.EMD
    
    input:
         signal: input signal
         sr (positive real): sampling frequency in Hz
         maxIMFn (positive integer; optional, default=10): maximum number of IMFs
         if equal to 0, the procedure will be stopped when the obtained 
         IMF contains no extrema.
         nMasks (positive integer; optional, default= 22 ): number of mask signal used in sifting 
         ampCoeff ( positive real, optional, default= 2): coefficient determining the amplitude of the masks as a
                proportion of 4*std(signal), which is meant to be a rough estimate of the signal's 
                range from the std if the signal's values are normally distributed.
         m (positive integer; optional, default= 16): length of the Savitzky-Golay differentiator.  
         n (positive integer; optional, default= 5): order of the Savitzky-Golay differentiator.  
         stdRatioThresh (positive real; optional, default= 1E-6): threshold for EMD convergence
    output:
        outIMFs: the extracted IMFs
        IMFfreqs: the estimated frequencies of the extracted IMFs 
    """
    nPks=1  #initialize variable containing the unmber of extrema in the current IMF
    resid=sigIn  #initialize input signal for masked sifting 
    outIMFs=np.zeros((np.size(sigIn),maxIMFn))  #initialize matrix containing IMFs
    IMFfreqs=np.zeros(maxIMFn); #initialize vector containing IMFs frequencies
    nn=0 # number of iterations so far
    emdObj = EMDm.EMD()
    emdObj.MAX_ITERATION=2
    emdObj.FIXE = 1
    
    emdObj1 = EMDm.EMD()
    emdObj1.MAX_ITERATION=10
    
    stdRatio=1
    while nPks>0 and nn <maxIMFn-1 and stdRatio>stdRatioThresh:
        

        myMode=emdObj.emd(resid,maxImf=2)[0]
        # myMode,_=emd.sift.get_next_imf(resid,
        #         stop_method='fixed', max_iters=1)
        myMode=myMode[0]#[:,0]
        
        if ampCoeff is not None and nn==0:
            thisCoeff=ampCoeff*4*np.std(myMode) # set amplitude of mask signal (it is based on the first IMF for denoising purposes)
    
        elif ampCoeff is None:
            thisCoeff=None; # the empty values triggers the use of the default value in mask_sift
                      # this corresponds to twice the range of the input signal
        myModeN=normalize_cycle_amp(myMode)
        PHI=np.angle(signal.hilbert(myModeN))
        
        sigFreq=get_omega(PHI,sr,m,n)

        outIMFs[:,nn],_,_=do_mask_sift(resid,sigFreq,nMasks,thisCoeff,emdObj=emdObj1); # do masked sifting
        
        
        myModeN=normalize_cycle_amp(outIMFs[:,nn],maxIterN=10) # apply refined amplitude normalization 

        PHIcNew=np.angle(signal.hilbert(myModeN));#hilbert phase (better suited than quadr. to infer local frequancy)

        PHIu=np.unwrap(PHIcNew); # unwrap phase
        medF=np.median((np.diff(PHIu)/(2*np.pi))); # compute median frequency

        IMFfreqs[nn]=medF; # store median frequency
        thisStd=np.std(resid)
        resid=resid-outIMFs[:,nn]# update input signal for next application of masked sifting 
        nPks=np.size(signal.find_peaks(resid)[0]); # count number of peaks
        nPks=nPks+np.size(signal.find_peaks(-resid)[0]); # get number of extrema
        stdRatio=np.std(resid)/thisStd
        if (ampCoeff is not None) and nn==0:
            thisCoeff=ampCoeff*4*np.std(outIMFs[:,nn])
        
        nn+=1
    if nn<maxIMFn-1:
        outIMFs=outIMFs[:,:nn]
        IMFfreqs=IMFfreqs[:nn]
        
    return outIMFs,IMFfreqs


def mEMDdenoise(data,sr,nMasks=22,ampCoeff=2,alpha=0.05,nReps=100,m=16,n=5):
    """
    remove noise from signal in Data sampled at freq. sr 
    input:
       data: input signal
       sr (positive real): sampling rate
       nMasks (positive integer; optional, default: 22): number of mask signal for masked EMD application .  
       ampCoeff (positive real; optional, default: 2): amplitude gain of the mask signals 
           (it is multiplied by 4sd(IMF1), where IMF1 is the first IMF or the centered data at the first iteration ).  
       nReps (positive integer; optional, default: 100): number of repetitions of simulated random processes .  
       m (positive integer; optional, default: 16): length of the Savitzky-Golay differentiator.  
       n (positive integer; optional, default: 5): order of the Savitzky-Golay differentiator.  

    output:
       filtered: filtered signal
       imf: all signal's IMFs
       imfF: frequencies of extracted IMFs
       filteredidx: indexes of IMFs composing the filtered signal
       noiseStd: standard deviation of the estimated noise (sum of the random components).
"""
    if np.diff(np.shape(data))>0: # be sure that the input signal is a column vector
        data=data.T
    
    data=zscore(data)
    
    lenData=np.size(data)
    
    #apply masked EMD  to get the signal's IMFs
    [imf,imfF]=maskEMD(data,sr,10,nMasks,ampCoeff,m,n)
    
    # H0=hurst_exponent(zscore(imf[:,0])) # estimate nois hurst exponent through the first IMF
    H0,_,_=compute_Hc(zscore(imf[:,0]))
    prdata=FractionalGaussianNoise(hurst=H0, t=1) # generate noise simulation object
    
    nObs = np.shape(data)[0] # signal length
    nImf = np.shape(imf)[1] #number of IMFs


    # produce nReps random signals with Hurst exponent set to H0 
    f=np.empty((nObs,nReps))
    for i in np.arange(nReps):
        f[:,i] = prdata.sample(lenData)
    
    noiseLev=np.empty((nReps,nImf));# build storage for energy of the IMFs extracted from the random signals

    # emdObj=EMDm.EMD(DTYPE=np.float32,MAX_ITERATION=nImf,FIXE = 1 ) # build EMD object
    emdObj = EMDm.EMD()
    emdObj.MAX_ITERATION=nImf
    #emdObj.FIXE = nImf-1
    
    for i in np.arange(np.shape(noiseLev)[0]):
       rndImfs=emdObj.emd(f[:,i])[0]# apply EMD
       rndImfsArr=np.empty((np.size(rndImfs[0]),len(rndImfs)))# need to transform the dictionary into an array
       for h in rndImfs.keys():
           rndImfsArr[:,h]=rndImfs[h]
       # rndImfs,_=emd.sift.get_next_imf(f[:,i],
       #          stop_method='rilling', rilling_thresh=[0.05,0.5,0.05],max_iters=1000)
       #rndImfs=rndImfs[:,0]
       noiseLev[i,0:nImf] = np.sum(rndImfsArr**2,axis=0)[0:nImf]/nObs#compute energy of ectracted IMFs

    myQuants=np.quantile(noiseLev, [alpha, 1-alpha],axis=0).T
    filteredidxs = np.zeros(nImf)# build storage for chosen IMFs' indexes
    filtered = np.zeros(nObs)#initialize filtered signal
    energy = np.zeros(nImf)# build storage for IMFs' energy values

    for i in np.arange(nImf):# compute the energy of each IMF extraced from real data
        energy[i] = np.sum(imf[:, i]**2, axis=0)/nObs
        if (energy[i] >= myQuants[i,1]):# if test passed 
            filteredidxs[i] = i # store the index of the selected mode
            filtered = filtered + imf[:,i]# add the selected mode to the fitlered signal

    noiseStd=np.std(data-filtered); # compute rel std of the random component
    
    #indexes of statistically significant IMFs
    filteredidxs = filteredidxs[np.where(filteredidxs>0)]
    
    return filtered, imf, imfF, filteredidxs, noiseStd

    
def getPhaseMask(sigIn,sr,m=16,n=5,nMasks=22,ampCoeff=2, quadMethod=['h','h'], threshs=[1e-10,1e-10]):
    """
   performs EMD via Masked Sifting
    input: 
        sigIn: input signal
        sr: sampling rate
        m (positive integer; optional, default: 16): number of filtered points for Savitzky-Golay differentiator 
        n (positive integer; optional, default: 5): polinomial order of the differentiator
        nMasks (positive integer; optional, default: 22): number of masks used to extract the independent mode function via masked EMD
        ampCoeff (positive real; optional, default: 2): amplitude of the mask signals used for sifting (optional,
                 default= twice a rough estimate of the range of the signal sent to 
                 masked sifting). Non default values are meant to be used in denoising 
                 applications.
        quadMethod (a string or a cell of two strings, default: ['h','h']). Method to be used in the
                computation of the quadrature signal 'h' stands for Hilbert and 'q'. If two 
                strings are provided a different method can be adopted in in the first or 
                the second part of the algorithm.  
        threshs (scalar or vector of two positive real values close to zero, default: [1E-10,1E-10]):
            threshold for refined amplitude normalization. If two values, different thresholds
            can be used in the two parts of the algorithm.  
    output:
        PHI: instantaneous phase
        IMF: signal obtained from Masked Sifting
        PHIc: instantaneous phase of the centered signal
        centeredSig: centred signal
        mask: first masking signal used in Masked Sifting
    """
    
    if type(quadMethod)==str:
        quadMethod=[quadMethod,quadMethod]
    if type(threshs)==int or type(threshs)==float:
        threshs=[threshs,threshs]
    
    emdObj = EMDm.EMD()
    emdObj.MAX_ITERATION=2
    emdObj.FIXE = 1
    IMF0=emdObj.emd(sigIn,maxImf=2)[0]
    #
    # IMF0,_=emd.sift.get_next_imf(sigIn,
    #          stop_method='fixed', max_iters=1)
    IMF0=IMF0[0]#[:,0]
    centredSig=normalize_cycle_amp(IMF0)
    
    if quadMethod[0]=='q':
        PHI=quadAngle(centredSig)[:,0] 
    elif quadMethod[0]=='h':
        PHI=np.angle(signal.hilbert(centredSig))
    elif quadMethod[0]=='hq':
            PHI=quadAngle(centredSig,hilbQuad=1)[:,0] 
    else:
        raise('only methods avialable to get the quadrature signal are Hilbert: \
              ''h'', direct quadrature: ''q'' or Hilbert/direct quadrature ''hq''')

    
    sigFreq = get_omega(PHI,sr,m,n)
    
    if ampCoeff is not None:
        ampCoeff=ampCoeff*4*np.std(sigIn)
    
    emdObj.MAX_ITERATION=10
 
    IMF1,_,mask=do_mask_sift(sigIn, sigFreq,nMasks,ampCoeff,emdObj)#get signal via masked sifting
    
    IMF=normalize_cycle_amp(IMF1); #refined amplitude normalization

    if quadMethod[1]=='q':
        newPHI=quadAngle(IMF)[:,0] # quadrature phase estimation (here we need to be unbiased so we may not want to use Hilbert)
    elif quadMethod[1]=='h':
        newPHI=np.angle(signal.hilbert(IMF))
    elif quadMethod[1]=='hq':
            PHI=quadAngle(centredSig,hilbQuad=1)[:,0] 
    else:
        raise('only methods avialable to get the quadrature signal are Hilbert: \
              ''h'', direct quadrature: ''q'' or Hilbert/direct quadrature ''hq''')

    return newPHI, IMF, PHI, centredSig,mask

# 
```
