## Supplementary material for "Instantaneous phase of rhythmic behaviour under volitional control": Code and data files: paper_analyses.m

```
% Script to reproduce the computations in the results section of the paper
% Select one example from the following list by setting the value of the
% variable exampleN at line 16 to the corresponding example number.
%
% 1: gait data
% 2: tongue, lip and jaw data
% 3: tongue, lip and jaw data with noise
% 4: knee flexion data
% 5: vocal folds vibration data

% Leonardo Lancia 14/10/2023
% mailto:

clear all;

exampleN=5; % select one example with an integer from 1 to 6

addpath('./gen_utils');
addpath(genpath('./get_phase')); 

%Savitzky-Golay differentiator parameters:
m = 16;      % number of filtered points (FIR order)
n = 5;      % approximation polynomial order

phaseMethod='h'; % set method to extract quadrature signal. Possible methods: 'h','q','hq' (the last one is expereimental, do not use)
threshs=1e-10; %threshold for the (refined) amplitude normalization operation 
ampCoeff=2; %coefficient modulating the amplitude of the masking signal
nMasks=22; % number of masking signals for masked sifting 
nRandNoiseRep=100; % number of repetitions of random processes in the denoising algorithm


switch exampleN
    case 1  % gait example, sampling rate = 200 Hz, 
            % The data set includes the ground reaction forces as well as the marker positions of
            % the markers that were placed on the shoes above the first (FM1), second (FM2) and
            % fifth metatarsal (FM5) heads and on the aspect of the Achilles tendon insertion on
            % the calcaneus (FCC) for dynamic tracking. Data low pass filtered at % 6hz
        
        dataIn=load('./data./GaitPhase_Database_example.mat');
        dataIn=dataIn.my_struct;
        sr=200;
        sig=dataIn.L_FCC_z';
        sigT=[1:length(sig)]./sr;% time stamps
        options.MAXITERATIONS=10; % set maximum sifting iteration to 10
        Decomp=emd(sig,options)'; % get classic decomposition

        [PHI,newIMF,PHI0,centSig,mask]=getPHImask(sig,sr,m,n,nMasks,ampCoeff,phaseMethod,threshs);
        figure
        axx=tight_subplot(4,2,0.04,0.08); %initialize subplots
        
        axes(axx(1));
        plot(sigT,sig,'k'); % plot signal
        xticks([])
        title('\textbf{Heel height}','interpreter','latex')
        axes(axx(3));
        plot(sigT,wrapTo2Pi(unwrap(angle(hilbert(zscore(sig))))),'k'); % plot hilbert phase of z-scored signal
        setPhaseAx(gca) % set y ticks
        xticks([])
        title('\textbf{Hilbert phase of Heel height}','interpreter','latex')
        axes(axx(5));
        plot(sigT,wrapTo2Pi(unwrap(angle(hilbert(normalize_cycle_amp(Decomp(:,1)))))),'k');% plot hilber phase of amplitude normalized first IMF from EMD
        setPhaseAx(gca)
        xticks([])
        title('\textbf{Hilbert phase of $$IMF_{1}$$}','interpreter','latex')
        axes(axx(7));
        plot(sigT,wrapTo2Pi(unwrap(angle(hilbert(normalize_cycle_amp(Decomp(:,2)))))),'k');% plot hilber phase of amplitude normalized second IMF from EMD
        setPhaseAx(gca)
        title('\textbf{Hilbert phase of $$IMF_{2}$$}','interpreter','latex')
        xlabel('Time (sec)')

        axes(axx(2));
        plot(sigT,newIMF,'k');% plot signal after masked sifting
        xticks([])
        title('\textbf{Output of Masked Sifting ($$\hat{x}$$)}','interpreter','latex')

        axes(axx(4));
        IMFF=normalize_cycle_amp(newIMF,[],[],5);%refined amplitude normalizatin algo 

        plot(sigT,wrapTo2Pi(unwrap(angle(hilbert(IMFF)))),'k');% plot phase of the obtained signal
        setPhaseAx(gca)
        xlabel('Time (sec)')
        title('\textbf{Hilbert phase of $$\hat{x}$$}','interpreter','latex')

        linkaxes(axx,'x')
         xlim([2500/sr,5500/sr])
        delete([axx(6),axx(8)])
    
    case 2
        
        artDimN=[16,18,20];%select relevant channels: ttip 16, llip 18, jawY 20 
        cycle1=[3150,3350];%boundaries of slow cycle example
        cycle2=[2479,2645];%boundaries of fast cycle example
        pathIn='./data./cl_pata_2.mat';
        dataIn=load(pathIn); %load data 
        sigIn=dataIn.data(1:end,:);%(:,artDimN);%get jawY 20 ,ulip 17, llip 18, ttip 16 sensor (cf. strmatch('lev-sY ',dataIn.descriptor,'exact'),'lev-sY','lev-iY ','lang-aY' )
        sr=dataIn.samplerate;% sampling rate
        sigT=[1:length(sigIn)]./sr;% time steps

        %low pass filter parameters
        hiCut=20; %Max allowed frequency
        filtLen=1000;%filter order
        hiCutNorm=hiCut./( sr/2 ); %normalized cut off for the low-pass filter
        b1=fir1(filtLen,hiCutNorm,'low'); %filter coeffs
        filtSig=filter(b1,1,sigIn); %apply filter to input sig.
        filtSig=zscore(filtSig);%at least now it oscillates around 0 like a real sinusoid.
        sigInf=filtSig(filtLen/2+1:end,:); %correct filter delay
        sigTf=sigT(1:length(sigInf)); %remove exceding samples' time points
        
        sigInf=sigInf(1:end,:);% remove artefacts at the onset
        sigTf=sigTf(1:end);% remove artefacts at the onset
        myData=[sigInf(:,artDimN(1)),sigInf(:,artDimN(2)),sigInf(:,artDimN(3))];%extract relevant channels
        
        txtY=[-1.2,-2.2,-1.5];
        txtX=8;
        artLabels={'TTIP','LLIP','JAW'};
        fSize=8;
        
        figure;
        
        subplot(4,4,[1 2 3 5 6 7 9 10 11 13 14 15]);%initialize subplots
        haa=stackedplot(sigTf,myData(:,:),'DisplayLabels',{'' '' '' } ,'color','k');% plot all sequence
        ax = flipud(findobj(haa.NodeChildren, 'Type','Axes'));
        for h =1:length(artLabels)
            myLims=get(ax(h),'ylim');
            text(ax(h),txtX,myLims(1)+diff(myLims)/6,artLabels{h});
        end
        xlabel('Time (sec)')
        
        subplot(4,4,[4 8]);% plot slow cycle
        h=stackedplot(sigTf(cycle1(1):cycle1(2)),myData(cycle1(1):cycle1(2),:),'DisplayLabels',{'' '' '' } ,'color','k');
        ax = flipud(findobj(h.NodeChildren, 'Type','Axes'));
        ax(end).XTick=[];
        for h =1:length(artLabels)
            myXLims=get(ax(h),'xlim');
            myLims=get(ax(h),'ylim');
            text(ax(h),myXLims(1)+diff(myXLims)/100,myLims(2)-diff(myLims)/6,artLabels{h},'FontSize',fSize);
        end
        
        subplot(4,4,[12 16]);% plot fast cycle
        h=stackedplot(sigTf(cycle2(1):cycle2(2)),myData(cycle2(1):cycle2(2),:),'DisplayLabels',{'' '' '' } ,'color','k');
        ax = flipud(findobj(h.NodeChildren, 'Type','Axes'));
        for h =1:length(artLabels)
            myXLims=get(ax(h),'xlim');
            myLims=get(ax(h),'ylim');
            text(ax(h),myXLims(1)+diff(myXLims)/100,myLims(2)-diff(myLims)/6,artLabels{h},'FontSize',fSize);
        end
        xlabel('Time (sec)');
        
        myPhases=zeros(size(myData)); %initialize matrix containing phase values
        for dataN=1:size(myData,2)
            [PHI,newIMF,PHI0,centeredSig,mask]=getPHImask(myData(:,dataN),sr,m,n,nMasks,ampCoeff,phaseMethod,threshs);
            myPhases(:,dataN)=PHI;
        end
        
        figure;
        stackedplot(sigTf,[myData(:,3),myPhases(:,:)],'DisplayLabels',{'' '' '' '' } ,'color','k');
        xlabel('Time (sec)');
        
    case 3
        
        pathIn='./data./cl_pata_2.mat';
        dataIn=load(pathIn); %load data 
        artDimN=[16,18,20];%select relevant channels: ttip 16, llip 18, jawY 20 
        sigsIn=nan(size(dataIn.data,1),length(artDimN));% build matrix to store signals
        newPHIs=nan(size(dataIn.data,1),length(artDimN));% build matrix to store phase values
        filtereds=nan(size(dataIn.data,1),length(artDimN));% build matrix to store denoised signals
        filteredidx={};% initialize vector with the indexes of the deterministic components as empty
        figure
        for nDim=1:length(artDimN)
            sigsIn(:,nDim)=dataIn.data(:,artDimN(nDim));% get sensor data
            sr=dataIn.samplerate;% sampling rate
            sigInT=[1:size(sigsIn,1)]./sr; % time steps
            [filtereds(:,nDim),imf,imfF,filteredidx{nDim},noiseStd]=mEMDdenoise(sigsIn(:,nDim),sr,nMasks,2,0.05,nRandNoiseRep,m,n);

            [PHI,~,~,~,~]=getPHImask(filtereds(:,nDim),sr,m,n,nMasks,ampCoeff,phaseMethod,threshs);
            newPHIs(:,nDim)=PHI;
            
            a(2*(nDim-1)+1)=subplot(6,1,2*(nDim-1)+1);
            plot(sigInT,sigsIn(:,nDim),'k');
            a(2*(nDim-1)+2)=subplot(6,1,2*(nDim-1)+2);
            plot(sigInT,newPHIs(:,nDim),'k');

        end
        xlabel('Time (sec)');

        linkaxes(a,'x')
    
    case 4 % Knee flexion data
        
        dataIn=load('./data./kneemovs.mat');%load data
        sigIn=dataIn.kneemovs([2480:19450]);% extractrelevant portion
        sr=dataIn.sr;% smaple rate
        sigInT=[1:length(sigIn)]./sr; % build signal's time steps
        cutOff=7;% cut off frequency as in original study
        filtLen=1000;
        lpfilt =fir1(filtLen, cutOff/(sr/2));%low pass filter coefficients
        sigInf=filter(lpfilt,1,sigIn); % filter input
        shift=floor(filtLen/2);
        sigInf=[sigInf(shift+1:end);ones(shift,1).*1e-7];
        sigInTf=sigInT(1:length(sigInf));% remove exceding time steps
        [PHI,~,~,~,~]=getPHImask(sigInf,sr/2,m,n,nMasks,ampCoeff,phaseMethod,threshs);

        options.MAXITERATIONS=10; % set maximum sifting iteration to 10
        Decomp=emd(sigInf,options)';% apply EMD
        normSig=normalize_cycle_amp(Decomp(:,1));% %refined amplitude normalizatin algo 

        figure
        c(1)=subplot(411);
        plot(sigInTf,sigInf,'k')% plot filtered signal
        
        ylim([min(-findpeaks(-sigIn))-1,max(findpeaks(sigIn))+1])% adjust axis y limits
        c(2)=subplot(412);
        plot(sigInTf,wrapTo2Pi(unwrap(angle(hilbert(zscore(sigIn))))),'k')%plot hilbert phase of input signal
        setPhaseAx(gca)
        c(3)=subplot(413);
        plot(sigInTf,wrapTo2Pi(unwrap(angle(hilbert(normSig)))),'k')%plot hilbert phase of first IMF after amplitude normalization
        setPhaseAx(gca)
        c(4)=subplot(414);
        plot(sigInTf,wrapTo2Pi(unwrap(PHI)),'k')%plot hilbert phase from our approach
        setPhaseAx(gca)

        linkaxes(c,'x')
        xlim([0,7000./sr])% set x axis limits
        xlabel('Time (sec)')

        % test the effect of the low pass filter frequency on the number of
        % cycles observed via EMD and via our approach
        niter=0;
        nCyclesEMD=nan(10,1); %initialize matrix containing number of cycles observed via EMD
        nCycles=nan(10,1);%initialize matrix containing number of cycles observed via our approach with inclusive behavior
        nCycles0=nan(10,1);%initialize matrix containing number of cycles observed via our approach with exclusive behavior
        filtFreqs=linspace(7,120,10);% low-pass cut-off frequencies to be tested
        
        for cutoff = round(filtFreqs) % for each cut-off freq.
            niter=niter+1;
            lpfilt =fir1(filtLen, cutoff/(sr/2)); % get filter coefficients
            sigInf=filter(lpfilt,1,sigIn);    % filter input signal
            shift=floor(filtLen/2);
             sigInf=[sigInf(shift+1:end);ones(shift,1).*1e-7];
            [PHI,~,~,~,~]=getPHImask(sigInf,sr/2,m,n,nMasks,ampCoeff,phaseMethod,threshs);
            
            Decomp=emd(sigInf)'; % apply EMD
            normSig=normalize_cycle_amp(Decomp(:,1));% apply refined amplitude normalization
            emdPhase=angle(hilbert(normSig)); % get Hilbert phase
            nCyclesEMD(niter)=sum(diff(unwrap(emdPhase))./(2*pi)); % count number of observed cycles via EMD
            nCycles(niter)=sum(diff(unwrap(PHI))./(2*pi)); % count number of observed cycles with inclusive behaviour

        end
        
        figure;
        plot(filtFreqs,nCycles,'k')% num of cycles observed with inclusive behavior
        hold on
        plot(filtFreqs,nCyclesEMD,'k:','linewidth',1);% num of cycles observed with EMD
        xlim([filtFreqs(1),filtFreqs(end)]);% set x axis limits
        xticks(round(filtFreqs));%set x axis ticks
        ylabel('Number of cycles')
        xlabel('Low pass filter cut-off frequency (Hz.)')
        
	case 5 %EGG data
        
        sr=5000;% set new signal sr 
        pathIn='./data/eggSig.wav';
        [audioIn,oldSr]=audioread(pathIn);% load data
        mySig=resample(audioIn(:,2),sr,oldSr);% resample at new sr
        mySigT=[1:length(mySig)]./sr; % build time samples vector
                       
        options.MAXITERATIONS=10;% set maximum num of sifting iterations
        [imf]=emd(mySig,options);% apply EMD
        options.MAXITERATIONS=1;% set maximum number of iterations 
        [imf1]=emd(mySig,options);% apply centring (sifing with one iteration)
        [PHI,newIMF,PHI0,centSig,mask]=getPHImask(mySig,5000,m,n,nMasks,ampCoeff,phaseMethod,threshs);
        
        options.MAXITERATIONS=1;% set maximum number of sifting iterations
        partialSift=emd(mySig+mask,options);%apply centering algo (sifting with one iteration) to signal plus mask
        partialSift=partialSift(1,:)'-mask;% remove mask

        mySigs=[mySig,imf1(1,:)',mask,mySig+mask,partialSift,newIMF];%build matrix to be plitted via stackedplot
        figure;
        %plot signals: input signal, centered signal (first IMF of EMD with one
        %sifting iteration); mask, input signal plus mask; partial IMF
        %obtained via masked sifting; final IMF obtained averaging
        %partial IMFs
        [bb,bbpos]=tight_subplot(6,1,0.03,0.08); %initialize subplots

        axes(bb(1))
        plot(mySig,'k');
        title('\textbf{Electroglottographic signal}','interpreter','latex')
        axes(bb(2))
        plot(imf1(1,:),'k');
        title('\textbf{Centred signal}','interpreter','latex')
        axes(bb(3))
        plot(mask,'k');
        title('\textbf{Mask signal}','interpreter','latex')
        axes(bb(4))
        plot(mySig+mask,'k');
        title('\textbf{Mask signal plus Electroglottographic signal}','interpreter','latex')
        axes(bb(5))
        plot(partialSift,'k');
        title('\textbf{Partial Sifting output}','interpreter','latex')
        axes(bb(6))
        plot(newIMF,'k');
        title('\textbf{Sifting output}','interpreter','latex')
        for nn=1:length(bb)
           if nn<length(bb)
             set(bb(nn),'xtick',[]) 
           end
           set(bb(nn),'box','off')
        end
        linkaxes(bb,'x')
        xlabel('Time (sec)')
        normSig=normalize_cycle_amp(imf(2,:)',[],[],5);%Refined normalization applied to second IMF from EMD
        PHI1=angle(hilbert(normSig)); % hilbert phase estimation 

        figure;
        [aa,aapos]=tight_subplot(3,1,0.05,0.08); %initialize subplots
        axes(aa(1));
        plot(mySigT,mySig,'k'); %plot input signal
        set(gca,'xtick',[])
        title('\textbf{Electroglottographic signal}','interpreter','latex','fontsize',14)
        axes(aa(2))
        plot(mySigT,PHI,'k');% plot phase from our approach
        set(gca,'xtick',[])        
        title('\textbf{Hilbert phase via the proposed method}','interpreter','latex','fontsize',14)
        axes(aa(3))
        plotMe=wrapTo2Pi(unwrap(PHI1));
        plot(mySigT,plotMe,'k');% plot phase from the approach in Huang et al. (2009)
        title('\textbf{Hilbert phase via the method in [5]}','interpreter','latex','fontsize',14)
        linkaxes(aa,'x')
        xlabel('Time (sec)')
end
```
