## Supplementary material for "Instantaneous phase of rhythmic behaviour under volitional control": Code and data files: paper_example_application.ipynb

```
# %% [markdown]
# Phase analysis of behaviuoral data
<p style='text-align: right;'> 
Leonardo Lancia</br>
Laboratoire Parole et Langage</br>
CNRS / Aix Marseille Univesité</br>
 

# %% [code]
import warnings
import numpy as np
from matplotlib import pylab, pyplot as plt
from scipy.io import loadmat, wavfile
from getPhaseToolsM import  getPhaseMask, mEMDdenoise
from scipy.stats import zscore
from scipy import signal

# %% [markdown]
We can now load our signal and plot it

# %% [code]

pathIn='./data/cl_pata_2.mat';
dataIn=loadmat(pathIn)
sigIn=dataIn['data'][:,15]
sr=dataIn['samplerate'][0][0]
plt.figure()
plt.plot(sigIn);


# %% [markdown]
## Phase estimation

# %% [markdown]
We set up a filter and apply it to our signal (and visualize the results)

# %% [code]
hiCut=20; #Max allowed frequency
filtLen=1000;#filter order

hiCutNorm=hiCut/( sr/2 );
b1=signal.firwin(filtLen,hiCutNorm,pass_zero='lowpass');

filtSig=signal.lfilter(b1,1,sigIn);
sigInf=filtSig[np.floor(filtLen/2).astype(int):];
plt.figure
plt.plot(sigInf);

# %% [markdown]
We then set the optional parameters of our phase extraction funcion

**phaseMethod** is a string or a cell of two strings (either 'h' or 'q') indicating the method to be used for the extraction of the quadrature signal. 'h' stands for Hilbert and 'q' for direct quadrature. If two strings are provided two different methods will be used in the two computations of the phase  (the first one is that aimed at estimating the frequency to be used in masked sifting ).

**thresh** is either a real (or a two element vector of reals) indicating the thershold used int the normalization of the cycles' amplitudes before the computation of the phase. Even in this case two different threshold are used in the two computations of the phase if two values are provided.

**m**, and **n** are the length and the order of the Savitsky Golay differentiator.

**nMasks** indicates the number of mask signals used in Masked Sifting (default val.: 22)

**ampCoeff** determines the amplitude of the masks as a proportion of  where r is the signal submitted to masked sifting. The value  represents a rough approximation of the range of a normal distribution given its standard deviation.

# %% [code]
phaseMethod='h'
thresh=5e-10
m=16
n=5
nMasks=22
ampCoeff=2

# %% [markdown]
The algorithm can now be launched and the phase plotted. The first output is the final estimation of the phase, the second output is the signal obtained from masked sifting; the third output is the phase estimated from the centered signal in the first part of the algorithm; the fourth output is the centered signal used to estimate phase and frequency in the first part and the fifth output is the masking signal used in masked sifting.

# %% [code]
PHI,imfout,phi0,centredS,mask0=getPhaseMask(sigInf,sr,16,5,22,2, ['h','h'], [1e-10,1e-10])
plt.figure()
ax1 = plt.subplot(211)
ax1.plot(sigIn);
ax2 = plt.subplot(212)
ax2.plot(PHI);

# %% [markdown]
## Denoising algorithm

We may want to process our data in a more adaptive way, which is without setting a low-pass cut-off frequency for the filter. In this case we can use the function  mEMDdeNoise

For that purpose, we may need to define the value of optional the parameter **nRandNoiseRep** determining the number of simulated random processes, and that of the optional parameter **alphaVal**, determining the significance threshold.

The first output of the function represents the filtered signal, the second output represents the IMFs, the fourth output corresponds to their their frequencies, the fifth output represents the indexes of the IMFs that added produce the filtered signal and the last output represents the standard deviation of the estimated random component.

# %% [code]
nRandNoiseRep=100;
alphaVal=0.05;
#apply denoising

filtered, imf, imfF, filteredidxs, noiseStd=mEMDdenoise(sigIn,sr,nMasks=22,ampCoeff=2,alpha=0.05,nReps=100,m=16,n=5)
plt.figure()
ax1 = plt.subplot(211)
ax1.plot(sigIn);
ax2 = plt.subplot(212)
ax2.plot(filtered);

# %% [markdown]
We can again estimate the phase of the clean signal and plot the results

# %% [code]
PHI,imfout,phi0,centredS,mask0=getPhaseMask(filtered,sr,16,5,22,2, ['h','h'], [1e-10,1e-10])
plt.figure()
ax1 = plt.subplot(211)
ax1.plot(sigIn);
ax2 = plt.subplot(212)
ax2.plot(PHI);
```
