## Supplementary material for "Instantaneous phase of rhythmic behaviour under volitional control": Code and data files: paper_illustrations.m

```
% Script to reproduce the illustration of the paper
% sections should be independent one from the other
%
% WARNING if it has to be distributed IT NEEDS TO BE CLEANED CHECKED AND COMMENTED 

% Leonardo Lancia 14/10/2023
% mailto:

clear
warning 'off'

addpath('./gen_utils');
addpath(genpath('./get_phase')); 
%% problems solvable with Huang's approach.


% addpath(['.\package_emd\EMDs']);
sr=1000;
dur=0.8;
t=[1:dur*sr]./sr;
omg=12;
phaseY=t.*omg.*2*pi;
y=sin(phaseY);
phaseY1=t*(omg/1.75)*2*pi;
y1=0.8.*sin(phaseY1);
phaseY2=t*(omg/32)*2*pi;
y2=2.*sin(phaseY2);

figure;
subplot(4,2,[1,3,5,7])
ss=stackedplot([y;y1;y2]','DisplayLabels',{'x_{1}','x_{2}','x_{3}'},'color','k');
for i=1:length(ss.DisplayLabels)
    axesProps = struct(ss.AxesProperties(i));
    axesProps.Axes.YLabel.Interpreter = 'tex';
end
xlabel('Time steps')
aa(1)=subplot(4,2,2);
plot(y+y1,'k');
set(gca,'xtick',[])
title('x_{1}+x_{2}','Interpreter','tex')
aa(2)=subplot(4,2,4);
plot(wrapTo2Pi(unwrap(angle(hilbert(y+y1)))),'k');
set(gca,'xtick',[])
title('\phi(x_{1}+x_{2})','Interpreter','tex')

aa(3)=subplot(4,2,6);
plot(y+y1+y2,'k');
set(gca,'xtick',[])
title('x_{1}+x_{2}+x_{3}','Interpreter','tex')
aa(4)=subplot(4,2,8);
plot(wrapTo2Pi(unwrap(angle(hilbert(y+y1+y2)))),'k');
title('\phi(x_{1}+x_{2}+x_{3})','Interpreter','tex')
xlabel('Time steps')
linkaxes(aa,'x');

[maxEnv,minEnv,meanEnv]=get_envelopes([y+y1+y2]');
myIMF=(y+y1+y2)-meanEnv;

figure;
[ha, pos] = tight_subplot(3,2,[0.02,0.05],0.12);

txtY=3.5;
txtX=25;
fSize=12;

axes(ha(1));
plot(y+y1+y2,'k')
set(gca,'xtick',[])
hold on 
plot(maxEnv,'k','linestyle',':','linewidth',1)
plot(minEnv,'k','linestyle',':','linewidth',1)
plot(meanEnv,'k','linestyle','-.','linewidth',1)
line(get(gca,'xlim'),[0,0],'color','k')
set(gca,'xtick',[])
text(txtX,txtY,'(a)','FontSize',fSize);
axes(ha(3));
plot(myIMF,'k')
xlabel('Time steps')
line(get(gca,'xlim'),[0,0],'color','k')
txtY=1.5;
text(txtX,txtY,'(b)','FontSize',fSize);

axes(ha(4));
plot(abs(myIMF),'k')
set(gca,'xtick',[])
hold on;
[~,pksLoc,proms]=findpeaks(abs(myIMF),'MinPeakProminence',0);
pksVal=abs(myIMF(pksLoc));
[pksVal,pksLoc]=SetBoundCond(pksVal',pksLoc',length(myIMF));
newIdxs=[1:length(myIMF)]';
refSig= interp1(pksLoc,pksVal,newIdxs, 'pchip');
plot(refSig,'k:','linewidth',1)
ylim([0,2])
txtY=1.8;
text(txtX,txtY,'(c)','FontSize',fSize);

axes(ha(6));
plot(normalize_cycle_amp([(y+y1+y2)-meanEnv]'),'k')
ylim([-1.8,1.8])
txtY=1.5;
text(txtX,txtY,'(d)','FontSize',fSize);

xlabel('Time steps')
delete(ha([2,5]))

Decomp=emd([y+y1+y2]');
normSig=normalize_cycle_amp(Decomp(1,:)');%Huang normalizatin algo

figure;
subplot_tight(4, 2, 1, [0.07, 0.08]);
plot(y+y1+y2,'k')
set(gca,'xtick',[])
title('\textbf{Input signal}','Interpreter','latex')
subplot_tight(4, 2, [3,5,7], [0.07, 0.08]);
ss=stackedplot([Decomp]','DisplayLabels',{'x_{1}','x_{2}','x_{3}'},'Color','k');
for i=1:length(ss.DisplayLabels)
    axesProps = struct(ss.AxesProperties(i));
    axesProps.Axes.YLabel.Interpreter = 'tex';
end

xlabel('Time steps')

subplot_tight(4, 2, 4, [0.07, 0.05]);

plot(normSig,'k');
set(gca,'xtick',[])
title('\textbf{N($$IMF_{1}$$)}','Interpreter','latex')
subplot_tight(4,2,6, [0.07, 0.05]);
plot(wrapTo2Pi(unwrap(angle(hilbert(normSig)))),'k')
xlabel('Time steps')
setPhaseAx(gca)
title('\textbf{ $$\phi(x_{1}+x_{2}+x_{3})$$ }','Interpreter','latex')

%% problems we can solve
addpath(genpath(['.\utils'])); 

sr=1000;
dur=1.5;
t=[1:dur*sr]./sr;
omg=12;
phaseY=t.*omg.*2*pi;
y=sin(phaseY);
phaseY1=t*(omg/1.75)*2*pi;
y1=0.8.*sin(phaseY1);
phaseY2=t*(omg/32)*2*pi;
y2=2.*sin(phaseY2);
phaseYOO=t.*3*omg.*2*pi.*(square(t*(omg/6)*2*pi)>0);
yOnOff=sin(t.*3*omg.*2*pi).*(square(t*(omg/6)*2*pi)>0);

threshs1=[NaN,NaN];threshs2=[NaN,NaN];
m = 16;      % number of filtered points (FIR order)
M = (m+1)/2;      % output point , M = (m+1)/2; % middle point for odd m
n = 5;      % approximation polynomial order
handleNegPHI='interp'; % set strategy to adopt in intervals with negative frequency
phaseMethod={'h','h'};

inclusiveRef=[0,0];

nMasks=16;

YY=y+y1+y2+0.2*yOnOff;

Decomp=emd(YY)';
normSig=normalize_cycle_amp(Decomp(:,1));%Huang normalizatin algo
normSig1=normalize_cycle_amp(Decomp(:,2));%Huang normalizatin algo
figure;
[ha, pos] = tight_subplot(6,3,[0.04,0.05],0.08);
plotNums=[1,4,7,10,13];

plotData=[YY;yOnOff;y;y1;y2]';
titles={'\textbf{Input signal}',...
    '\textbf{Intermittent component}',...
    '\textbf{$$x_{1}$$}',...
    '\textbf{$$x_{2}$$}',...
    '\textbf{$$x_{3}$$}'};
for i = 1:length(plotNums)
    axes(ha(plotNums(i)));
    plot(plotData(:,i),'k');
    if i< length(plotNums)
        set(gca,'xtick',[])
    else
        xlabel('Time steps')
    end
    title(titles{i},'Interpreter','latex')
end
titles={'\textbf{Input signal}',...
    '\textbf{$$IMF_{1}$$}',...
    '\textbf{$$IMF_{2}$$}',...
    '\textbf{$$IMF_{3}$$}',...
    '\textbf{$$IMF_{4}$$}',...
    '\textbf{$$IMF_{5}$$}'};
i0=i;
plotNums=[2,5,8,11,14,17];
plotData=[YY',Decomp];
for i = 1:(min(size(plotData,2),length(plotNums)))
    axes(ha(plotNums(i)));
    plot(plotData(:,i),'k');
    if i< length(plotNums)
        set(gca,'xtick',[])
    else
        xlabel('Time steps')
    end
	title(titles{i},'Interpreter','latex')
end
i0=i+i0;
plotNums=[3,6,9,12,15,18];
plotData=[wrapTo2Pi(phaseY)',Decomp];
titles={'\textbf{$$\phi(x_{1})$$}',...
    '\textbf{$$\phi(IMF_{1})$$}',...
    '\textbf{$$\phi(IMF_{2})$$}',...
    '\textbf{$$\phi(IMF_{3})$$}',...
    '\textbf{$$\phi(IMF_{4})$$}',...
    '\textbf{$$\phi(IMF_{5})$$}'};
for i = 1:min(size(plotData,2),length(plotNums))
    axes(ha(plotNums(i)));
    if i==1
        plot(plotData(:,i),'k');
    else
        plot(wrapTo2Pi(unwrap(angle(hilbert(normalize_cycle_amp(plotData(:,i)))))),'k');
    end
    if i< length(plotNums)
        set(gca,'xtick',[])
    else
        xlabel('Time steps')
    end
    title(titles{i},'Interpreter','latex')
    setPhaseAx(gca);
end
delete(ha(16))
% 
[PHI,siftedSig,PHI0,ceteredSig,mask]=getPHImask(YY,sr,m,n,nMasks,[],phaseMethod,threshs1);

figure;
[hb, pos] = tight_subplot(4,1,[0.04,0.05],0.1);
axes(hb(1));
plot(YY,'k')
set(gca,'xtick',[])
title('\textbf{Input signal}','Interpreter','latex')
normSig=normalize_env_peaks(Decomp(:,1));%Huang normalizatin algo
axes(hb(2));

plot(PHI0,'k');
set(gca,'xtick',[])
setPhaseAx(hb(2))
title('\textbf{Initial phase estimate}','Interpreter', 'latex')

axes(hb(3));
plot(PHI,'k')
setPhaseAx(hb(3))
title('\textbf{Final phase estimate}','Interpreter', 'latex')

set(gca,'xtick',[])
axes(hb(4))
plot((wrapTo2Pi(angle(hilbert(normSig)))'),'k');
setPhaseAx(hb(4))
title('\textbf{Phase estimate through EHHT}','Interpreter', 'latex')

xlabel('Time steps')


%% problems we can solve 2

sr=1000;
dur=1.5;
t=[1:dur*sr]./sr;
omg=12;
phaseY=t.*omg.*2*pi;
y=sin(phaseY);
phaseY1=t*(omg/1.75)*2*pi;
y1=0.8.*sin(phaseY1);
phaseY2=t*(omg/32)*2*pi;
y2=2.*sin(phaseY2);
phaseYOO=t.*3*omg.*2*pi.*(square(t*(omg/6)*2*pi)>0);
omg1=(sin(t.*2*pi*omg*1.5+pi)+1.2);
omg1=omg.*omg1;
y=cos(2*pi*cumsum(omg1./sr));

yOnOff=sin(2*pi*cumsum(omg1./sr)).*(square(t*(omg/6)*2*pi)>0);
YY=y+y1+y2+0.2*yOnOff;

options.MAXITERATIONS=10;
Decomp=emd(YY,options)';

normSig=normalize_env_peaks(Decomp(:,1));

[pksVals, pksLocs]=findpeaks(normSig(:,1));
[valVals, valsLocs]=findpeaks(-normSig(:,1));
pksLocs=pksLocs(pksVals<0);
valsLocs=valsLocs(valVals<0);
centers=[pksLocs;valsLocs];

radius=0.17;
yradius = radius;
% Create a circle "template" with a trailing NaN to disconnect consecutive circles
t = linspace(0, 2*pi, 100);
t(end+1) = NaN;

figure;
[ha, pos] = tight_subplot(2,1,[0.02,0.05],0.12);

axes(ha(1));
plot(Decomp(:,1),'k')
set(gca,'xtick',[])
axes(ha(2));
plot(normSig,'k');
aspectRatio=daspect;
xradius = 0.1*radius * aspectRatio(1)/aspectRatio(2);
circle = [xradius*cos(t(:)), yradius*sin(t(:))];

for nn =1:length(centers)
    xx=centers(nn);
    yy=normSig(centers(nn));
    circles = arrayfun(@(xx,yy)bsxfun(@plus, [xx,yy], circle), xx, yy, 'uni', 0);
    circles = cat(1, circles{:});
    hold on
    plot(circles(:,1), circles(:,2), 'color', 'r')
end

linkaxes(ha,'x')
xlabel('time steps')
xlim([700,1250]);


resid1=Decomp(:,1);
resid2=Decomp(:,1);

totNiters=5;
figure;
[ha, pos] = tight_subplot(totNiters,2,[0.03,0.05],0.12);

nSubIter=0;
for nIter=1:totNiters
    [extr,extrLoc]=findpeaks(abs(resid1));
    [extr,extrLoc]=SetBoundCond(extr,extrLoc,length(resid1));
    
    newIdxs1=1:length(resid1);
    newIdxs2=1:length(resid2);
    [minV,minLoc]=findpeaks(-resid2);
    [maxV,maxLoc]=findpeaks(resid2);
    [minLoc,maxLoc,minV,maxV] = boundary_conditions(minLoc',maxLoc',newIdxs2,resid2',resid2',2);
    [extrLoc2,extrOrd]=sort([minLoc,maxLoc]);
    extr2=[abs(minV),abs(maxV)];
    extr2=extr2(extrOrd);
    
    env1=interp1(extrLoc,extr,newIdxs1,'pchip');
    env2=interp1(extrLoc2,extr2,newIdxs2,'pchip');
    
    nSubIter=nSubIter+1;
    axes(ha(nSubIter));

    plot(resid1,'k');
    hold on;
    plot(abs(resid1),'r:','linewidth',1)
    plot(env1,'b--');
    if nSubIter<totNiters*2-1
        set(gca,'xtick',[])
    else
        xlabel('Time steps')
    end
    if nSubIter>1
        ylim([-1.2,1.2])
    end
    if nIter==1
        title('\textbf{Amplitude Normalization Iterations}','interpreter','latex')
    end
    nSubIter=nSubIter+1;
    axes(ha(nSubIter));
    plot(resid2,'k')
    hold on;
    plot(abs(resid2),'r:','linewidth',1)
    plot(env2,'b--');
    if nSubIter<totNiters*2-1
        set(gca,'xtick',[])
    else
        xlabel('Time steps')
    end
	set(gca,'ytick',[])
	if nSubIter>2
        ylim([-1.2,1.2])
    end
    if nIter==1
        title('\textbf{Refined Amplitude Normalization Iterations}','interpreter','latex')
    end
    resid1=resid1./env1';
    resid2=resid2./env2';

end
linkaxes(ha,'x')
xlim([1025,1215])
```
