## Supplementary material for "Instantaneous phase of rhythmic behaviour under volitional control": Code and data files: readme.md

# Get Phase Tools
Code to extract the continuous instantaneous phase of experimental signals according to the paper: 

Instantaneous phase of rhythmic behaviour under volitional control  
Leonardo Lancia  
bioRxiv 2023.11.01.564135; doi: https://doi.org/10.1101/2023.11.01.564135

## Content:

**get_phase** folder and its content: Matlab scripts implementing the methods proposed.  
**gen_utils** folder: Matlab functions required to reproduce the analyses reported in the paper.  
**data** folder: files analysed in the paper and used to exemplify the method proposed there.  
**demo.mlx** file: Matlab Live Script demo file illustrating the functioning of the matlab functions.  
**paper_analyses.m** file: Matlab script to launch the analyses reported in the paper.  
**getPhaseToolsM.py** file: Python functions implementing the methods proposed.  
**paper_example_application.ipynb** file: Jupyter notebook demo file illustrating the functioning of the Python functions.
**requirements.txt** file: requirement file to install Python dependencies via PIP.  
**requirements.yml** file: requirement file to install Python dependencies via conda.  


# Installation Instructions:

## Matlab:

In order to use the proposed methods in you own workflow you need to add the folder **get_phase** and its subfolders to the Matlab path. This can be done by going to the location containing the folder **get_phase** and run:

addpath(genpath('./get_phase'))

This is not required when running demo.mlx from the location containing the folders **get_phase** and **data** or when running **paper_analyses.m** 
from the location containing the folders **get_phase**, **gen_utils** and **data**.  

Note: the provided programs have been developed with Matlab 2018b endowed with the Statistics and the Signal Processing toolboxes. 
In order to make the programs work without these tolbxes, the present dirstibution includes several relacement functions 
(mainly gathered from the freely avialble Fieldtrip toolbox [1], see: https://www.fieldtriptoolbox.org/faq/matlab_replacements/). 
These functions are stored in the subflders 'get_phase\utils\tbxsSubst\signalProcessing' and 'get_phase\utils\tbxsSubst\statistics'. 
If however the required toolboxes are avialable, it advised to delete the corresponding subfolders.

## Python:  
To install the modules required for the functioning of the methods proposed in the paper
go to the folder containitng the files **requirements.txt** and **requirements.yml**

here, depending on the package manager of your choice, type: 

pip install -r requirements.txt

or

conda env update --name [envname] --file requirements.yml

where **[envname]** should be substituted by the name of the environment where you want to use Get Phase Tools.

# Usage: 
The scripts are accessible via two main functions having the same names and arguments in both Matlab and Python implementations: 
**getPhaseMask** and **mEMDdenoise**. Their functioning is illustrated by the two demo files **paper_example_application.mlx** (Matlab Live Editor)
and **paper_example_application.ipynb** (Python Jupyter notebook).

## getPhaseMask
This function computes the instantaneous phase of an input signal given its sampling frequency. Its first argument is the input signal and its second argument the sampling frequency. The following optional arguments are:

**m** (positive integer; optional, default: 16): number of values considered in the application of Savitzky-Golay differentiator.  
**n** (positive integer; optional, default: 5): polynomial order of the differentiator.  
**nMasks** (positive integer; optional, default 22): number of masking signals used to extract the independent mode function via masked sifting.  
**ampCoeff** (positive real; optional, default: 2): coefficient determining the amplitude of the masks as a proportion of 4*std(signal) (as this is a rough estimate of the signal's range from the std, if the signal's values are normally distributed).  
**quadMethod** (a string or a cell of two strings in Matlab and a string or a list of two strings in Python, default: {'h','h'} in Matlab and ['h','h'] in Python): Method to be use in the computation of the quadrature signal. 'h' stands for Hilbert and 'q'. If two strings are provided a different method will be adopted in in the first or the second part of the algorithm.  
**threshs** (scalar or vector of two positive real values close to zero, default: [1E-10,1E-10]  ): threshold for refined amplitude normalization. If two values, different thresholds will be used in the two parts of the algorithm.  

The function's outputs are:  
**PHI**: the signal's phase.  
**IMF1**: output of masked sifting.  
**PHI**: initial phase estimte.  
**centredSig**: centred signal.  
**mask**: masking signal with inital phase set to 0


## mEMDdeNoise
The function removes high frequency noise from an input signal given its sampling frequency. Its first argument is the input signal and its second argument the sampling frequency. The other (optional) arguments are:  

**nMasks** (positive integer; optional, default: 22): number of masking signals for masked EMD application.  
**ampCoeff** (positive real; optional, default: 2*estimated range of first IMF): amplitude of the masking signals.  
**nReps** (positive integer; optional, default: 100): number of repetitions of simulated random processes.  
**m** (positive integer; optional, default: 16): number of values considered in the application of the Savitzky-Golay differentiator.  
**n** (positive integer; optional, default: 5): order of the Savitzky-Golay differentiator.

the function's outputs are:  

**filtered**: filtered signal.
**imf**: all signal's IMFs.  
**imfF**: frequencies of extracted IMFs.  
**filteredidx**: indexes of IMFs composing the filtered signal.  
**noiseStd**: standard deviation of the estimated noise (sum of the random components).

# Copyright

This code is licensed under the GNU-GPL (see LICENSE file). If you use it, you have to refer to the companion paper avialable here: https://doi.org/10.1101/2023.11.01.564135  

# References  

[1] Robert Oostenveld, Pascal Fries, Eric Maris, and Jan-Mathijs Schoffelen. FieldTrip: Open Source Software for Advanced Analysis of MEG, EEG, and Invasive Electrophysiological Data. Computational Intelligence and Neuroscience, 2011; 2011:156869
